# Distinct substrate and intermediate recognition via mutation effects on *Mycobacterium tuberculosis* methionyl-tRNA synthetase

**DOI:** 10.1101/2025.06.06.658285

**Authors:** Shivani Thakur, Rukmankesh Mehra

## Abstract

Tuberculosis kills millions worldwide. Drug-resistance demands exploring new targets against this illness. Methionyl-tRNA synthetase (MetRS) is a crucial target in *Mycobacterium tuberculosis* (*Mtb*) that participates in initiation and elongation of translation and represents a protein of evolutionary interest. To elucidate the structure-function relationships of MetRS, we performed detailed sequence analyses and molecular dynamics simulations of *Mtb* MetRS in the substrate-bound (methionine and ATP) and intermediate (methionyl-AMP) states, for both the wild-type and three single-mutant forms (H21A, K54A, and E130A). Eight systems (two wild-type and six mutants) were simulated for 24 microseconds. Differential dynamics and binding effects of the substrate versus intermediate states were identified, along with the molecular reasons for the loss of activity in mutants. The wild-type substrate state was more stable than the intermediate state. In contrast, the mutants were more unstable in the substrate state, but incorporated stability into the intermediate state protein. These findings suggest that methionyl-AMP, being a reaction intermediate, exhibits a short residence time at the protein’s active site, while the substrate state shows a longer residence time of methionine and ATP. The increased instability of mutants in the substrate state indicates disruption of the pyrophosphate-ATP exchange by altering substrate-protein interactions. Once the intermediate is formed, the mutations have minimal or no effect. These observations are consistent with experimental data. In brief, our study finds the molecular basis for the distinct substrate and intermediate recognition by *Mtb* MetRS and establishes a mechanism for loss of activity in the mutants.

## 1 Introduction

Tuberculosis (TB), caused by *Mycobacterium tuberculosis* (*Mtb*), affects about a quarter of the world’s population [1]. The Global Tuberculosis Report (2024) revealed that TB is likely to become the world’s leading cause of death from a single infectious agent, surpassing COVID-19 [1]. Current TB therapies face challenges due to multidrug resistance, extensive drug resistance, and the latent bacterial forms [2]. As a result, continuous efforts have been made to identify new antitubercular targets [3,4,13,14,5–12]. Aminoacyl tRNA synthetase (aaRS) enzymes represent one such promising class of targets [15–17]. These enzymes play an essential role in translating the genetic code into protein sequence by attaching amino acids to their cognate tRNAs [18,19]. This function makes aaRS an irreplaceable component for protein synthesis and an attractive bacterial target for investigation [15–17].

Methionyl-tRNA synthetase (MetRS) charges tRNA with methionine (Met) and participates in both the initiation and elongation phases of protein synthesis [20]. This key role makes MetRS a target of interest for antimicrobial drug development [21,22,31–40,23–30]. *Mtb* MetRS consists of 519 amino acid residues (**Figure 1a**). It differs significantly from its human counterparts, sharing only ∼16.9% and ∼34.9% sequence identity with the human cytosolic and mitochondrial MetRS, respectively [20,41,42].

**Figure 1.**
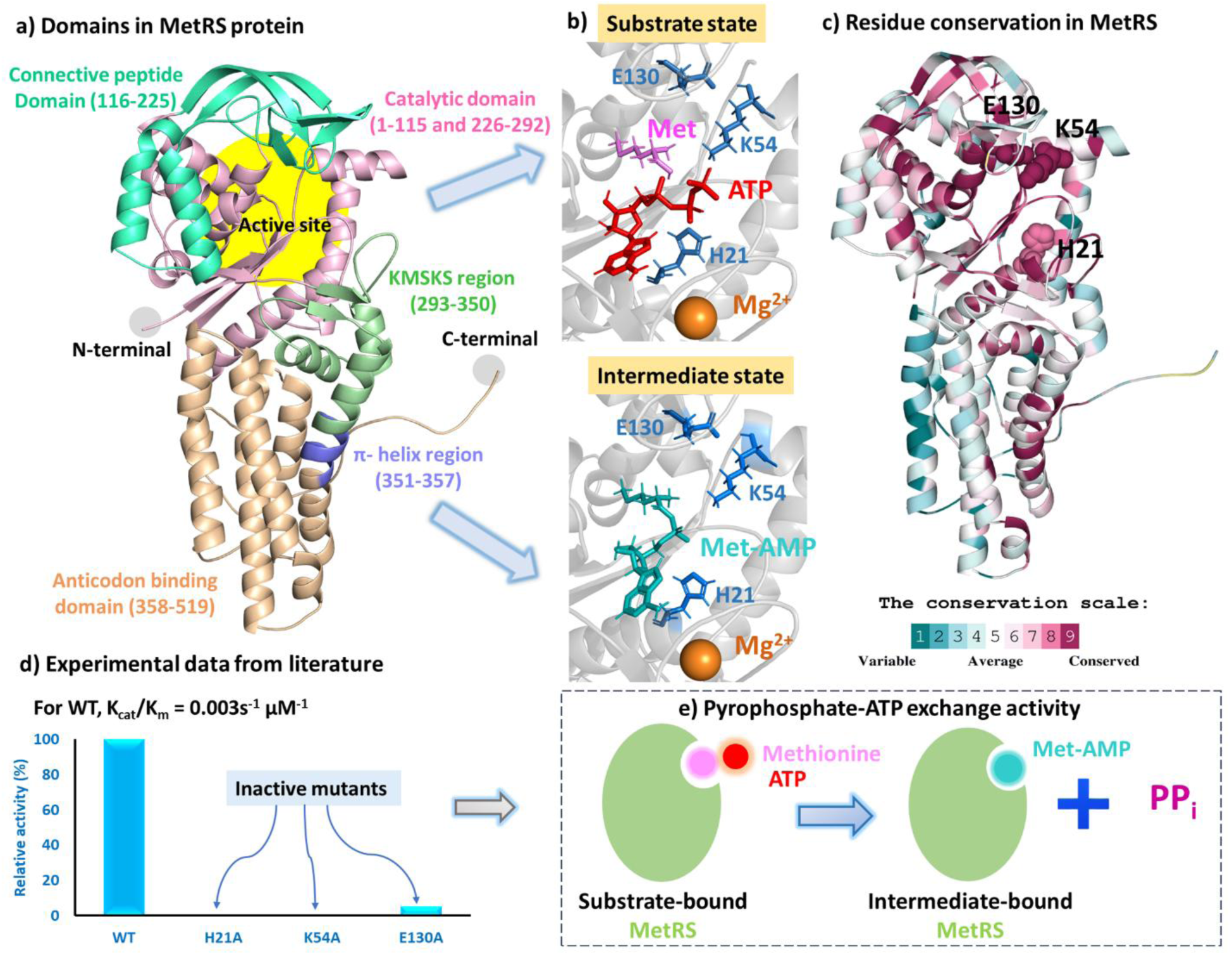
Structure and activity of *Mycobacterium tuberculosis* MetRS. **(a)** *Mtb* MetRS structure highlighting the protein domains and the active site (yellow). **(b)** The substrate state contains MetRS bound to methionine (purple) and ATP (red), while the intermediate state contains MetRS bound to Met-AMP (cyan). Binding site residues, displayed in dark blue sticks, were studied for alanine mutations in this work. **(c)** Protein residues colored based on evolutionary conservation. Dark maroon indicates a highly conserved regions (conservation score of 9) and dark cyan (score of 1) represents non-conserved parts. **(d)** Relative pyrophosphate-ATP exchange activity of wild-type (WT) and mutant (H21A, K54A, and E130A) MetRS, as reported in the literature [20]. All three mutants caused almost complete loss of activity. **(e)** Schematic representation of the pyrophosphate-ATP exchange activity, where methionine and ATP are converted to Met-AMP with the release of pyrophosphate (PP_i_) as a byproduct.

MetRS belongs to class I aaRS family, characterized by conserved signature motifs (_18_HVGH_21_ and _299_KMSKS_303_) and the presence of Rossmann folds in the catalytic domain [43]. Specifically, *Mtb* MetRS is classified as a MetRS1-type enzyme, distinguished by a single “knuckle motif” and the absence of a Zn^2+^ binding metal coordination site [20,41]. In contrast, class II aaRSs comprise unique α/β folds and three conserved signature motifs [44]. Structurally, *Mtb* MetRS can be divided into five regions, namely, catalytic domain (residues 1-115 and 226- 292), the connective peptide (CP) domain (116-225), the KMSKS region (293-350), the π-helix (351-357), and the anticodon binding domain (358-519) (**Figure 1a**). The active site is formed by contributions from the catalytic domain, CP domain and KMSKS regions, while the anticodon domain facilitates tRNA recognition and binding [20,41].

MetRS is known to catalyze a two-step reaction [41]. The substrates methionine and adenosine triphosphate (ATP) bind to the active site of the enzyme and are converted into the intermediate methionyl-AMP (Met-AMP), with pyrophosphate (PP_i_) released as a byproduct. The second step involves the binding of tRNA (either initiator or elongator) to MetRS [45]. A nucleophilic attack by the 3′ CCA end of the tRNA on Met-AMP facilitates the transfer of methionine to the tRNA. The intermediate-bound *Mtb* MetRS structure is available in the Protein Data Bank (PDB ID: 5XET) [20,46], whereas the substrate-bound state is absent. Consequently, the mechanism of substrate recognition in *Mtb* MetRS remains unclear.

Here, we aim to study the binding mechanism involved in the first step of substrate recognition and its intermediate formation. Three *Mtb* MetRS mutations, H21A, K54A, and E130A (**Figure 1b**), are known to disrupt its adenylation activity (**Figure 1d**), i.e., they affect the first step of the aminoacylation reaction (**Figure 1e**) [20]. His21 and Lys54 are part of an α-helix in the catalytic domain, whereas Glu130 is located in a turn region of the CP domain. His21 is present in _18_HVGH_21_ motif and stabilizes ATP phosphate chain during adenylation reaction [47]. Lys54 (in the catalytic domain) forms hydrogen bonding with CP domain residue Arg128, whereas Glu130 (in the CP domain) forms hydrogen bonding with catalytic domain residue Lys53, thus contributing to active site integrity and stability [20]. All three residues are highly conserved in MetRS1 category [48]. How do these mutations affect the substrate- and intermediate-bound states of MetRS? To understand these mechanisms, we performed evolutionary analysis using MetRS sequences and modeled 3D structures of the wild-type (WT) protein and its point mutants (H21A, K54A or E130A) in the substrate- and intermediate-bound states. Molecular dynamics (MD) simulations were carried out for total eight systems (2 WT and 6 mutants), for 24 microseconds (µs). This analysis revealed distinctive features between the substrate and intermediate states, as well as between WT and mutant complexes.

## 2 Methods

### 2.1 Multiple sequence alignment of MetRS

The quantitative analysis of *Mtb* MetRS sequence conservation was carried out using the CONSURF server [49]. It searches against the UNIREF90 cluster of UniProt [50] and performs multiple sequence alignment with up to 150 homologous sequences.

A phylogenetic tree of *Mtb* MetRS was built using iTOL v7 [51]. For this, multiple sequence alignment of *Mtb* MetRS with 25 other MetRS sequences was performed using Clustal Omega (**Table S1**) [52]. This alignment included representatives from *Mycobacterium* species, eukaryotes, archaea, and Gram-negative and Gram-positive bacteria.

### 2.2 Substrate and intermediate complexes

To model the protein states, we used the 3D structure of *Mtb* MetRS available in the Protein Data Bank (PDB ID: 5XET) [20]. The 5XET structure contains MetRS bound to the reaction intermediate Met-AMP and Mg^2+^ ions. However, the substrate bound structure was not available in the PDB. To model this state, the methionine coordinates were copied from the *Mycobacterium smegmatis* MetRS (PDB ID: 2X1L) [53] to the *Mtb* MetRS (5XET) structure. The two MetRS structures exhibit a structural root-mean-square deviation (RMSD) of 0.90 Å and share 75.54% sequence identity (based on BLAST analysis) [54]. ATP coordinates were obtained from the *Staphylococcus aureus* MetRS structure (PDB ID: 7WPN) [48], which shows a structural RMSD of 1.70 Å and sequence identity of 41.35% with *Mtb* MetRS. Since Mg^2+^ plays a role in both the activation and charging steps of aminoacylation, [55–59] one Mg^2+^ was retained at the active site of MetRS in both substrate and intermediate states.

### 2.3 Wild-type and mutant models

The starting models of the substrate and intermediate states were prepared using CHARMM-GUI [60]. Missing protein residues were added, and wild-type (WT) and mutant states were prepared. Three mutant models were generated for each state (substrate and intermediate), incorporating the point mutations H21A, K54A or E130A. These mutations are known to cause a significant loss of adenylation activity in *Mtb* MetRS (**Figure 1d**) [20]. Thus, a total of 8 systems were prepared: two WT and six mutants. All systems were prepared at pH 7 in TIP3P water, neutralized by counterions (Na^+^ or Cl^-^), and a salt concentration of 0.15 M was maintained.

### 2.4 Molecular dynamics simulations

The eight systems were energy minimized using the steepest descent integrator, [61] followed by a two-step equilibration process in the NVT and NPT ensembles for 100 picoseconds each, using leap-frog integrator [62]. Molecular dynamics (MD) simulations were performed for each system in the NPT ensemble for 1 microsecond (μs), in triplicate, with a time-step of 2 femtoseconds. In total, 24 μs simulations (8 systems × 1 μs × 3 runs) were performed. Simulations were carried our using GROMACS [63] and CHARMM36 [64] forcefield. 300 K temperature and 1 bar pressure were maintained using V-rescale thermostat and C-rescale barostat, respectively [65,66].

### 2.5 Analysis

The primary analysis was performed on complete 1 μs MD trajectories, including root mean square deviation (RMSD), radius of gyration (R_g_) and total solvent accessible surface area (SASA). The secondary analysis was conducted on the equilibrated final 500 ns trajectories, which included hydrophilic and hydrophobic SASA, root-mean-square-fluctuation (RMSF), intra- and inter- molecular hydrogen bonds, and the N- to C- terminal distance of the protein.

Free energy landscape principal component analysis (FEL-PCA) was performed on the final 500 ns runs to capture the fundamental protein motions. The covariance matrix was computed for backbone atoms and diagonalized to determine the characteristic eigenvectors (EVs) and their eigenvalues. We plotted the first twenty EVs along with their eigenvalues, and additionally visualized EV1 versus EV2 values to assess the conformational landscapes. The representative structure from each simulation was identified using Gromos clustering method with a backbone RMSD cut-off of 0.3 nanometer [63]. Protein’s secondary structure analysis was then performed on these representative structures.

## 3 Results and discussion

### 3.1 Sequence alignment revealed conservation of H21, K54 and E130 in MetRS

Sequence conservation of *Mtb* MetRS was quantitatively analyzed. **Figure 1c** displays the conservation score of residues mapped onto the MetRS structure, where higher values (maroon color) indicate greater conservation. The catalytic (residues 1-115 and 226-292) and CP (116-225) domain residues were highly conserved (dark maroon), whereas the anticodon domain was comparatively less conserved (blue). The three residues H21, K54 and E130 were highly conserved, with conservation scores of 8, 9, and 9, respectively (maroon). H21 is a part of the signature HIGH motif (18-21) of MetRS, which was also conserved. These residues are located within the protein’s active site: H21 and K54 in the catalytic domain, and E130 in the CP domain.

These results indicate that the residues surrounding the active site are highly preserved. Specifically, the high conservation of H21, K54 and E130 highlights their significant role in the aminoacylation reaction.

### 3.2 Phylogenetic tree placed *Mtb* MetRS close to eukaryotic mitochondrial MetRS

To identify the closest homologs of *Mtb* MetRS, a phylogenetic tree was built using 26 MetRS sequences from *Mycobacterium* species, eukaryotes, archaea, and Gram-negative and Gram-positive bacteria (**Figures S1** and **S2**). The *Mycobacterium* sequences clustered together, indicating high similarity among them. Eukaryotic cytoplasmic MetRS sequences were highly divergent from *Mtb* MetRS and were therefore placed distantly in the tree. In contrast, eukaryotic mitochondrial MetRS sequences were positioned adjacent to the mycobacterial.

This suggests that the mycobacterial MetRS sequences share close homology with eukaryotic mitochondrial MetRS, while the eukaryotic cytoplasmic MetRS sequences are distantly related to *Mtb* MetRS (**Figure S2**). This also supports the endosymbiotic origin of mitochondria [67,68].

### 3.3 Stability differences between WT and mutant proteins in substrate and intermediate states

Primary analysis was performed on the protein backbone atoms for the full 1 µs simulations. We made two comparisons: (i) substrate versus intermediate state, and (ii) WT versus mutant states. In WT systems, the protein RMSD was lower in the substrate state than in the intermediate state (**Figure 2a, b**), with the average RMSD (across three simulations) of 0.34 and 0.42 nm, respectively (**Figure 3a** and **Table S2-S3**).

**Figure 2.**
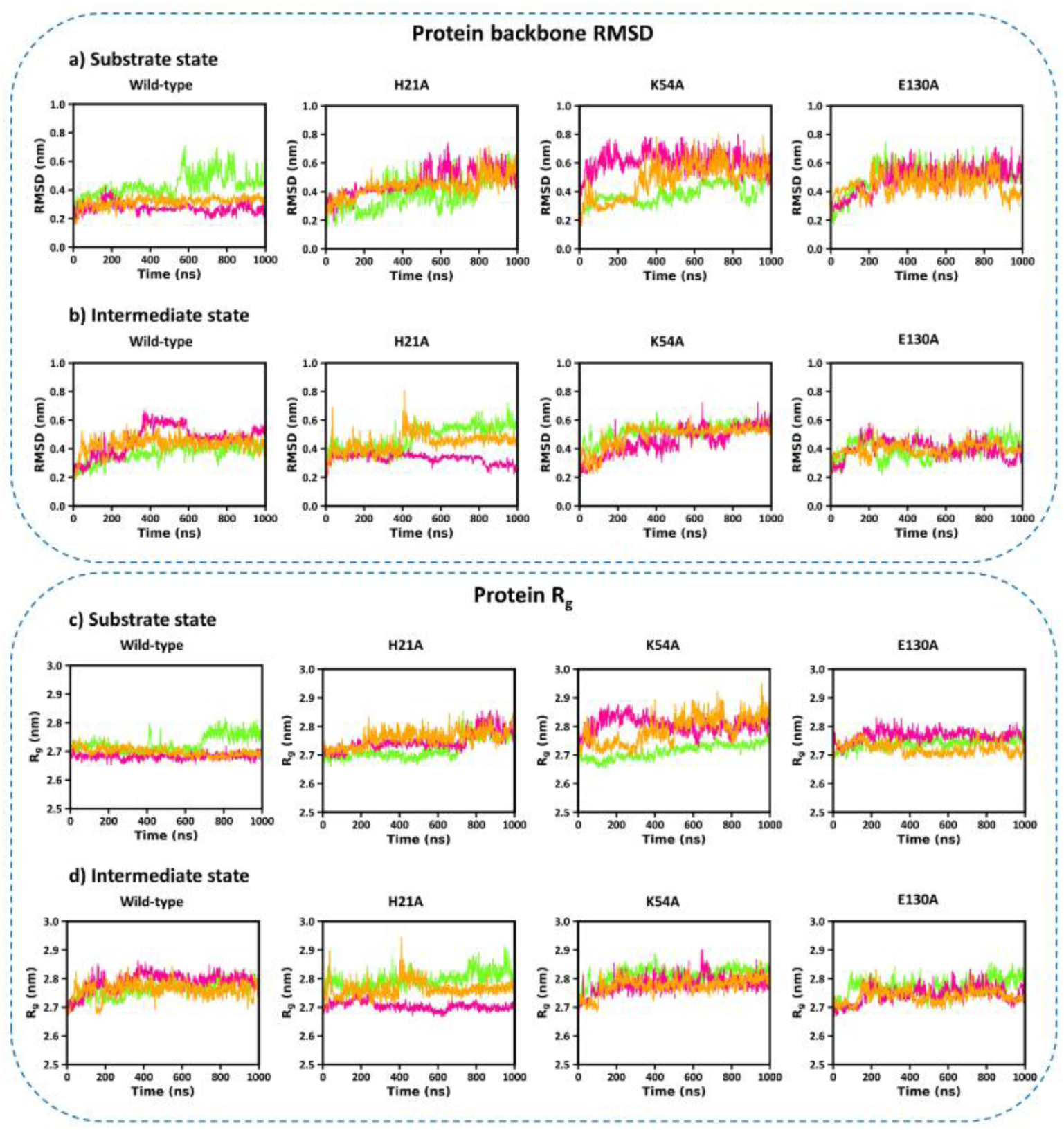
RMSD and R_g_ of the protein in wild-type and mutant models (H21A, K54A, and E130A) of the substrate and intermediate states. **(a)** RMSD in substrate state. **(b)** RMSD in intermediate state. **(c)** R_g_ in substrate state **(d)** R_g_ in intermediate state. Three MD runs of each system are represented in green, pink and orange, respectively, throughout the manuscript.

**Figure 3.**
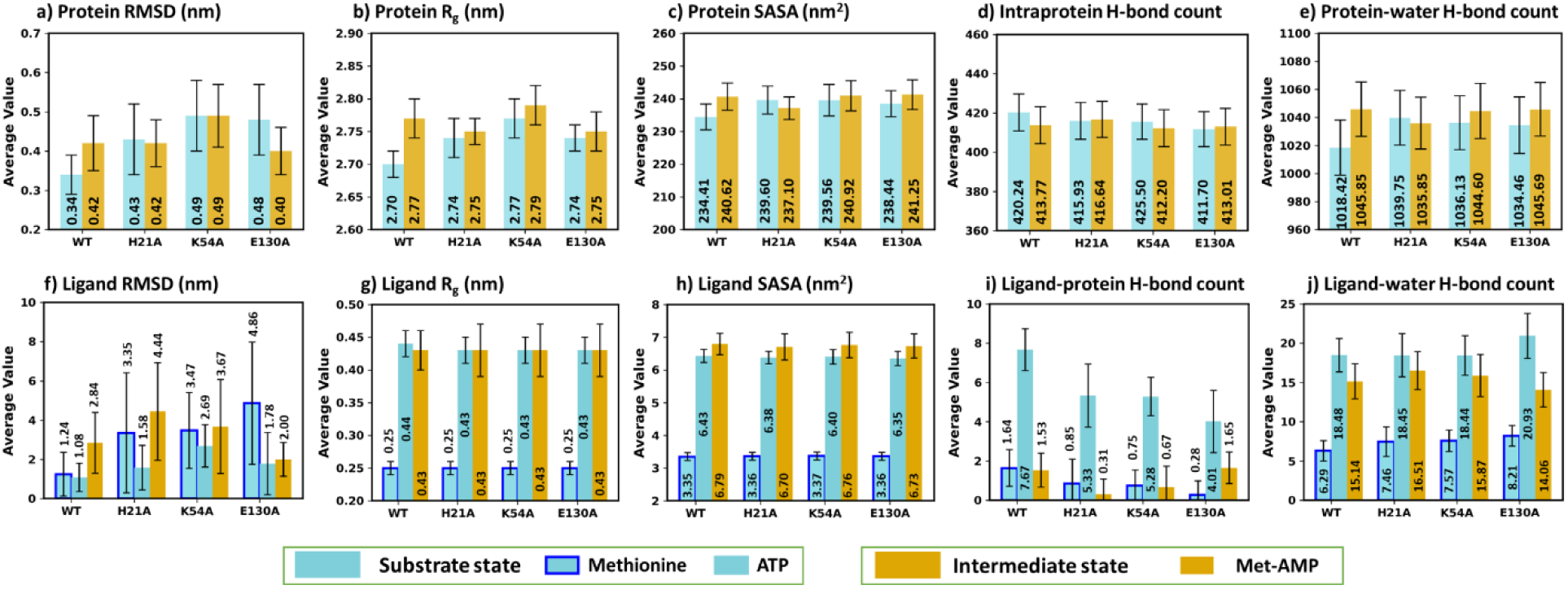
Average and standard deviation of three simulations for substrate and intermediate complexes. **(a)** Protein RMSD. **(b)** Protein R_g_. **(c)** Protein SASA. **(d)** Number of intraprotein hydrogen bond. **(e)** Protein-water hydrogen bond count. **(f)** Ligand RMSD. **(g)** Ligand R_g_. **(h)** Ligand SASA. **(i)** Protein-ligand hydrogen bond count. **(j)** Ligand-water hydrogen bond count. Average values for the substrate and intermediate states are displayed as cyan and yellow bars, respectively, with standard deviation error bars. The two ligands in the substrate state, methionine and ATP, are shown with (blue) and without borders, respectively.

The mutant complexes revealed comparatively stable behavior in the intermediate state (**Figure 2a, b**), with the main effect observed in E130A. The average RMSD for E130A mutant protein was 0.48 nm in the substrate state and 0.40 nm in the intermediate (**Figure 3a**). Interestingly, E130A protein in the intermediate state (RMSD = 0.40 nm) exhibited even higher stability than the WT (RMSD = 0.42 nm). E130 is a highly conserved residue in the CP domain (residues 116-225) and contributes to stabilizing the active site for adenylation activity through hydrogen bonding with the catalytic domain residue Lys53 [20]. Therefore, mutating this residue may disrupt the active site organization for the first step of conversion to Met-AMP. However, once Met-AMP is formed, E130A appears to eventually stabilize the protein.

The K54A mutant exhibited the highest protein RMSD (0.49 nm) in both states (**Figure 3a**). K54 forms a salt-bridge with the phosphate oxygen of ATP at active site [42] and participates in intraprotein hydrogen bond [20]. Its substitution with alanine disrupts these stabilizing interactions, resulting in increased structural fluctuations.

Within the substrate complexes (cyan bars in **Figure 3a**), WT showed a lower RMSD (average = 0.34 nm) than all mutants (average = 0.43 to 0.48 nm), suggesting higher stability of the WT protein in the substrate state. However, the intermediate state (yellow bars in **Figure 3a**) did not reveal any such trend. In addition, the WT protein in the substrate state showed the lowest RMSD (0.34 nm) compared to seven other states (0.40-0.49 nm), indicating its highest stability.

In line with the RMSD results, the R_g_ of the WT protein was lower in the substrate state (average = 2.70 nm) than the intermediate state (2.77 nm) (**Figure 2c-d, 3b** and **Table S2-S3**), suggesting a more compact structure in the substrate-bound form. Similarly, the mutant proteins consistently showed a minor increase in R_g_ in the intermediate states than the substrate states (**Figure 3b**). This indicates that the MetRS in the presence of Met-AMP exhibits a slightly open structure compared to the substrate-bound state (i.e., methionine and ATP). Within the substrate complexes (cyan bars in **Figure 3b**), the R_g_ of WT (2.70 nm) was lower than those of mutants (2.74-2.77 nm). The intermediate showed no such relation. Additionally, the WT protein in the substrate complex exhibited the lowest R_g_ (2.70 nm) among all eight states (other states R_g_: 2.74- 2.79 nm), indicating its highest structural compactness.

SASA analysis further supported these trends. The WT protein had a lower SASA in the substrate state (average = 234 nm^2^) than in the intermediate state (241 nm^2^) (**Figure 3c, S3**). This shows that the WT protein in substrate state maintains a comparatively compact state, while the presence of Met-AMP induces conformational changes leading to its higher solvent exposure. In contrast, the mutants displayed inconsistent SASA trends between the two states.

The comparison of substrate complexes showed that the WT protein SASA (234 nm^2^) was lower than that of three mutants (238-240 nm^2^) (**Figure 3c**), reflecting that mutations lead to a slightly more open structure due to the decreased stability. Consistent with the RMSD and R_g_ trends, the WT substrate complex had the lowest protein SASA (234 nm^2^) among all eight systems (SASA of others: 237-241 nm^2^), indicating its greater compactness and stability.

Collectively, these results indicate that the WT substrate protein is more stable than the intermediate state. This may be because of the fact that Met-AMP is a reaction intermediate and must undergo further conversion during tRNA charging [41]. However, the substrate entry into the MetRS active site is known to induce conformational rearrangements to promote substrate binding [69]. The elevated conformational dynamics observed in the mutant substrate states could be linked to the loss of pyrophosphate-ATP exchange activity, [20] suggesting impaired substrate recognition. In contrast, the mutants had little to no effect on protein in the intermediate state. While K54A appeared to induce the highest conformational changes in both states, E130A in the intermediate state contributed to even increased stability compared to WT.

### 3.4 Intraprotein interactions correlated with increased WT protein stability in the substrate state

The analysis of intraprotein hydrogen bonds revealed a higher average number in the WT protein of the substrate complex (average = 420) compared to the intermediate state (414) (**Figure 3d, S4**). This supports more globular and compact conformation of the WT protein in the substrate state. The H21A and E130A mutants showed slightly reduced average hydrogen bonds in the substrate (416 for H21A and 412 for E130A) compared to their respective intermediate states (417 for H21A and 413 for E130A), indicating reduced stability of these mutants in the substrate state. In contrast, the K54A mutant exhibited a higher number of hydrogen bonds in the substrate state (416) than the intermediate (412). K54 exhibited large conformational movement in both substrate and intermediate states (indicated by RMSD), but with more intraprotein hydrogen bonding in the substrate form.

The comparison of the substrate complexes (cyan bars in **Figure 3d**) showed that the WT protein had the highest intraprotein hydrogen bonds (average = 420) compared to the three mutants (412-416) (**Figure 3d**). In addition, the WT substrate complex exhibited the highest average intraprotein hydrogen bond count (420) among all eight systems (other systems = 412-417), suggesting that the WT protein in the substrate state is the most stable. It maintains a more globular structure, reduced solvent exposure, and the highest intraprotein hydrogen bonding.

### 3.5 Protein-water hydrogen bonds confirmed highest WT protein stability in the substrate state

Protein-water hydrogen bonds were analyzed over the final 500 ns of simulations, and their averages were compared (**Figure 3e, S5** and **Table S3**). The WT substrate state showed the lowest number of protein-water hydrogen bonds (average = 1018) compared to the seven other complexes (1035-1046). This coincides with the earlier observation that the WT protein in the substrate state exhibits the highest stability. The WT substrate state formed lesser hydrogen bonds (1018) than the WT intermediate state (1046), further establishing the conclusion that the substrate-bound WT protein adopts a more globular and compact conformation.

In the substrate state, mutations led to an increase in protein-water interactions (1035-1040) relative to the WT (1018). However, no clear trend was observed between the WT and mutants for the intermediate state. These results indicate that the major effects of the mutations occur in the substrate-bound form, where the WT protein maintains a stable, compact structure, while the mutants destabilize this conformation by inducing a more open and solvent-exposed state.

### 3.6 Mutants destabilized the protein in the substrate state but stabilized it in the intermediate state

To understand the dominant motions in the eight MetRS states, we performed free energy landscape principal component analysis (FEL-PCA) of protein dynamics over the final 500 ns of each simulation (**Figure 4**). The eigenvector versus eigenvalue plot showed that at least the first five eigenvectors (EV1-EV5) accounted for the majority of motion in the mutant proteins of the substrate state. In contrast, for all intermediate states, the first two eigenvectors (EV1 and EV2) captured most of the protein motion. One of the WT substrate simulations (green) exhibited higher movements during the last 500 ns run, whereas the other two replicates (pink and orange) showed lower eigenvalues (**Figure 4a**; top left). These observations suggest that MetRS mutants exhibit comparatively higher motion in the substrate state, while displaying more stable behavior in the intermediate states. This is likely due to the disruption of adenylation activity in the substrate complex, and this effect is minimized after the Met-AMP formation.

**Figure 4.**
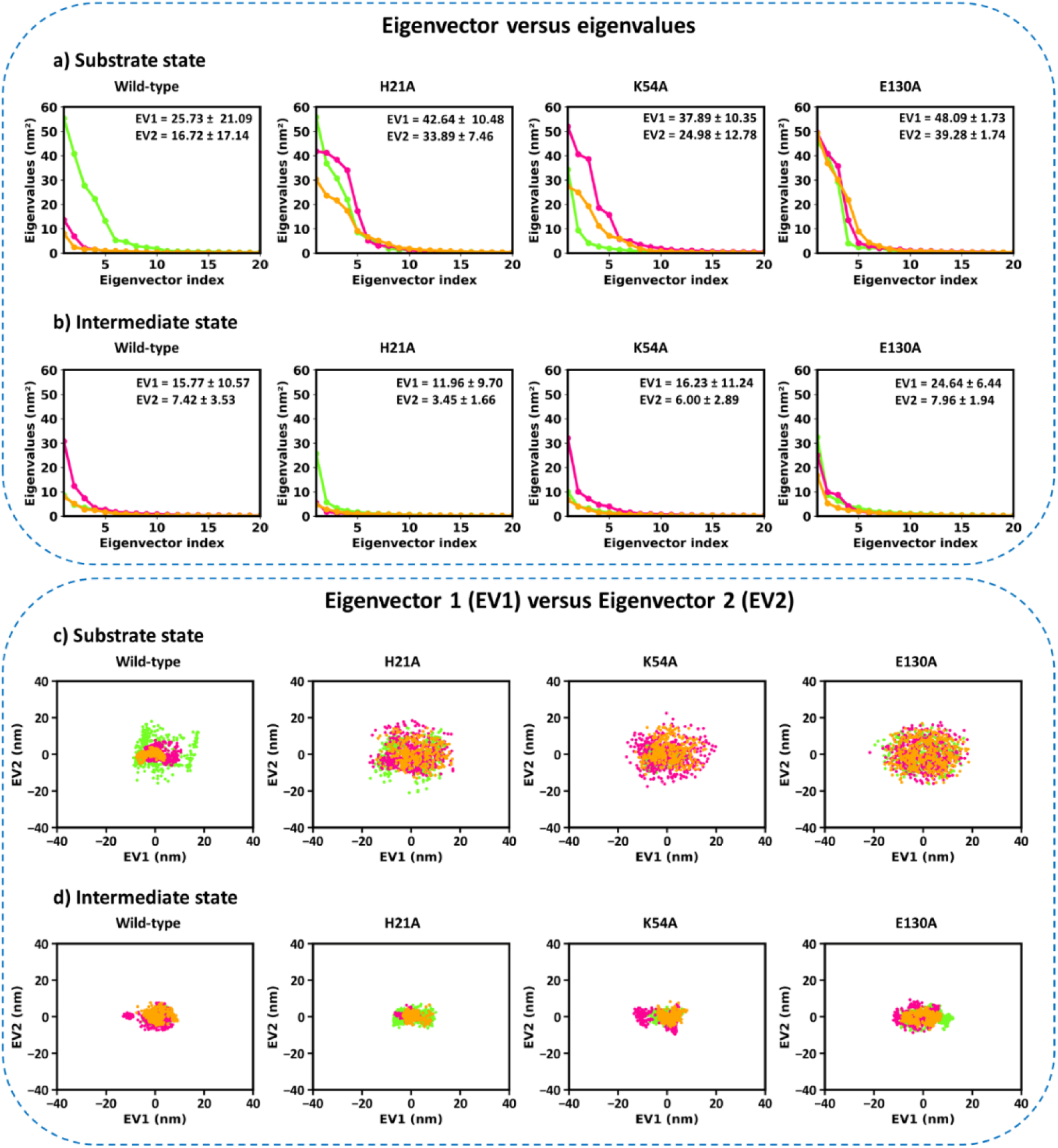
Free energy landscape principal component analysis of protein in wild-type and mutant models (H21A, K54A and E130A). Eigenvalue versus eigenvector plots for the first twenty principal components in: **(a)** substrate and **(b)** intermediate states. Conformational space projected onto the first two principal components (EV1 and EV2) in: **(c)** substrate and **(d)** intermediate states.

Comparisons among the substrate states revealed that the WT protein exhibited lower average values of EV1 (25.73 nm^2^) and EV2 (16.72 nm^2^) than the mutants (EV1 = 37.89-48.09 nm^2^ and EV2 = 24.98-39.28 nm^2^) (**Figure 4a, b**). These results show the relatively higher stability of the WT protein compared to the mutants in the substrate state, with the mutants inducing increased protein motion that may interfere with functional activity. EV1 and EV2 captured the most significant motions across all eight protein states. To explore the conformational landscape, we plotted EV1 versus EV2 (**Figure 4c, d**). Consistent with the previous observations, the mutant substrate states displayed a larger conformational landscape compared to their intermediate counterparts. Within the substrate state, the WT protein showed a more restricted conformational space than the mutants.

Overall, these results consistently highlight that the mutations destabilize the protein in the substrate state, likely impairing its pyrophosphate-ATP exchange activity. However, once the intermediate state is reached, the mutations have minimal effect on protein stability, suggesting that they are unlikely to impact the subsequent aminoacylation step.

### 3.7 Mutants exhibited higher fluctuations in the CP domain and KMSKS region in the substrate state

To identify the protein regions showing higher motion, we performed RMSF analysis on the residues over the final 500 ns runs (**Figure S6**). Since the primary effects of the mutations were observed in the substrate state, we compared RMSF values for the mutants relative to the WT in both substrate and intermediate states. Interestingly, the two regions displayed consistently higher fluctuations across all states: 126-156 (part of the CP domain) and 296-306 (the KMSKS region) (**Figure S6**). These regions showed more movements in the mutant proteins compared to the WT in the substrate state. The CP domain and KMSKS region are important for the integration of the MetRS active site, [20,70,71] and play key roles in both substrate activation and the subsequent acylation reaction through their conformational flexibility [70–72]. The H21A, K54A and E130A mutations are located within the catalytic and CP domains. They appear to induce conformational instability in these regions, thereby inactivating the substrate pyrophosphate-ATP exchange activity. They likely work by affecting domain integrity by disturbing the intraprotein interaction network, as further discussed below.

### 3.8 Ligand analysis revealed longer residence time of methionine and ATP in the active site compared to Met-AMP

To examine how protein motion correlates with ligand dynamics and how mutations affect ligand behavior, we analyzed the ligand properties (**Figure 5, 3f-h, S7-S8** and **Table S4-S7**). Three ligands, including methionine, ATP and Met-AMP were studied. Methionine and ATP are present in the substrate state, whereas Met-AMP is present in the intermediate. The primary analysis was performed on the full 1 µs simulations (RMSD, R_g_ and SASA).

**Figure 5.**
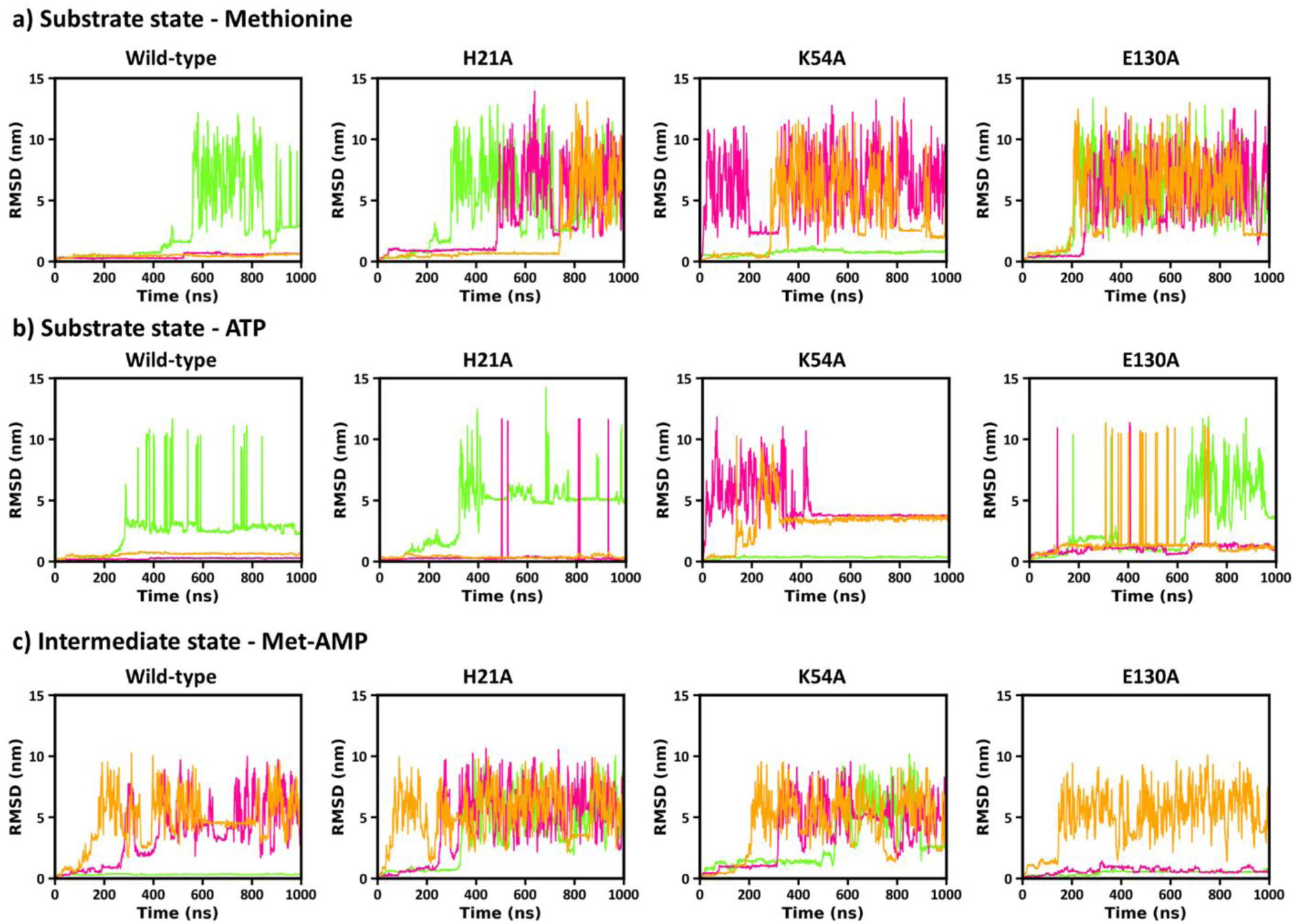
RMSD of ligands with respect to the protein in wild-type and mutant models (H21A, K54A and E130A) of the substrate and intermediate states. **(a)** Methionine (in the substrate state). **(b)** ATP (in the substrate state), **(c)** Met-AMP (in the intermediate state).

In general, methionine showed stable RMSD in the WT (average of three runs = 1.24 nm) compared to mutants (3.35-4.86 nm) in the substrate state (cyan bars with blue borders in **Figure 3f**; time series in **Figure 5a**). The increased protein flexibility in the mutants led to destabilization and eventual release of methionine from the active site (**Figure 5a**). These results align with experimental data showing loss of catalytic activity in these mutants [20]. However, as observed for protein, one WT trajectory (green) exhibited elevated motion during the second half of the simulation.

ATP is also present in the substrate state. It followed dynamics similar to those of the protein and methionine (**Figure 5b** and **3f, Table S6-S7**). The average ATP RMSD (of three runs) in WT (1.08 nm) was lower than in the mutants (1.58-2.69 nm), indicating a comparatively stable ATP conformation in the WT system. ATP also exhibited a longer residence time in the WT compared to the mutants (**Figure 5b**). The R_g_ and SASA of ATP remained comparable in all states.

A comparison of methionine (substrate; RMSD: 1.24 nm) and Met-AMP (intermediate; RMSD: 2.84 nm) in WT MetRS revealed that methionine was more stably retained in the protein’s active site (**Figure 3f**). This behavior strongly correlated to the protein RMSD trends discussed earlier. Methionine remained in the active site for a substantial time, while Met-AMP left the site in a shorter period (**Figure 5a, 5c**), indicating longer and shorter residence times for the substrate and intermediate, respectively. These findings also reflect the intermediate nature of Met-AMP, which must undergo further reaction in the aminoacylation pathway.

Notably, Met-AMP in E130A mutant remained at active site in two simulations, with an average RMSD of 2.00 nm, which is even lower than the WT state (2.84 nm) (**Figure 5c, 3f**). This suggests that E130A may stabilize Met-AMP. However, it destabilized methionine (substrate state), indicating its possible role in the early stages of substrate recognition or activation.

Both R_g_ and SASA displayed higher variability for Met-AMP than for methionine (**Figure S7-S8, 3g, 3h**), likely reflecting the lower stability of Met-AMP at the protein’s active site.

These findings lead to the concept that both ATP and methionine have longer residence times in MetRS, i.e., in the substrate state. In contrast, Met-AMP exhibits a shorter residence time. The mutants act by destabilizing both methionine and ATP at their binding sites, an effect observed primarily in the substrate state. In contrast, no such destabilizing trend was observed in the intermediate state, suggesting that the mutations primarily impair pyrophosphate-ATP exchange activity [20].

### 3.9 The WT substrate state showed more protein-ligand interactions than the mutants

We analyzed protein-ligand hydrogen bonds over the final 500 ns of simulations for methionine, Met-AMP, and ATP (**Figures S9** and **3i**). Methionine formed the highest average number of hydrogen bonds with the WT protein (1.64), compared to the mutants (0.28-0.85), suggesting stabilization of the productive state in WT MetRS, which appears disrupted in the mutants [20,47].

Met-AMP formed a comparable number of hydrogen bonds with the WT (1.53) and E130A (1.65) proteins, indicating that the E130A mutant may help stabilize Met-AMP at the active site. In contrast, the H21A and K54A mutants showed lower average hydrogen bond counts with Met- AMP (0.31 and 0.67, respectively), suggesting reduced stability of Met-AMP in these mutants.

ATP formed the highest average number of hydrogen bonds with the WT protein (7.67), compared to the mutants (4.01-5.33), supporting its greater stability in the WT complex. This interaction network appears to play a significant role in substrate activation and the productive state, [20,47] which is compromised in the mutants due to destabilization of domain integrity surrounding the protein’s active site.

The strong hydrogen bonding of methionine and ATP to the WT protein likely reflect an induced fit mechanism for substrate recognition by MetRS [20,42,73]. The presence of mutations disrupts this binding by increasing flexibility in the CP domain and KMSKS region, leading to the increased protein size and significantly decreased ligand residence times at the active site. This possibly forms an unproductive protein-ligand conformation, thereby disrupting pyrophosphate- ATP exchange activity by an early release of ligands. This differential recognition mechanism was further supported by the analysis of representative structures from the MD simulations (**Figure S10-S11**). **Figure 6a and b** illustrate this mechanism using the representative structures.

**Figure 6.**
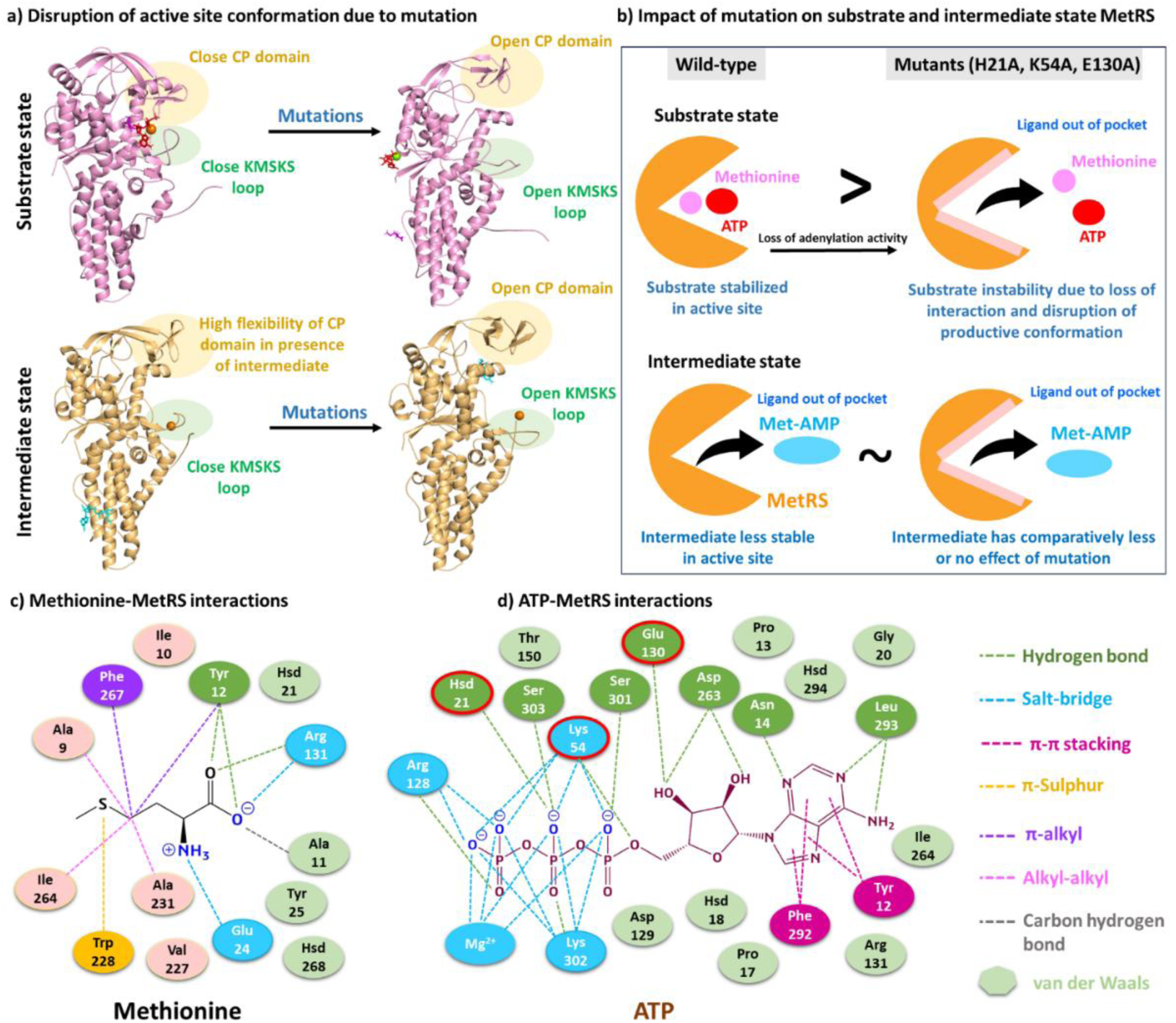
Protein-ligand interactions in simulations and the summary of the current work. **(a)** The substrate state maintains a productive conformation by an induced-fit mechanism, in which the CP domain and KMSKS loop adopt a closed active site conformation and form strong hydrogen bonding with methionine and ATP. In contrast, the Met-AMP intermediate reduces the stability of the CP domain and KMSKS loop, leading to opening of the active site and decreased stability of Met-AMP. The H21A, K54A, and E130A mutations open the CP domain and KMSKS loop, disrupting the active site integration and preventing the formation of a productive conformation for methionine and ATP reaction. **(b)** Methionine and ATP bind stably to MetRS in the substrate state to facilitate the pyrophosphate-ATP exchange reaction. The H21A, K54A, and E130A mutations impair this activity by reducing the residence time of the ligands at the active site. The Met-AMP intermediate shows reduced residence time at the MetRS active site, possibly due to its transient nature. Once the intermediate is formed, these mutations have little or no effect and may even increase stability of Met-AMP-MetRS complex. **(c)** Methionine and **(d)** ATP interactions at the active site of MetRS.

### 3.10 Protein-ligand interactions in the productive conformation

Specific interactions of methionine and ATP with the protein were analyzed in the WT substrate state (**Figure 6c**) using representative structures from MD simulations. Methionine formed two hydrogen bonds with Tyr12 and salt-bridges with Glu24 and Arg131. In contrast, ATP formed salt-bridges with Lys54, Arg128, and Lys302, and hydrogen bonds with seven protein residues including Asn14, Hsd21, Glu130, Asp263, Leu293, Ser301, and Ser303. These interacting residues are located in the catalytic domain (residues 1-115 and 226-292), the CP domain (116-225) and the KMSKS region (293-350). These three regions play a crucial role in the substrate activation [20]. Hsd21 is part of the conserved _18_HVGH_21_ motif and is known to stabilize ATP phosphate chain during the adenylation reaction [47]. In our analysis, we observed a hydrogen bond between Hsd21 and ATP. Lys54 is known to participate in salt-bridge with the phosphate oxygen of ATP at the active site, [42] and we observed multiple such interactions. The KMSKS motif (299-303) has been reported to neutralize the negative charges on ATP, thereby facilitating its reaction with methionine [74]. Consistently, we observed multiple electrostatic interactions between Lys302 and O^-^ atoms of ATP phosphates (**Figure 6d**). In addition, Mg²⁺ has been reported to stabilize ATP at the active site [57]. In agreement, we found that the Mg²⁺ ion was consistently formed many electrostatic interactions with the ATP phosphate group (**Figure 6d**).

These strong intermolecular interactions likely contribute to productive protein-ligand binding and substrate activation. Three residues, Hsd21, Lys54, and Glu130 interacted directly with ATP (highlighted in red border in **Figure 6d**). Hsd21 and Glu130 stabilized ATP with hydrogen bonds, while Lys54 neutralized its negative charges. Mutations of these residues to alanine disrupted the stability of both ATP and methionine in the active site, thereby inactivating the productive state. We propose that electrostatic and hydrogen bonding interactions are necessary for achieving and maintaining the productive state of MetRS.

## 4 Conclusion

MetRS is an essential target in *Mtb* due to its involvement in protein synthesis. In this study, we identified the molecular cause for the differential dynamic and binding features between its substrate and intermediate states, and the mechanism of three mutants (H21A, K54A, and E130A). The substrate state protein was more stable than the intermediate, with a longer residence time of methionine and ATP (substrate state) at the protein’s binding sites. In contrast, the intermediate product (Met-AMP) remained at the active site for a comparatively shorter duration. The mutant proteins were more unstable in the substrate state compared to the intermediate state. These mutations worked at the substrate complex by destabilizing the protein-ligand interactions and impairing the adenylation activity. Interestingly, once the intermediate Met-AMP was formed, the mutants had little or no disruptive effect. They may even stabilize Met-AMP (the case of E130A mutant). In summary, this study establishes a mechanistic framework that explains the distinctive recognition of the substrate and intermediate in *Mtb* MetRS, as well as the functional consequences of specific point mutations.

## Author contributions

RM conceptualized and supervised the work. ST performed the work. RM and ST analyzed the work and wrote the manuscript.

## Acknowledgements

RM acknowledges Science and Engineering Research Board, Government of India, for Start-up Research Grant (SRG) SRG/2022/000304 and IIT Bhilai for Research Initiation Grant (RIG), 2005900. ST acknowledges the Ministry of Education, Government of India, for the PMRF fellowship grant 5001800.

## Data availability statement

**Supplementary information.** Detailed analysis is provided in the supplementary file. It contains method of multiple sequence alignment (Section 1), results of phylogenetic analysis (Section 2), hydrophobic and hydrophilic SASA (Section 3), N to C terminal distance analysis of protein (Section 4), mutation effects on catalytic and connective peptide domain (Section 5), ligand-water hydrogen bond and RMSF analysis (Section 6), hydrogen bond analysis of mutated residues (Section 7), secondary structure analysis (Section 8), Protein binding site volume analysis (Section 9), protein-ligand interaction (Section 10), and trajectory visualization (section 11).

