## Supplementary file for "Distinct substrate and intermediate recognition via mutation effects on *Mycobacterium tuberculosis* methionyl-tRNA synthetase"

*Rukmankesh Mehra: 0000-0001-6010-1514*

*Shivani Thakur: 0000-0002-7869-6648*

**Supplementary Section 1. Multiple sequence alignment of MetRS sequences**

We identified conservation scores of *Mtb* MetRS residues to determine crucial regions and residues required for the proper functioning of the enzyme. Conservation score calculations involved multiple sequence alignment (using the MAFFT algorithm) of *Mtb* MetRS with other homologous MetRS (total 150) in the CONSURF server [1–3]. The program uses the Bayesian method for calculating conservation scores and selects homologous with minimum query coverage of 60% and sequence identity of 35%.

Further, for multiple sequence alignment, 26 MetRS protein sequences from different organisms, including mycobacteria, gram-positive and gram-negative bacteria, eukaryotes, archaea, etc., were downloaded from UniProt (**Table S1**) [4,5]. Multiple sequence alignment was performed using Clustal Omega [6] to analyze the area of similarity and identify highly conserved motif/regions in MetRS of different eukaryotic and prokaryotic species. Gram-positive bacterial MetRS included *Staphylococcus aureus, Staphylococcus saprophyticus, Geobacillus stearothermophilus, Bacillus subtilis, Streptococcus pneumoniae, Bacillus cereus, Bifidobacterium longum* and *Streptococcus pyogenes*. Gram-negative bacterial MetRS included *Escherichia coli, Aquifex aeolicus, Thermus thermophilus, Helicobacter pylori,* and *Haemophilus influenzae*. Eukaryotic mitochondrial MetRS included *Homo sapiens* (mitochondrial), *Candida albicans* (mitochondrial), and *Saccharomyces cerevisiae* (mitochondrial). Eukaryotic cytoplasmic MetRS included *Saccharomyces cerevisiae* (cytoplasmic) and *Homo sapiens* (cytoplasmic). Archaea MetRS contains *Pyrococcus abyssi*. Phylogenetic analysis was performed to determine substitutions during evolution and understand the evolutionary relationship among different MetRS protein sequences. A guide tree in rectangular format was built along with branch length using the iTOL v7 [7,8] program.

**Supplementary Section 2. Multiple sequence alignment and phylogenetic analysis**

The sequence alignment results showed the overall evolutionary conserved nature of MetRS protein in different organisms (**Figure 1c, S1-S2**). The H21, K54 and E130 residues were retained in prokaryotic MetRS and eukaryotic mitochondrial MetRS but not in eukaryotic cytoplasmic MetRS (**Figure S1**). His21 residue (signature motif HIGH) was conserved in all MetRS sequences except for cytoplasmic eukaryotic and *Streptococcus pyogenes* MetRS. The other two residues (K54 and E130) were not conserved in some gram-positive and gram-negative MetRS. The amino acid sequence of *Mycobacterium* species was found to be preserved during evolution. These species formed a tight cluster, indicating a conserved MetRS sequence within the *Mycobacterium* genus, with minimal divergence between strains like *Mtb* H37Rv and H37Ra (**Figure S2**). The longest MetRS protein (homo sapiens cytoplasmic) has the least similarity with other prokaryotic MetRS. Eukaryotic mitochondrial MetRS proteins are distinct but share a closer evolutionary relationship with bacterial MetRS sequences, supporting the endosymbiotic origin of mitochondria [9,10]. Notably, *Mtb* MetRS has the least sequence identity with cytoplasmic MetRS of *Homo sapiens* and *Saccharomyces cerevisiae*, suggesting evolutionary distance and potential differences in MetRS function across different taxa. MetRS sequences associated with MetRS1 and MetRS2 families formed separate clusters and showed higher sequence similarity among themselves (**classification shown in Table S1 and Figure S2**). MetRS1 protein is generally found in gram-positive bacteria, mitochondria, and protozoa, whereas MetRS2 is mainly found in gram-negative bacteria, eukaryotes, specifically in cytoplasm and archaea. Additional insertion of 20 amino acids in the CP domain of MetS2 leads to significant differences in sequences of MetRS1 and MetRS2 proteins [11,12]. Since MetRS plays a critical role in translation, its conservation across species underscores its essential nature.

**Supplementary Section 3. Hydrophobic and hydrophilic SASA**

SASA of the hydrophobic and hydrophilic residues was computed for the last 500 ns runs, which showed minor differential effects (**Figure S12-S14**). Hydrophobic SASA of the WT protein was lower in the substrate complex (195 $nm^2$) than the intermediate (197 $nm^2$) (**Table S2-S3**). However, the mutant comparisons showed largely consistent values. In addition, hydrophilic SASA did not reveal any consistent trend.

However, within the substrate states, WT protein showed lower hydrophilic SASA (average = 284 $nm^2$) compared to mutants (285-287 $nm^2$). The hydrophilic exposure increased upon mutations.

This suggests that MetRS maintains a comparatively more globular state in the substrate state, which was slightly disturbed in the intermediate with increased hydrophobic residue exposure. The mutants seem to disrupt activity in the substrate state (as observed above). These might be linked to their increased hydrophilic residue exposure.

**Supplementary Section 4. N to C terminal distance analysis of protein over the simulations**

The N to C terminal distance revealed mutations H21A and K54A do not significantly alter the N to C terminal distance compared to the wild-type (**Figure S14-15** and **Table S2-S3**). This suggests that the two mutations do not impact the overall protein conformation in the substrate state. In contrast, the E130A mutation shows a marked effect, with average N to C terminal distances of 5.72 ± 0.44 nm in the substrate state and 5.54 ± 0.55 nm in the intermediate state. These increased distances indicate a more extended or open protein conformation relative to the wild-type.

**Supplementary Section 5. Mutation effects on catalytic and connective peptide domain**

Detailed analyses explored how specific mutations impact the catalytic and connective peptide (CP) domains. RMSD calculations revealed that the CP domain exhibits greater flexibility in the intermediate state compared to the substrate state (**Figures S16-S17**). Mutations H21A, K54A, and E130A enhanced the flexibility of the CP domain in both states (**Figure S6**).

Average RMSF values of the catalytic domain residues showed that the wild-type protein maintained lower flexibility in the substrate state (**Figure S18a**). In contrast, H21A and K54A mutants exhibited increased flexibility, particularly for the loop region (residues 9-25) and a loop-helix region (residues 48-65), which include key substrate-binding residues. The E130A mutant had minimal impact on the catalytic domain, displaying flexibility similar to the wild type. In the intermediate state, mutations K54A and E130A showed increased flexibility for catalytic domain residues, particularly in the loop (residues 10-20) and helix-associated (residues 50-70) regions. At the same time, H21A reduced flexibility relative to the wild-type (**Figure S18b**).

H21A mutation caused very high fluctuations (~6 Å) in the CP domain residues (120-155) in the substrate-bound state, pointing to noticeable conformational changes in this region (**Figure S18c**). Interestingly, the E130A mutation reduced the flexibility of surrounding residues compared to the wild-type. Additionally, increased flexibility of loops L1 (residue 126-130) and L2 (residue 133-138) for H21A mutation suggest major structural rearrangements of these loops. In the intermediate state, CP domain analysis (residues 116-225) revealed that loops (L1 and L2) in the

125-150 region are stabilized by E130A mutation compared to wild-type. Other mutations displayed mixed effects (**Figure S18d**).

### Supplementary section 6. Ligand-water hydrogen bond and RMSF analysis

The ligand-water interactions revealed that methionine formed more ligand-water hydrogen bonds in mutant models (particularly E130A), indicating more solvent exposure of the ligand (**Figure S19, 3j** and **Table S5**). The average number of ATP-water hydrogen bonds remained largely unchanged across all systems, although E130A displayed a slightly higher value (20.93), suggesting increased solvent exposure (**Figure S19, 3j** and **Table S7**). The higher standard deviations observed for H21A and E130A also point to greater fluctuations in ATP's solvent accessibility. For Met-AMP, H21A and K54A mutations showed a similar trend in ligand-water hydrogen bonds, while E130A had a comparatively less solvent interaction (**Figure 3j**).

Ligand RMSF showed higher fluctuations in the intermediate state, suggesting greater flexibility of Met-AMP. Notably, H21A and K54A mutations showed increased RMSF, indicating destabilization of Met-AMP (**Figure S20-S21**).

### Supplementary section 7. Hydrogen bond analysis of mutated residues

**Figures S22-S24** illustrate hydrogen bond interactions between residues H21, K54, and E130 with the ligands in both substrate (methionine and ATP) and intermediate (Met-AMP) states. His21 formed weak hydrogen bonds with methionine and Met-AMP in the wild-type, which were lost entirely in the H21A mutant model (**Figure S22**). ATP showed weak hydrogen bonding with His21 residue in the wild-type, H21A, and E130A mutant models, while no interactions were observed in the K54A mutant model.

Lys54 showed no hydrogen bonding with Met-AMP in either wild-type or mutant forms, possibly due to alternative interaction types (**Figure S23**). However, in H21A and E130A mutants, Lys54 exhibited hydrogen bonding with methionine, suggesting that these mutations may induce conformational changes that bring Lys54 closer to the ligand. Lys54 frequently formed two to three hydrogen bonds with ATP in the wild-type, which reduced to one or two hydrogen bonds in the H21A and E130A mutants. Interestingly, mutating Lys54 to alanine abolished its interactions with all ligand molecules (methionine, ATP, and Met-AMP). Lys54 is an important residue that forms a salt bridge with Met-AMP's negatively charged phosphate oxygen [13].

Glu130 showed weak hydrogen bonding with methionine and ATP in the wild-type but not in any mutant. Interestingly, it showed no interaction with Met-AMP in wild-type and mutant form (**Figure S24**).

These findings highlight the importance of these residues (His21, Lys54, and Glu130) in preserving protein-ligand binding. The enzymatic function may be impacted if these residues are mutated since they might disrupt essential interactions required for amino acid activation.

**Supplementary section 8. Secondary structure analysis of protein's representative structures**

Secondary structure analysis of representative structure protein showed minor changes in the secondary structural elements during simulations (**Figure S25-S28**). The overall secondary structure was largely preserved in both the substrate and intermediate state. The H21A and K54A mutation does not affect the secondary structure of its surrounding residues, as α-helix was maintained during simulations. The residues near E130 showed continuous conversion of turn ↔ $3_{10}$-helix in both states. Similarly, turn ↔ $3_{10}$-helix conversion was also observed in two other protein regions (residues 125-135 and 155-165). The residue in the range 40-50 showed loss of turn to form coil due to mutation in substrate and intermediate state. High secondary structural changes were observed for residues 272-302 of the active site, showing β-sheet ↔ coil ↔ turn conversions. Similarly, residues 340-350 also showed higher structural changes due to the loss of α-helix to form turn or $3_{10}$-helix or β-sheet. These findings suggest that the introduced mutations may impact the overall secondary structure of the protein.

**Supplementary section 9. Active pocket volume analysis of protein's representative structure**

Analysis of active site volume revealed a slightly larger binding pocket in the intermediate state compared to the substrate state of the wild-type protein (**Table S8**). Mutations led to an increase in pocket size across both states, with H21A and E130A mutations causing notable enlargement in the substrate state and K54A and E130A contributing to increased pocket size in the intermediate state.

**Supplementary section 10. Protein-ligand interaction analysis in representative structures**

The 2D interaction plots of the methionine, ATP, and Met-AMP with protein showed interactions involved in ligand stabilization in the wild-type protein, including π-sulphur, hydrogen bonds, π-alkyl, and π-cation interactions (**Figure S29-S31**). Mutations led to the loss of specific interactions associated with the mutated residues.

Methionine-protein interactions showed that E130A mutation resulted in the disappearance of nearly all interactions with methionine (**Figure S29**). In the wild-type, methionine binding was primarily stabilized by hydrogen bonding and various other interactions within the active site.

ATP-protein interaction plots revealed extensive interactions in the wild-type protein, primarily dominated by electrostatic contacts between the ATP phosphate groups and charged protein residues (**Figure S30**). The $Mg^{2+}$ ion consistently engaged in multiple electrostatic interactions with ATP phosphates across wild-type and mutant models. In the wild-type, ATP formed 6 to 9 hydrogen bonds with protein residues, which decreased to 2 or 3 hydrogen bonds in the K54A mutant and 6 or 7 hydrogen bonds in the H21A and E130A mutants across different runs. Although hydrogen bonding patterns changed, most electrostatic interactions involving ATP phosphates were retained in the mutants, albeit with different interacting residues. Overall, the analysis highlights the roles of H21, K54, and E130 in mediating interactions with either ATP or methionine, depending on the active site conformation.

Met-AMP interactions with protein residues were analyzed in three representative structures across different protein variants (**Figure S31**). Met-AMP formed relatively few interactions during the simulation in the wild-type protein, and most of these interactions were further diminished in the mutant models. A maximum of 3 to 4 hydrogen bonds were observed in one of the wild-type and E130A mutant structures. The lack of stable interactions indicates weak and unstable binding of Met-AMP at the active site.

**Supplementary section 11. Trajectory visualization**

The visualization of whole trajectory containing 1000 frames (one for each ns) showed that the substrate (methionine and ATP) as well as intermediate (Met-AMP) molecule came out of the binding site after sometime during simulations. For the WT structure, methionine and ATP were out of pocket only in one simulation (green); however, in mutant structures, both molecules came out of pocket in all three runs. This shows mutation has significantly affected substrate binding to MetRS protein by disrupting the essential interactions observed in Galyna *et al.* [14] Small molecules like methionine and its analogs showed high RMSD and fluctuation in *Proteus mirabilis* MetRS simulations [15]. The weak binding of methionine and ATP molecule to MetRS protein can be explained based on previously reported high $K_m$ values of methionine and ATP for *Mtb* MetRS as compared to other organism MetRS (**Table S9**). MetRS1 proteins are reported to have enlarged and open, active site conformation due to the negatively charged surface of connective peptide knuckle facing ATP site [16]. L-Methionine binding causes bent conformation of the CP domain due to the formation of interactions. Mutation in the CP domain (E130A) might disturb L-methionine interactions, allowing easy ligand movement from the pocket.

**Table S1. Description of species used in identification of MetRS evolutionary relationships.** 1-7: *Mycobacterium* genus, 8-12: eukaryotes, 13: archaea, 14-18: gram negative, 19-26: gram-positive.

| S.N. | Name of species | UniProt Id | Length | Strain |
|---|---|---|---|---|
| 1. | *Mycobacterium tuberculosis H37Rv* **(MetRS1)** | P9WFU5 | 1-519 | ATCC 25618 / H37Rv |
| 2. | *Mycobacterium tuberculosis H37Ra* **(MetRS1)** | A5U150 | 1-519 | ATCC 25177 / H37Ra |
| 3. | *Mycobacterium tuberculosis Oshkosh* **(MetRS1)** | P9WFU4 | 1-519 | CDC 1551 / Oshkosh |
| 4. | *Mycobacterium bovis* **(MetRS1)** | P59952 | 1-519 | ATCC BAA-935 / AF2122/97 |
| 5. | *Mycolicibacterium smegmatis* **(MetRS1)** | A0R3E2 | 1-515 | ATCC 700084 / mc (2)155 |
| 6. | *Mycobacterium marinum* **(MetRS1)** | B2HDK3 | 1-518 | ATCC BAA-535 / M |
| 7. | *Mycobacterium leprae* **(MetRS1)** | Q9CD55 | 1-537 | TN |
| 8. | *Homo sapiens cytoplasmic* **(MetRS2)** | P56192 | 1-900 | |
| 9. | *Homo sapiens mitochondrial* **(MetRS1)** | Q96GW9 | 1-593 | |
| 10. | *Saccharomyces cerevisiae cytoplasmic* **(MetRS2)** | P00958 | 1-751 | ATCC 204508 / S288c |
| 11. | *Saccharomyces cerevisiae mitochondrial* **(MetRS1)** | P22438 | 1-575 | ATCC 204508 / S288c |
| 12. | *Candida albicans mitochondrial* **(MetRS1)** | O74634 | 1-577 | |
| 13. | *Pyrococcus abyssi* **(MetRS2)** | Q9V011 | 1-722 | GE5 / Orsay |
| 14. | *Escherichia coli* **(MetRS2)** | P00959 | 1-677 | K12 |
| 15. | *Thermus thermophilus* **(MetRS1)** | P23395 | 1-618 | ATCC 27634 / DSM 579 / HB8 |
| 16. | *Haemophilus influenzae* **(MetRS2)** | A5UBS8 | 1-682 | PittEE |
| 17. | *Aquifex aeolicus* **(MetRS1)** | O67298 | 1-497 | VF5 |
| 18. | *Helicobacter pylori* | Q9ZKG9 | 1-656 | J99 / ATCC 700824 |
| 19. | *Staphylococcus aureus* **(MetRS1)** | P67578 | 1-657 | Mu50 / ATCC 700699 |
| 20. | *Staphylococcus saprophyticus* | Q49UZ9 | 1-658 | ATCC 15305 / DSM 20229 / NCIMB 8711 / NCTC 7292 / S-41 |
| 21. | *Streptococcus pneumoniae* **(MetRS2)** | B2IQ55 | 1-545 | CGSP14 |
| 22. | *Streptococcus pyogenes* **(MetRS1)** | P0DG49 | 1-665 | SSI-1 |
| 23. | *Bacillus subtilis* | P37465 | 1-664 | 168 |
| 24. | *Geobacillus stearothermophilus* **(MetRS1)** | P23920 | 1-649 | |
| 25. | *Bacillus cereus* | B7HD62 | 1-544 | B4264 |
| 26. | *Bifidobacterium longum* | P59076 | 1-621 | NCC 2705 |

**Table S2. The properties of protein for each simulation run.**

| Protein properties | | Substrate state | | | | Intermediate state | | | |
|---|---|---|---|---|---|---|---|---|---|
| | | WT | H21A | K54A | E130A | WT | H21A | K54A | E130A |
| RMSD (nm) | RUN 1 | 0.43 (0.08) | 0.38 (0.11) | 0.38 (0.07) | 0.50 (0.09) | 0.36 (0.06) | 0.48 (0.09) | 0.52 (0.06) | 0.39 (0.07) |
| | RUN 2 | 0.27 (0.04) | 0.47 (0.09) | 0.61 (0.08) | 0.48 (0.09) | 0.46 (0.10) | 0.33 (0.03) | 0.45 (0.10) | 0.40 (0.06) |
| | RUN 3 | 0.32 (0.04) | 0.44 (0.07) | 0.48 (0.12) | 0.45 (0.08) | 0.43 (0.06) | 0.44 (0.07) | 0.49 (0.08) | 0.39 (0.04) |
| Radius of gyration (nm) | RUN 1 | 2.73 (0.03) | 2.72 (0.04) | 2.71 (0.02) | 2.74 (0.02) | 2.76 (0.03) | 2.80 (0.03) | 2.81 (0.03) | 2.77 (0.03) |
| | RUN 2 | 2.68 (0.01) | 2.75 (0.03) | 2.81 (0.03) | 2.77 (0.22) | 2.79 (0.03) | 2.70 (0.01) | 2.78 (0.03) | 2.74 (0.03) |
| | RUN 3 | 2.70 (0.01) | 2.76 (0.03) | 2.79 (0.05) | 2.72 (0.02) | 2.76 (0.03) | 2.76 (0.03) | 2.77 (0.03) | 2.74 (0.02) |
| Total SASA ($nm^2$) | RUN 1 | 237.75 (5.03) | 237.61 (4.25) | 236.30 (4.94) | 238.72 (3.56) | 237.97 (4.02) | 242.08 (3.78) | 241.46 (4.22) | 245.92 (5.20) |
| | RUN 2 | 233.19 (3.09) | 239.57 (4.12) | 243.01 (5.79) | 241.79 (4.39) | 242.27 (4.48) | 231.46 (3.04) | 243.68 (5.47) | 239.50 (3.95) |
| | RUN 3 | 232.29 (3.74) | 241.61 (4.53) | 239.37 (3.73) | 234.81 (4.03) | 241.60 (3.97) | 237.76 (3.54) | 237.62 (4.20) | 238.34 (4.50) |
| Hydrophobic SASA ($nm^2$) | RUN 1 | 194.73 (2.03) | 194.79 (1.99) | 194.89 (1.74) | 196.43 (1.88) | 196.79 (2.23) | 195.72 (2.02) | 194.07 (1.86) | 197.38 (2.05) |
| | RUN 2 | 195.42 (1.58) | 196.63 (2.00) | 196.89 (2.15) | 196.56 (1.99) | 196.74 (1.78) | 193.56 (1.78) | 194.76 (2.38) | 194.84 (1.81) |
| | RUN 3 | 195.65 (1.71) | 193.97 (2.13) | 194.19 (1.76) | 195.08 (2.18) | 197.06 (1.84) | 194.57 (1.80) | 193.00 (1.78) | 192.77 (1.90) |
| Hydrophilic SASA ($nm^2$) | RUN 1 | 287.36 (4.34) | 285.67 (3.31) | 285.58 (3.63) | 285.92 (4.15) | 281.40 (2.94) | 283.77 (3.08) | 284.75 (3.61) | 286.99 (4.06) |
| | RUN 2 | 283.52 (2.85) | 287.19 (3.52) | 286.80 (4.87) | 286.77 (3.10) | 285.27 (3.44) | 279.41 (2.91) | 286.78 (5.07) | 283.99 (3.63) |
| | RUN 3 | 281.49 (3.19) | 289.28 (3.90) | 286.21 (2.87) | 282.84 (2.79) | 285.85 (3.52) | 283.45 (3.10) | 284.26 (3.76) | 285.21 (3.62) |
| Intra-protein Hydrogen bond | RUN 1 | 414.12 (9.60) | 415.49 (9.43) | 415.32 (9.13) | 410.81 (9.56) | 423.49 (9.31) | 412.27 (8.76) | 415.07 (10.02) | 412.96 (9.34) |
| | RUN 2 | 418.37 (9.30) | 417.49 (9.31) | 406.68 (9.03) | 413.72 (8.43) | 408.16 (9.64) | 425.07 (9.64) | 412.10 (9.54) | 411.73 (9.37) |
| | RUN 3 | 428.21 (9.29) | 414.81 (9.64) | 424.49 (8.77) | 410.57 (8.86) | 409.65 (9.34) | 412.58 (9.22) | 409.42 (8.68) | 414.33 (9.27) |
| Protein-water Hydrogen bond | RUN 1 | 1044.71 (20.68) | 1044.05 (18.84) | 1035.09 (20.07) | 1053.57 (20.76) | 1028.28 (19.74) | 1046.27 (18.61) | 1040.09 (20.77) | 1054.96 (19.13) |
| | RUN 2 | 1010.63 (19.54) | 1032.96 (20.09) | 1058.64 (18.89) | 1018.74 (20.10) | 1058.20 (18.13) | 1023.55 (17.41) | 1039.36 (18.98) | 1046.21 (19.00) |
| | RUN 3 | 999.93 (18.73) | 1042.23 (19.35) | 1014.66 (18.54) | 1031.08 (19.66) | 1051.06 (20.33) | 1037.72 (19.35) | 1054.36 (19.25) | 1035.89 (19.12) |
| RMSF (nm) | RUN 1 | 0.17 (0.30) | 0.17 (0.29) | 0.12 (0.17) | 0.11 (0.28) | 0.12 (0.10) | 0.14 (0.11) | 0.12 (0.08) | 0.14 (0.17) |
| | RUN 2 | 0.10 (0.11) | 0.16 (0.32) | 0.17 (0.32) | 0.13 (0.30) | 0.13 (0.17) | 0.09 (0.07) | 0.14 (0.17) | 0.13 (0.16) |
| | RUN 3 | 0.10 (0.08) | 0.15 (0.25) | 0.13 (0.23) | 0.13 (0.30) | 0.12 (0.08) | 0.10 (0.07) | 0.11 (0.08) | 0.12 (0.12) |
| N-C terminal distance (nm) | RUN 1 | 5.39 (0.55) | 5.78 (0.61) | 5.41 (0.61) | 5.50 (0.41) | 5.09 (0.33) | 4.80 (0.23) | 5.21 (0.28) | 5.86 (0.61) |
| | RUN 2 | 5.63 (0.57) | 5.12 (0.40) | 6.03 (0.62) | 5.62 (0.41) | 5.55 (0.74) | 5.06 (0.36) | 5.48 (0.59) | 5.69 (0.58) |
| | RUN 3 | 4.99 (0.59) | 5.37 (0.70) | 4.80 (0.47) | 6.05 (0.51) | 5.33 (0.21) | 4.81 (0.25) | 5.44 (0.18) | 5.08 (0.47) |

**Table S3. The average and standard deviation values for properties of the simulated protein from three simulation runs.**

| Protein Properties | Substrate state | | | | Intermediate state | | | |
|---|---|---|---|---|---|---|---|---|
| | **WT** | **H21A** | **K54A** | **E130A** | **WT** | **H21A** | **K54A** | **E130A** |
| **RMSD (nm)** | 0.34 (0.05) | 0.43 (0.09) | 0.49 (0.09) | 0.48 (0.09) | 0.42 (0.07) | 0.42 (0.06) | 0.49 (0.08) | 0.40 (0.06) |
| **Radius of gyration (nm)** | 2.70 (0.02) | 2.74 (0.03) | 2.77 (0.03) | 2.74 (0.02) | 2.77 (0.03) | 2.75 (0.02) | 2.79 (0.03) | 2.75 (0.03) |
| **Total SASA ($nm^2$)** | 234.41 (3.95) | 239.60 (4.30) | 239.56 (4.82) | 238.44 (3.99) | 240.62 (4.16) | 237.10 (3.45) | 240.92 (4.63) | 241.25 (4.55) |
| **Hydrophobic SASA ($nm^2$)** | 195.27 (1.77) | 195.13 (2.04) | 195.32 (1.88) | 196.02 (2.02) | 196.86 (1.95) | 194.61 (1.86) | 193.94 (2.01) | 195.00 (1.92) |
| **Hydrophilic SASA ($nm^2$)** | 284.12 (3.46) | 287.38 (3.58) | 286.19 (3.79) | 285.17 (3.35) | 284.17 (3.30) | 282.21 (3.03) | 285.26 (4.15) | 285.40 (3.77) |
| **Intra-protein Hydrogen bond** | 420.24 (9.40) | 415.93 (9.46) | 415.50 (8.98) | 411.70 (8.95) | 413.77 (9.43) | 416.64 (9.21) | 412.20 (9.41) | 413.01 (9.33) |
| **Protein-water Hydrogen bond** | 1018.42 (19.65) | 1039.75 (19.43) | 1036.13 (19.17) | 1034.46 (20.17) | 1045.85 (19.40) | 1035.85 (18.46) | 1044.60 (19.67) | 1045.69 (19.08) |
| **RMSF (nm)** | 0.12 (0.16) | 0.16 (0.29) | 0.14 (0.24) | 0.12 (0.29) | 0.12 (0.12) | 0.11 (0.08) | 0.12 (0.11) | 0.13 (0.15) |
| **N-C terminal distance (nm)** | 5.34 (0.57) | 5.42 (0.57) | 5.45 (0.56) | 5.72 (0.44) | 5.32 (0.43) | 4.89 (0.28) | 5.38 (0.35) | 5.54 (0.55) |

**Table S4. The properties of ligand in substrate (methionine) and intermediate (Met-AMP) state for each simulation run.**

| Ligand properties | | Substrate state (methionine) | | | | Intermediate state (Met-AMP) | | | |
|---|---|---|---|---|---|---|---|---|---|
| | | WT | H21A | K54A | E130A | WT | H21A | K54A | E130A |
| RMSD (nm) | RUN 1 | 2.80 (3.04) | 4.47 (3.21) | 0.71 (0.20) | 4.99 (3.02) | 0.33 (0.03) | 3.72 (2.67) | 3.03 (2.40) | 0.46 (0.14) |
| | RUN 2 | 0.44 (0.18) | 3.55 (3.17) | 6.03 (2.66) | 4.76 (3.24) | 3.67 (2.53) | 4.44 (2.78) | 3.76 (2.38) | 0.62 (0.24) |
| | RUN 3 | 0.47 (0.10) | 2.03 (2.81) | 3.67 (2.92) | 4.85 (3.11) | 4.52 (2.09) | 5.16 (2.00) | 4.24 (2.43) | 4.91 (2.19) |
| Radius of gyration (nm) | RUN 1 | 0.25 (0.01) | 0.25 (0.01) | 0.25 (0.01) | 0.25 (0.01) | 0.42 (0.01) | 0.43 (0.04) | 0.45 (0.04) | 0.41 (0.04) |
| | RUN 2 | 0.25 (0.01) | 0.25 (0.01) | 0.25 (0.01) | 0.25 (0.01) | 0.43 (0.04) | 0.43 (0.04) | 0.42 (0.04) | 0.45 (0.03) |
| | RUN 3 | 0.25 (0.01) | 0.25 (0.01) | 0.25 (0.01) | 0.25 (0.01) | 0.43 (0.04) | 0.42 (0.04) | 0.43 (0.04) | 0.42 (0.04) |
| Total SASA ($nm^2$) | RUN 1 | 3.34 (0.12) | 3.36 (0.13) | 3.37 (0.13) | 3.36 (0.12) | 6.91 (0.19) | 6.72 (0.37) | 6.86 (0.40) | 6.55 (0.43) |
| | RUN 2 | 3.34 (0.12) | 3.36 (0.11) | 3.36 (0.12) | 3.36 (0.12) | 6.72 (0.40) | 6.72 (0.41) | 6.68 (0.37) | 6.99 (0.30) |
| | RUN 3 | 3.37 (0.12) | 3.37 (0.12) | 3.36 (0.12) | 3.36 (0.12) | 6.74 (0.39) | 6.67 (0.43) | 6.74 (0.39) | 6.65 (0.39) |
| Protein-ligand Hydrogen bond | RUN 1 | 1.02 (1.43) | 0.65 (1.13) | 1.07 (0.43) | 0.21 (0.63) | 3.24 (0.54) | 0.38 (0.92) | 0.86 (1.23) | 3.46 (1.02) |
| | RUN 2 | 1.19 (0.61) | 0.51 (1.19) | 0.22 (0.69) | 0.21 (0.64) | 0.68 (1.22) | 0.11 (0.38) | 0.68 (1.01) | 1.35 (0.89) |
| | RUN 3 | 2.72 (0.75) | 1.40 (1.41) | 0.96 (1.24) | 0.41 (0.82) | 0.67 (0.83) | 0.44 (1.00) | 0.46 (0.94) | 0.13 (0.47) |
| Ligand-water Hydrogen bond | RUN 1 | 7.51 (1.81) | 7.78 (1.89) | 6.80 (1.21) | 8.30 (1.28) | 13.37 (1.73) | 16.35 (2.52) | 15.30 (3.18) | 10.67 (2.29) |
| | RUN 2 | 6.03 (1.23) | 8.00 (1.53) | 8.26 (1.30) | 8.28 (1.33) | 16.19 (2.66) | 16.85 (2.23) | 16.05 (2.33) | 14.71 (1.99) |
| | RUN 3 | 5.34 (0.86) | 6.59 (2.21) | 7.66 (1.58) | 8.06 (1.33) | 15.87 (2.34) | 16.32 (2.53) | 16.26 (2.51) | 16.80 (2.33) |
| RMSF (nm) | RUN 1 | 0.11 (0.05) | 0.11 (0.04) | 0.11 (0.04) | 0.11 (0.04) | 0.06 (0.03) | 0.24 (0.08) | 0.26 (0.09) | 0.14 (0.05) |
| | RUN 2 | 0.10 (0.05) | 0.12 (0.04) | 0.10 (0.05) | 0.11 (0.05) | 0.24 (0.08) | 0.23 (0.09) | 0.24 (0.09) | 0.20 (0.07) |
| | RUN 3 | 0.11 (0.04) | 0.10 (0.04) | 0.11 (0.05) | 0.10 (0.05) | 0.23 (0.09) | 0.27 (0.09) | 0.27 (0.09) | 0.26 (0.09) |

**Table S5. The average and standard deviation values for the properties of the simulated ligand molecule (In substrate state - methionine and intermediate state - Met-AMP) from three simulation runs.**

| Ligand properties | Substrate state (Methionine) | | | | Intermediate state (Met-AMP) | | | |
|---|---|---|---|---|---|---|---|---|
| | WT | H21A | K54A | E130A | WT | H21A | K54A | E130A |
| RMSD (nm) | 1.24 (1.11) | 3.35 (3.06) | 3.47 (1.93) | 4.86 (3.12) | 2.84 (1.55) | 4.44 (2.48) | 3.67 (2.40) | 2.00 (0.86) |
| Radius of gyration (nm) | 0.25 (0.01) | 0.25 (0.01) | 0.25 (0.01) | 0.25 (0.01) | 0.43 (0.03) | 0.43 (0.04) | 0.43 (0.04) | 0.43 (0.04) |
| Total SASA ($nm^2$) | 3.35 (0.12) | 3.36 (0.12) | 3.37 (0.12) | 3.36 (0.12) | 6.79 (0.33) | 6.70 (0.40) | 6.76 (0.39) | 6.73 (0.37) |
| Protein-ligand Hydrogen bonds | 1.64 (0.93) | 0.85 (1.24) | 0.75 (0.79) | 0.28 (0.70) | 1.53 (0.86) | 0.31 (0.77) | 0.67 (1.06) | 1.65 (0.80) |
| Ligand-water Hydrogen bonds | 6.29 (1.30) | 7.46 (1.88) | 7.57 (1.36) | 8.21 (1.32) | 15.14 (2.24) | 16.51 (2.43) | 15.87 (2.67) | 14.06 (2.20) |
| RMSF (nm) | 0.10 (0.05) | 0.11 (0.04) | 0.10 (0.05) | 0.11 (0.05) | 0.17 (0.07) | 0.25 (0.09) | 0.26 (0.09) | 0.20 (0.07) |

**Table S6. The properties of ATP molecule in substrate state for each simulation run.**

| ATP properties | | Substrate state | | | | Intermediate state | | | |
|---|---|---|---|---|---|---|---|---|---|
| | | WT | H21A | K54A | E130A | WT | H21A | K54A | E130A |
| RMSD (nm) | RUN 1 | 2.42 (1.99) | 4.05 (2.38) | 0.34 (0.07) | 2.91 (2.70) | - | - | - | - |
| | RUN 2 | 0.23 (0.05) | 0.33 (0.95) | 4.56 (1.60) | 1.06 (0.62) | - | - | - | - |
| | RUN 3 | 0.58 (0.13) | 0.37 (0.10) | 3.17 (1.56) | 1.36 (1.44) | - | - | - | - |
| Radius of gyration (nm) | RUN 1 | 0.45 (0.02) | 0.43 (0.02) | 0.41 (0.01) | 0.43 (0.02) | - | - | - | - |
| | RUN 2 | 0.45 (0.01) | 0.43 (0.01) | 0.45 (0.02) | 0.43 (0.03) | - | - | - | - |
| | RUN 3 | 0.42 (0.02) | 0.43 (0.01) | 0.44 (0.02) | 0.43 (0.02) | - | - | - | - |
| Total SASA ($nm^2$) | RUN 1 | 6.46 (0.23) | 6.42 (0.20) | 6.37 (0.22) | 6.36 (0.22) | - | - | - | - |
| | RUN 2 | 6.56 (0.17) | 6.36 (0.18) | 6.42 (0.21) | 6.39 (0.24) | - | - | - | - |
| | RUN 3 | 6.28 (0.19) | 6.37 (0.19) | 6.42 (0.23) | 6.31 (0.21) | - | - | - | - |
| Protein-ATP Hydrogen bond | RUN 1 | 4.93 (1.27) | 4.53 (1.50) | 5.01 (1.04) | 1.31 (2.07) | - | - | - | - |
| | RUN 2 | 10.28 (1.05) | 6.18 (1.73) | 4.93 (1.14) | 4.57 (1.36) | - | - | - | - |
| | RUN 3 | 7.80 (0.90) | 5.27 (1.58) | 5.90 (0.75) | 6.15 (1.31) | - | - | - | - |
| ATP-water Hydrogen bond | RUN 1 | 21.44 (2.84) | 20.47 (2.89) | 18.13 (2.27) | 23.19 (3.25) | - | - | - | - |
| | RUN 2 | 18.77 (1.72) | 17.30 (2.58) | 18.07 (2.65) | 21.55 (2.91) | - | - | - | - |
| | RUN 3 | 15.24 (1.84) | 17.58 (2.79) | 19.12 (2.62) | 18.03 (2.39) | - | - | - | - |
| RMSF (nm) | RUN 1 | 0.15 (0.06) | 0.13 (0.04) | 0.06 (0.03) | 0.13 (0.05) | - | - | - | - |
| | RUN 2 | 0.04 (0.03) | 0.07 (0.03) | 0.07 (0.03) | 0.18 (0.06) | - | - | - | - |
| | RUN 3 | 0.05 (0.03) | 0.09 (0.04) | 0.15 (0.05) | 0.13 (0.05) | - | - | - | - |

**Table S7. The average and standard deviation values for the properties of the ATP molecule in substrate state from three simulation runs.**

| ATP properties | Substrate state | | | | Intermediate state | | | |
|---|---|---|---|---|---|---|---|---|
| | WT | H21A | K54A | E130A | WT | H21A | K54A | E130A |
| RMSD (nm) | 1.08 (0.72) | 1.58 (1.14) | 2.69 (1.08) | 1.78 (1.58) | - | - | - | - |
| Radius of gyration (nm) | 0.44 (0.02) | 0.43 (0.02) | 0.43 (0.02) | 0.43 (0.02) | - | - | - | - |
| Total SASA ($nm^2$) | 6.43 (0.20) | 6.38 (0.19) | 6.40 (0.22) | 6.35 (0.22) | - | - | - | - |
| Protein-ATP Hydrogen bonds | 7.67 (1.07) | 5.33 (1.61) | 5.28 (0.98) | 4.01 (1.58) | - | - | - | - |
| ATP-water Hydrogen bonds | 18.48 (2.13) | 18.45 (2.75) | 18.44 (2.51) | 20.93 (2.85) | - | - | - | - |
| RMSF (nm) | 0.08 (0.04) | 0.10 (0.04) | 0.09 (0.04) | 0.15 (0.05) | - | - | - | - |

**Table S8. Volume of active site in representative structures obtained using CavitOmiX plugin of Pymol.**

| MetRS protein model | | Substrate state volume ($nm^3$) | | Intermediate state volume ($nm^3$) | |
|---|---|---|---|---|---|
| WT | Representative structure 1 | 4.575 | 9.133 | 3.031 | 9.202 |
| | Representative structure 2 | 1.865 | | 1.836 | |
| | Representative structure 3 | 2.693 | | 4.335 | |
| H21A | Representative structure 1 | 3.531 | 11.676 | 3.591 | 9.230 |
| | Representative structure 2 | 3.726 | | 3.412 | |
| | Representative structure 3 | 4.419 | | 2.227 | |
| K54A | Representative structure 1 | 1.290 | 5.843 | 2.717 | 9.637 |
| | Representative structure 2 | 3.102 | | 2.432 | |
| | Representative structure 3 | 1.451 | | 4.488 | |
| E130A | Representative structure 1 | 2.922 | 9.732 | 2.808 | 9.979 |
| | Representative structure 2 | 3.924 | | 3.822 | |
| | Representative structure 3 | 2.886 | | 3.349 | |

**Table S9. The MetRS enzyme activity ($K_m$ value) in different organisms.**

| Name of organism | Description | $K_m$ value for substrate molecules (µM) | | |
|---|---|---|---|---|
| | | $K_m$ (Methionine) | $K_m$ (ATP) | $K_m$ ($tRNA^{met}$) |
| *Mycobacterium tuberculosis* | Mycobacteria | **50** (Wang *et al.*)[17] | **4282** (Wang et al.)[17] | **3.2** (Kim et al.)[18] |
| *Saccharomyces cerevisiae* (cytoplasmic) | Yeast (Fungus) | **5** (Schwob *et al.*)[19] | | **1.0** (Schwob et al.) |
| *Saccharomyces cerevisiae* (mitochondrial) | Yeast (Fungus) | **6.8** (Schwob *et al.*)[19] | | **1.1** (Schwob *et al.*) |
| *Bacillus stearothermophilus* | Gram-positive, thermophilic bacterium | **6** (Kalogerakos *et al.*)[20], **8** (Schmitt *et al.*)[21] | **12** (Kalogerakos *et al.*)[20], **11** (Schmitt *et al.*)[21] | |
| *Homo sapiens* (mitochondrial) | | **18** (Spencer et al.)[22], **20** (Green et al.)[23] | **85** (Spencer *et al.*)[22], (Green *et al.*)[23] | **2.1** (Spencer *et al.)*[22] |
| *Escherichia coli* | Gram-negative bacterium | **21** (Ghosh *et al.*)[24], **20** **(**Crepin *et al.*)[25], (Jakubowski *et al.*)[26] | **528** (Ghosh *et al.*)[24] | **1.2** (Ghosh *et al.*)[24] |
| *Thermus thermophilus* | Gram-negative thermophilic bacterium | **27** (Kohda *et al.*)[27] **18** (Nureki *et al.)*[28] | **180** (Kohda *et al.*)[27] **1600** (Kohda *et al.*)[27], **180** (Nureki *et al.*)[28], **1500** (Nureki *et al.)*[28] | **1.4** (Kohda *et al.*)[27], (Nureki *et al.*)[28] |
| *Streptococcus pneumoniae (MetRS1)* | Gram-positive bacterium | **53** (Green *et al.)*[23] | **514** (Green *et al.*)[23] | |
| *Staphylococcus aureus* | Gram-positive bacterium | **100** (Green *et al.*)[23] | **500** (Green *et al.*)[23] | |
| *Haemophilus influenzae* | Gram-negative bacteria | **250** (Green *et al.*)[23] | **1200** (Green *et al.*)[23] | |

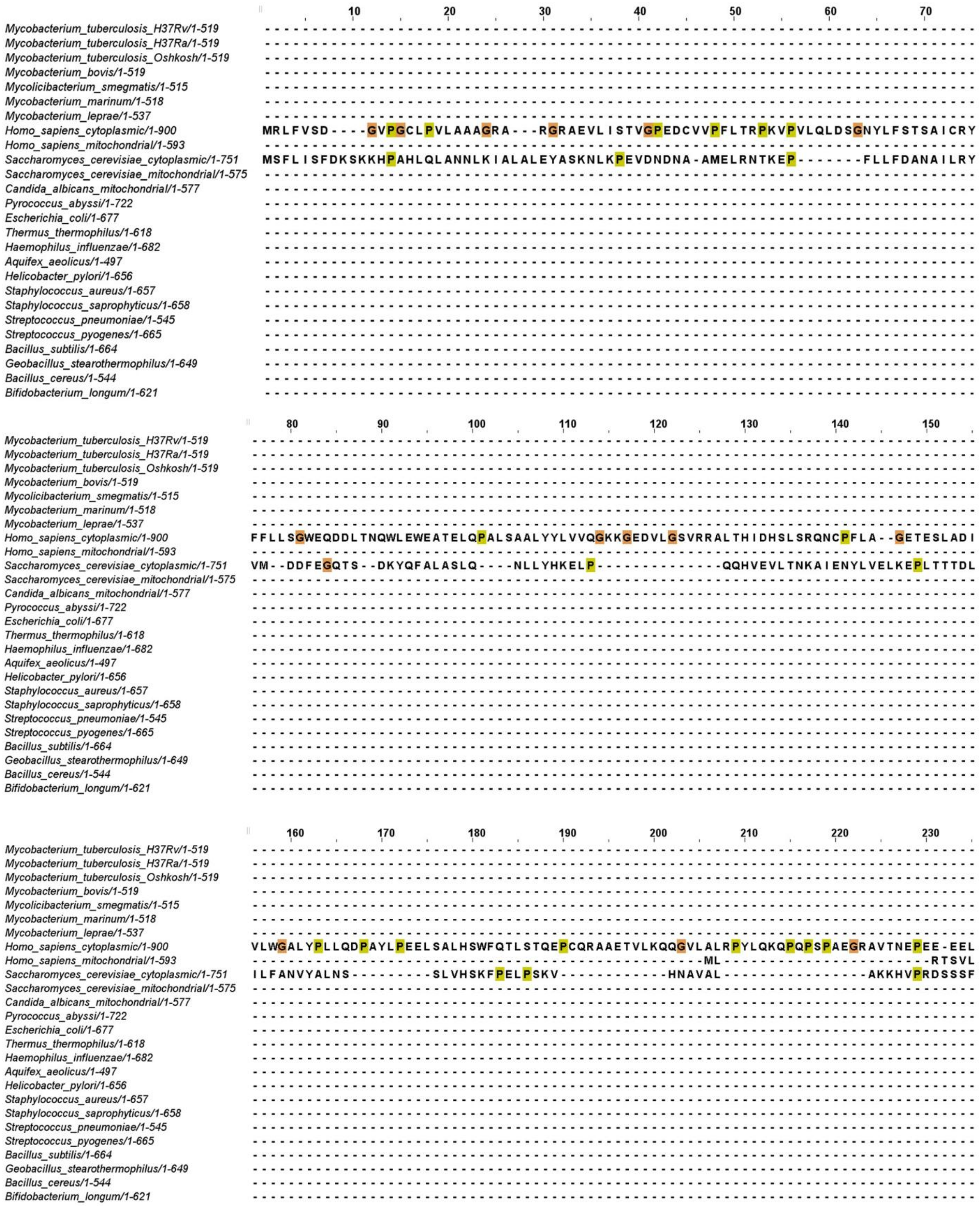
10 20 30 40 50 60 70
Mycobacterium_tuberculosis_H37Rv/1-519
Mycobacterium_tuberculosis_H37Ra/1-519
Mycobacterium_tuberculosis_Oshkosh/1-519
Mycobacterium_bovis/1-519
Mycolicibacterium_smegmatis/1-515
Mycobacterium_marinum/1-518
Mycobacterium_leprae/1-537
Homo_sapiens_cytoplasmic/1-900
Homo_sapiens_mitochondrial/1-593
Saccharomyces_cerevisiae_cytoplasmic/1-751
Saccharomyces_cerevisiae_mitochondrial/1-575
Candida_albicans_mitochondrial/1-577
Pyrococcus_abyssi/1-722
Escherichia_coli/1-677
Thermus_thermophilus/1-618
Haemophilus_influenzae/1-682
Aquifex_aeolicus/1-497
Helicobacter_pylori/1-656
Staphylococcus_aureus/1-657
Staphylococcus_saprophyticus/1-658
Streptococcus_pneumoniae/1-545
Streptococcus_pyogenes/1-665
Bacillus_subtilis/1-664
Geobacillus_stearothermophilus/1-649
Bacillus_cereus/1-544
Bifidobacterium_longum/1-621
MRLFVSD----GVPGCLPVLAAAGRA---RGRAEVLISTVGPEDCVVPFLTRPKVPVLQLDSGNYLFSTSAICRY
MSFLISFDKSKKHPAHLQLANNLKIALALEYASKNLKPEVDNDNA-AMELRNTKEP------FLLFDANAILRY
80 90 100 110 120 130 140 150
FFLLSGWEQDDLTNQWLEWEATELQPALSAALYYLVVQGKKGEDVLGSVRRALTHIDHSLSRQNCPFLA--GETESLADI
VM--DDFEGQTS--DKYQFALASLQ---NLLYHKELP-------------QQHVEVLTNKAIENYLVELKEPLTTTDL
160 170 180 190 200 210 220 230
VLWGALYPLLQDPAYLPEELSALHSWFQTLSTQEPCQRAAETVLKQQGVLALRPYLQKQPQPSPAEGRAVTNEPEE-EEL
ML RTSVL
ILFANVYALNS---------SLVHSKFPELPSKV------------HNAVAL---------------AKKHVPRDSSSF

**Continue…...**

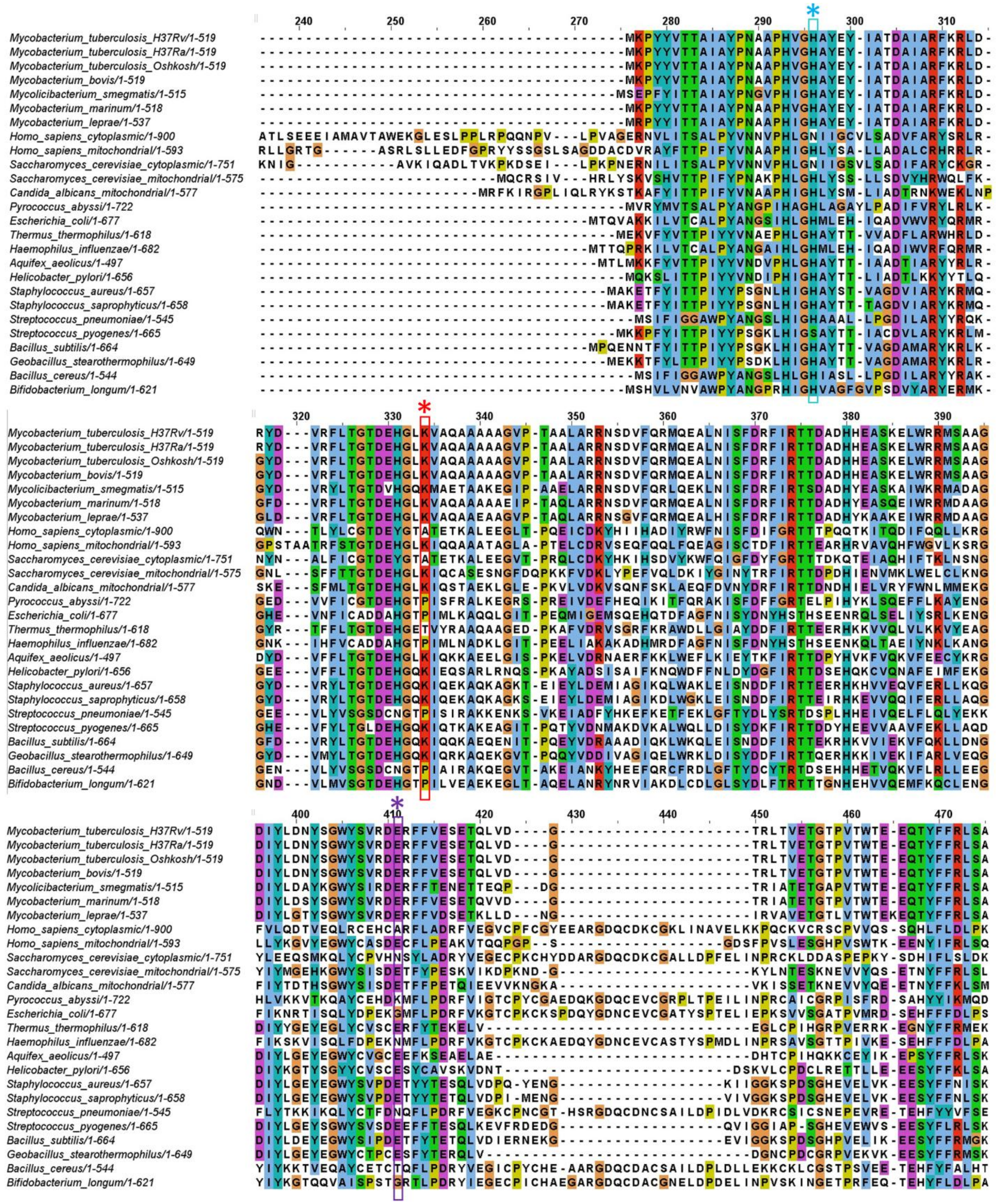
Mycobacterium_tuberculosis_H37Rv/1-519
Mycobacterium_tuberculosis_H37Ra/1-519
Mycobacterium_tuberculosis_Oshkosh/1-519
Mycobacterium_bovis/1-519
Mycolicibacterium_smegmatis/1-515
Mycobacterium_marinum/1-518
Mycobacterium_leprae/1-537
Homo_sapiens_cytoplasmic/1-900
Homo_sapiens_mitochondrial/1-593
Saccharomyces_cerevisiae_cytoplasmic/1-751
Saccharomyces_cerevisiae_mitochondrial/1-575
Candida_albicans_mitochondrial/1-577
Pyrococcus_abyssi/1-722
Escherichia_coli/1-677
Thermus_thermophilus/1-618
Haemophilus_influenzae/1-682
Aquifex_aeolicus/1-497
Helicobacter_pylori/1-656
Staphylococcus_aureus/1-657
Staphylococcus_saprophyticus/1-658
Streptococcus_pneumoniae/1-545
Streptococcus_pyogenes/1-665
Bacillus_subtilis/1-664
Geobacillus_stearothermophilus/1-649
Bacillus_cereus/1-544
Bifidobacterium_longum/1-621

**Continue…...**

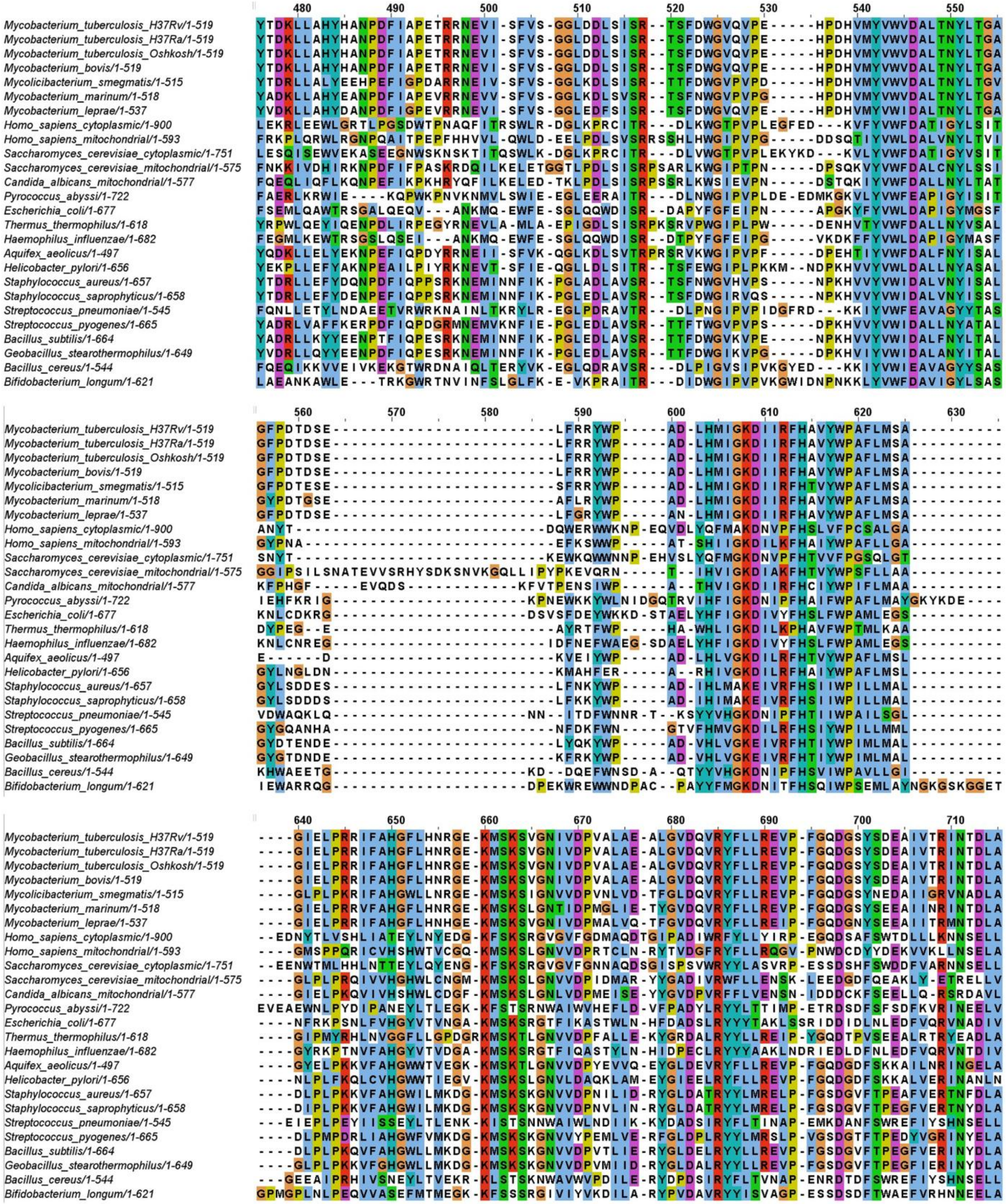
480 490 500 510 520 530 540 550
560 570 580 590 600 610 620 630
640 650 660 670 680 690 700 710
Mycobacterium_tuberculosis_H37Rv/1-519
Mycobacterium_tuberculosis_H37Ra/1-519
Mycobacterium_tuberculosis_Oshkosh/1-519
Mycobacterium_bovis/1-519
Mycolicibacterium_smegmatis/1-515
Mycobacterium_marinum/1-518
Mycobacterium_leprae/1-537
Homo_sapiens_cytoplasmic/1-900
Homo_sapiens_mitochondrial/1-593
Saccharomyces_cerevisiae_cytoplasmic/1-751
Saccharomyces_cerevisiae_mitochondrial/1-575
Candida_albicans_mitochondrial/1-577
Pyrococcus_abyssi/1-722
Escherichia_coli/1-677
Thermus_thermophilus/1-618
Haemophilus_influenzae/1-682
Aquifex_aeolicus/1-497
Helicobacter_pylori/1-656
Staphylococcus_aureus/1-657
Staphylococcus_saprophyticus/1-658
Streptococcus_pneumoniae/1-545
Streptococcus_pyogenes/1-665
Bacillus_subtilis/1-664
Geobacillus_stearothermophilus/1-649
Bacillus_cereus/1-544
Bifidobacterium_longum/1-621

**Continue…...**

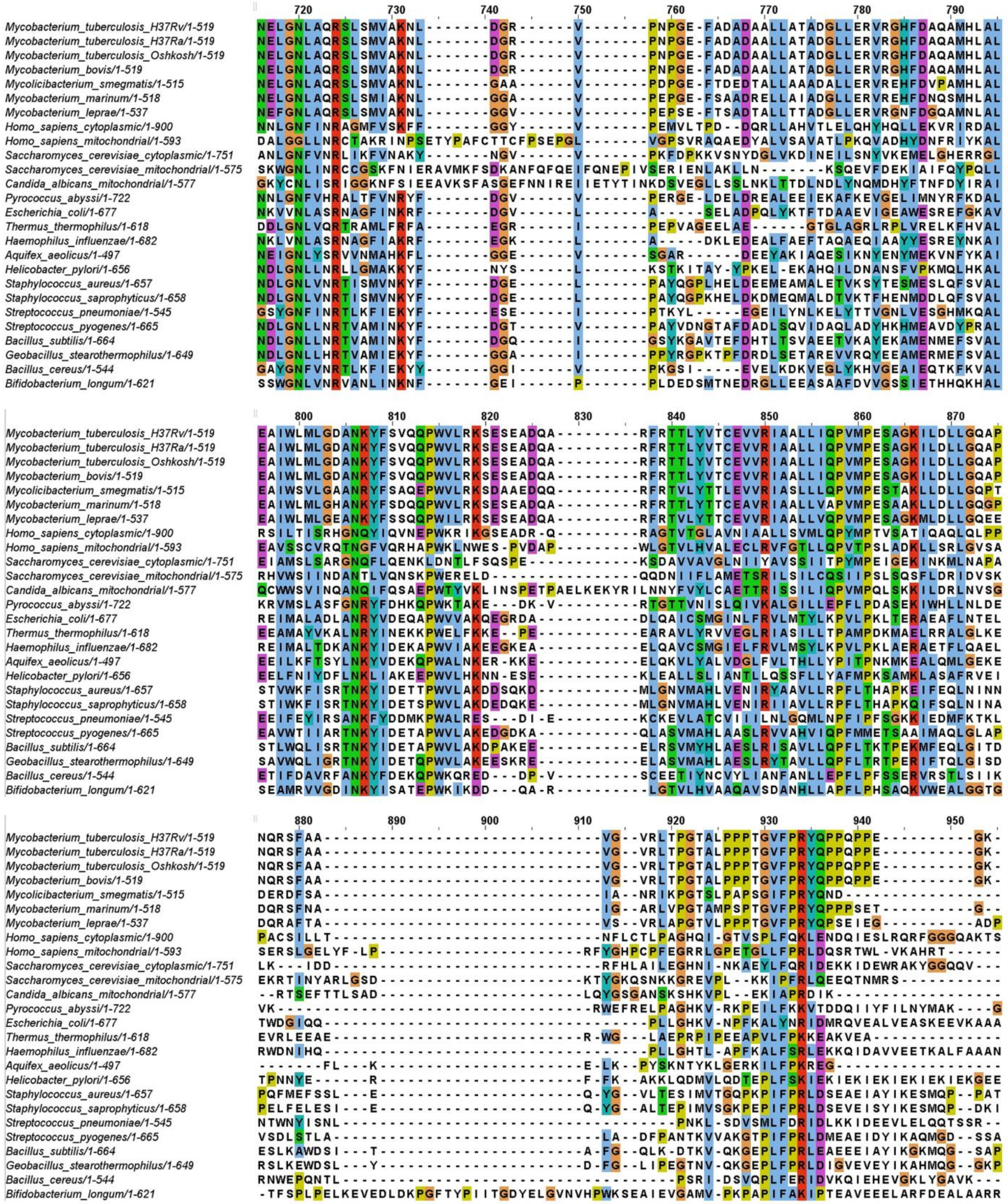

720 730 740 750 760 770 780 790
800 810 820 830 840 850 860 870
880 890 900 910 920 930 940 950
Mycobacterium_tuberculosis_H37Rv/1-519
Mycobacterium_tuberculosis_H37Ra/1-519
Mycobacterium_tuberculosis_Oshkosh/1-519
Mycobacterium_bovis/1-519
Mycolicibacterium_smegmatis/1-515
Mycobacterium_marinum/1-518
Mycobacterium_leprae/1-537
Homo_sapiens_cytoplasmic/1-900
Homo_sapiens_mitochondrial/1-593
Saccharomyces_cerevisiae_cytoplasmic/1-751
Saccharomyces_cerevisiae_mitochondrial/1-575
Candida_albicans_mitochondrial/1-577
Pyrococcus_abyssi/1-722
Escherichia_coli/1-677
Thermus_thermophilus/1-618
Haemophilus_influenzae/1-682
Aquifex_aeolicus/1-497
Helicobacter_pylori/1-656
Staphylococcus_aureus/1-657
Staphylococcus_saprophyticus/1-658
Streptococcus_pneumoniae/1-545
Streptococcus_pyogenes/1-665
Bacillus_subtilis/1-664
Geobacillus_stearothermophilus/1-649
Bacillus_cereus/1-544
Bifidobacterium_longum/1-621


**Continue…...**

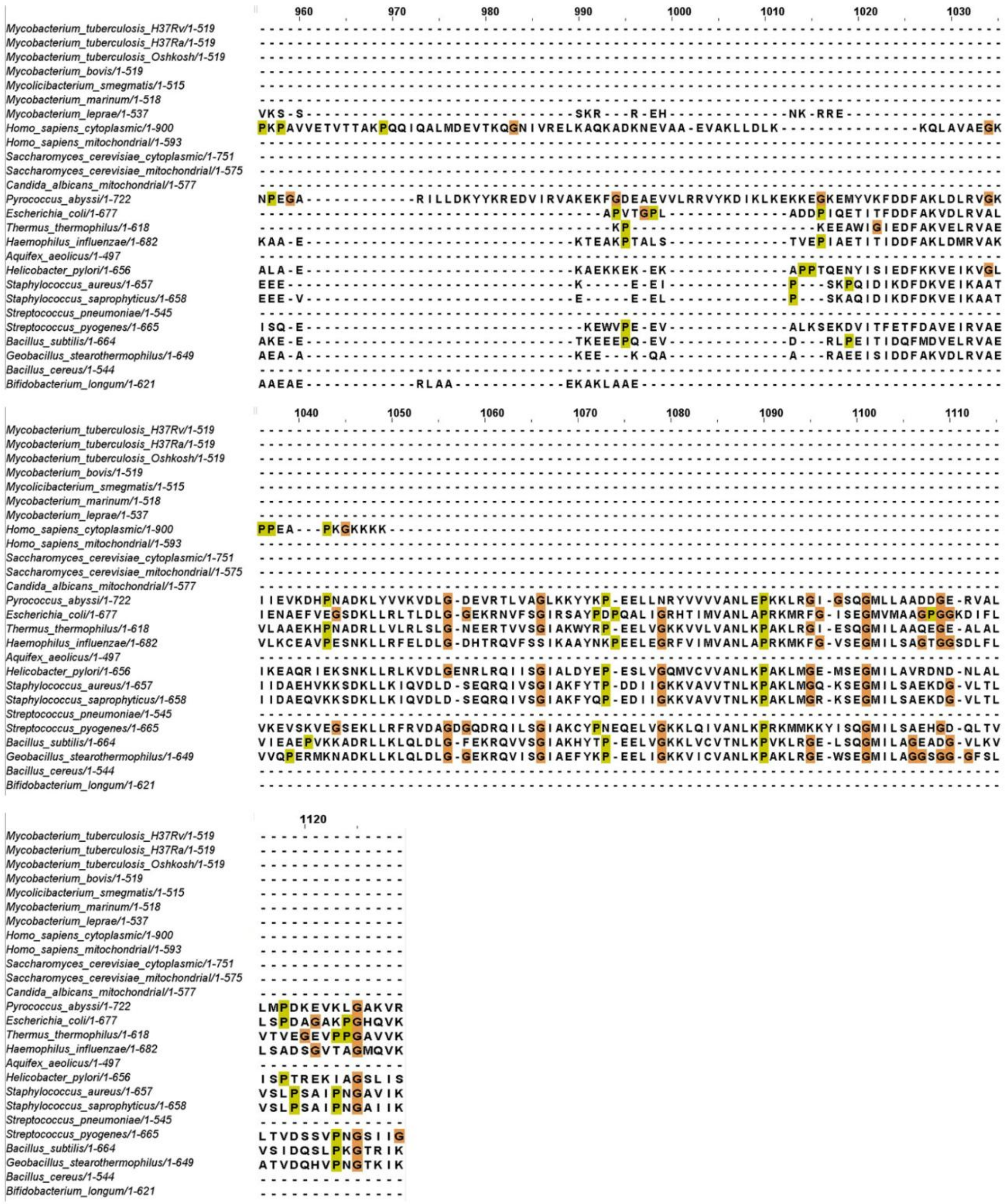

**Figure S1. Multiple sequence alignment of MetRS protein sequence from 26 organisms including bacteria from mycobacterium genus, gram-positive and gram-negative bacteria, eukaryotes, archaeon etc.** The residues which are mutated in this study (H21, K54 and E130) of *Mtb* MetRS are observed to be conserved in almost all the MetRS proteins of other organisms (highlighted in a square box with * sign). Each amino acid is given a unique color in the alignment.

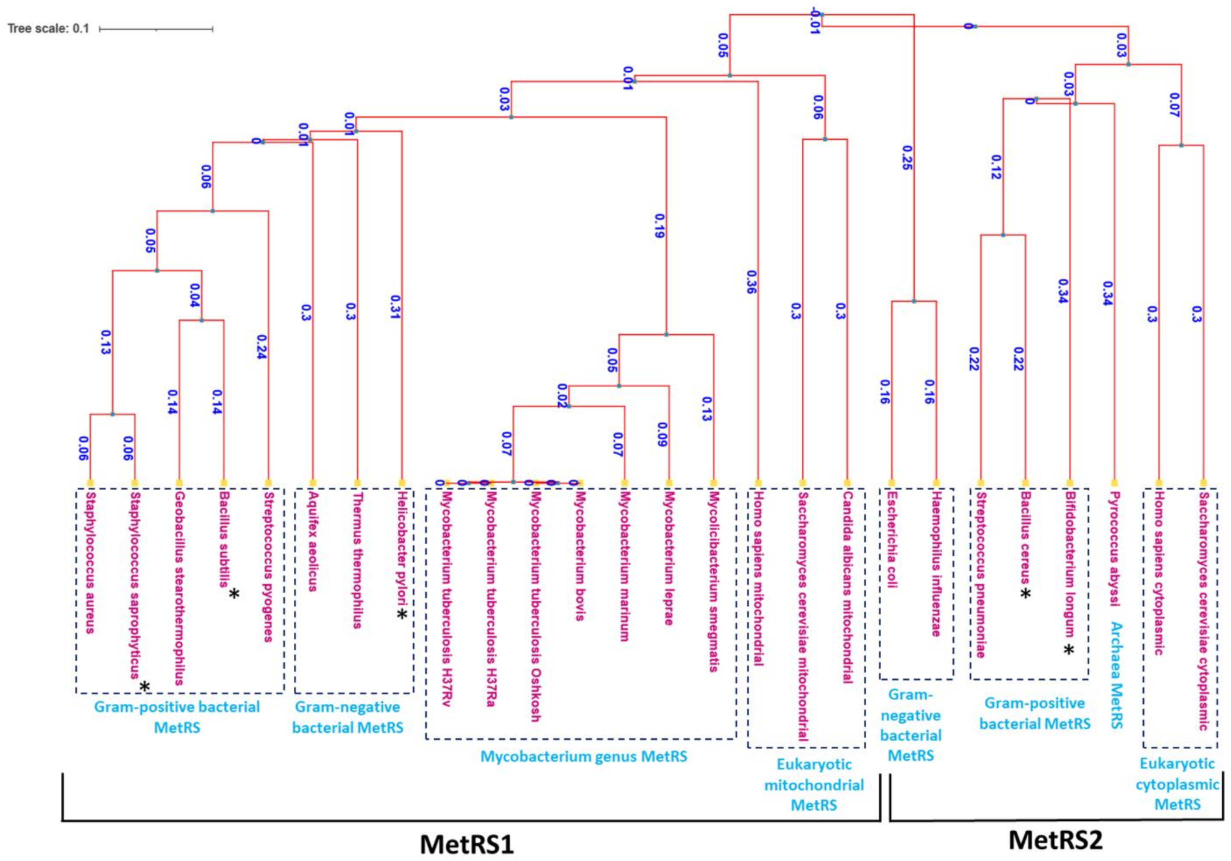


**Figure S2. A rectangular phylogenetic tree depicting evolutionary relationships among various species with values on the branches of the dendrogram represent the distance between the organisms at the end of each branch (dark blue color)**. Various clusters formed by a group of species are encircled into dashed box. Shorter branch length represents close evolutionary relationship between MetRS of two organisms or species and vice versa. We have placed this MetRS sequences into MetRS1 and MetRS2 category based on sequence identity with other representative of that class for which category is confirmed. The MetRS sequences for which category is not confirmed are marked with asterisk sign (*).

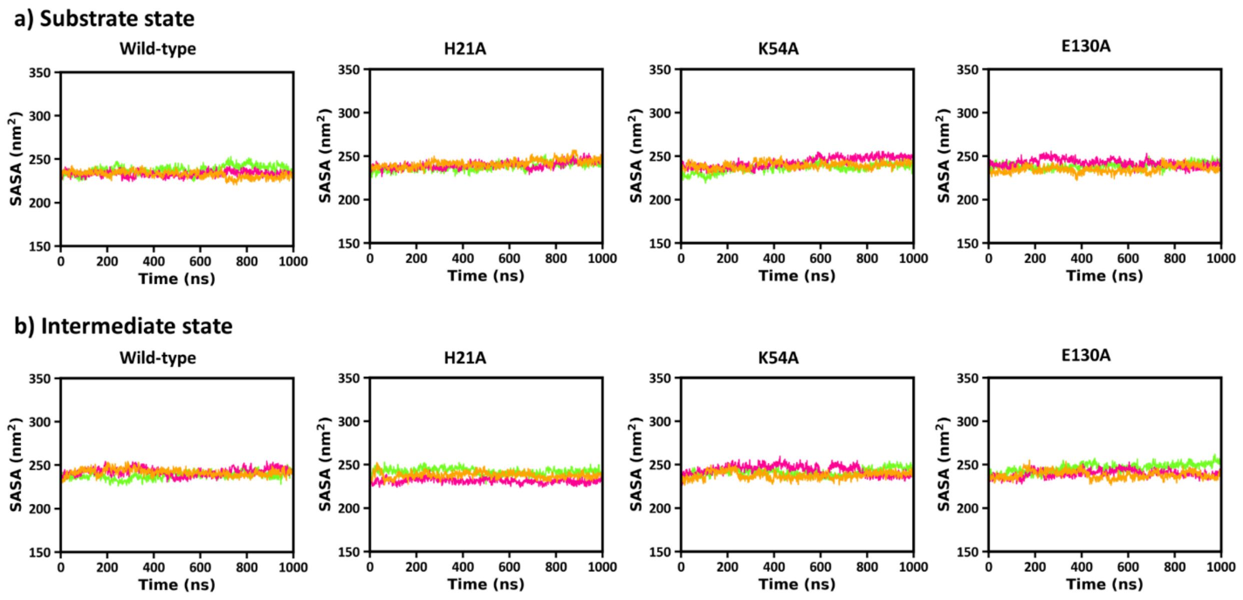


**Figure S3. Total SASA of protein.** SASA values for **(a)** substrate state and **(b)** intermediate state in wild-type and mutant models (H21A, K54A and E130A).

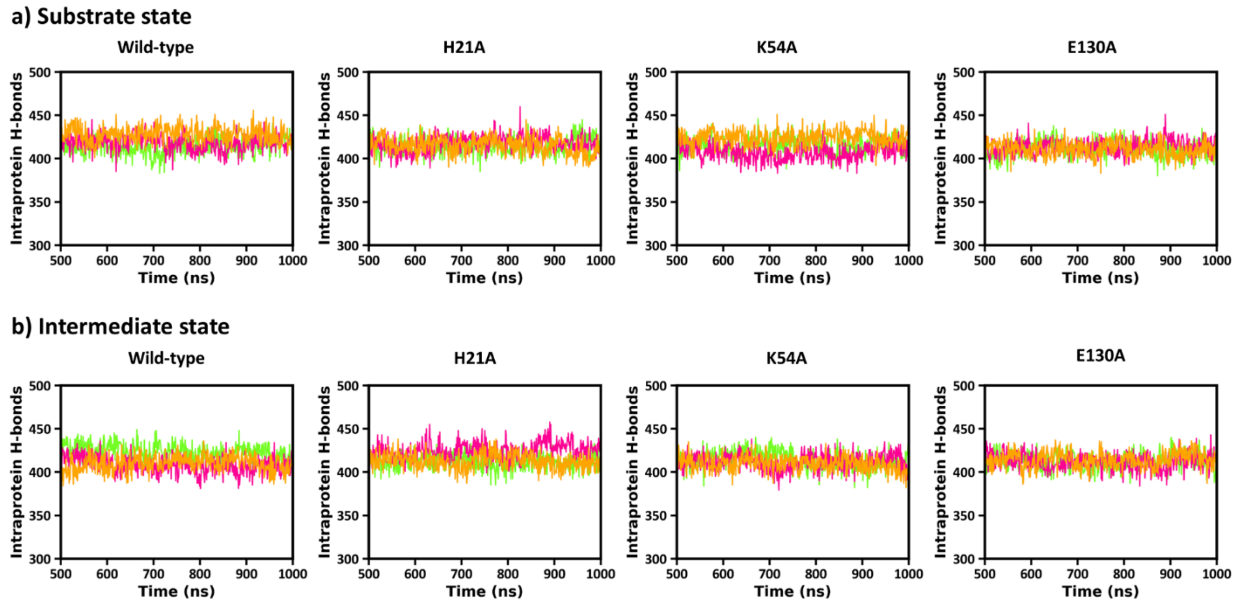


**Figure S4. Intraprotein hydrogen bonds.** Intramolecular hydrogen bonds of protein for **(a)** substrate state and **(b)** intermediate state in wild-type and mutant models (H21A, K54A and E130A).

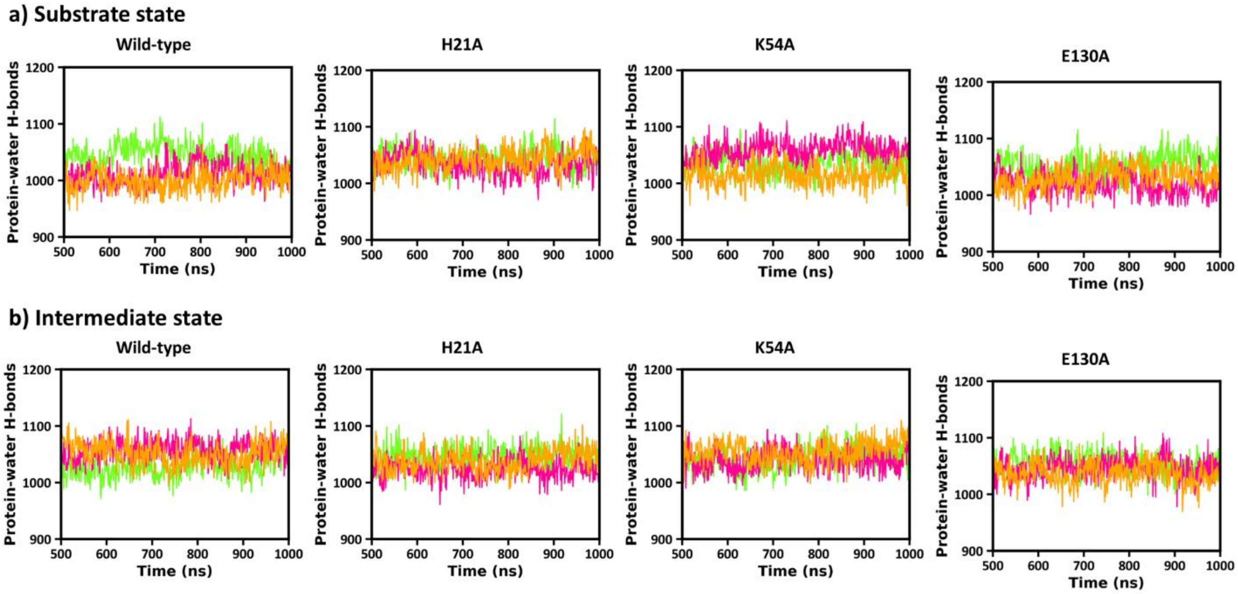


**Figure S5. Protein-water hydrogen bond interactions.** Hydrogen bond count for two protein states **(a)** substrate state and **(b)** intermediate state.

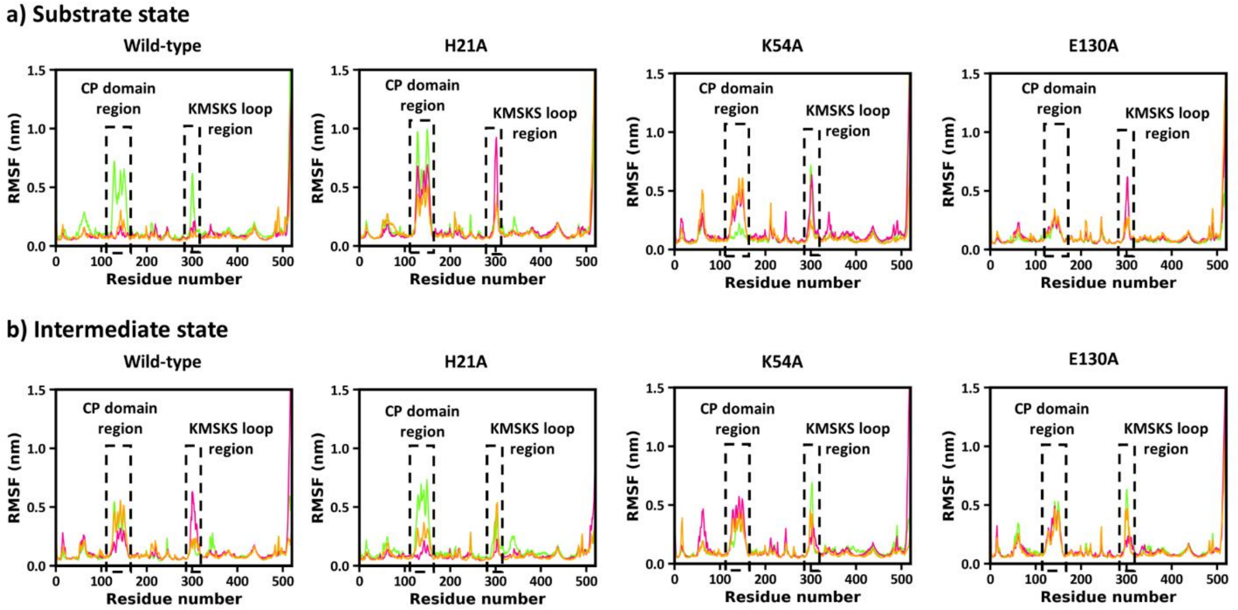


**Figure S6. RMSF value of each protein residue. (a)** Substrate state and **(b)** intermediate state MetRS protein in wild type and three mutant models (H21A, K54A and E130A).

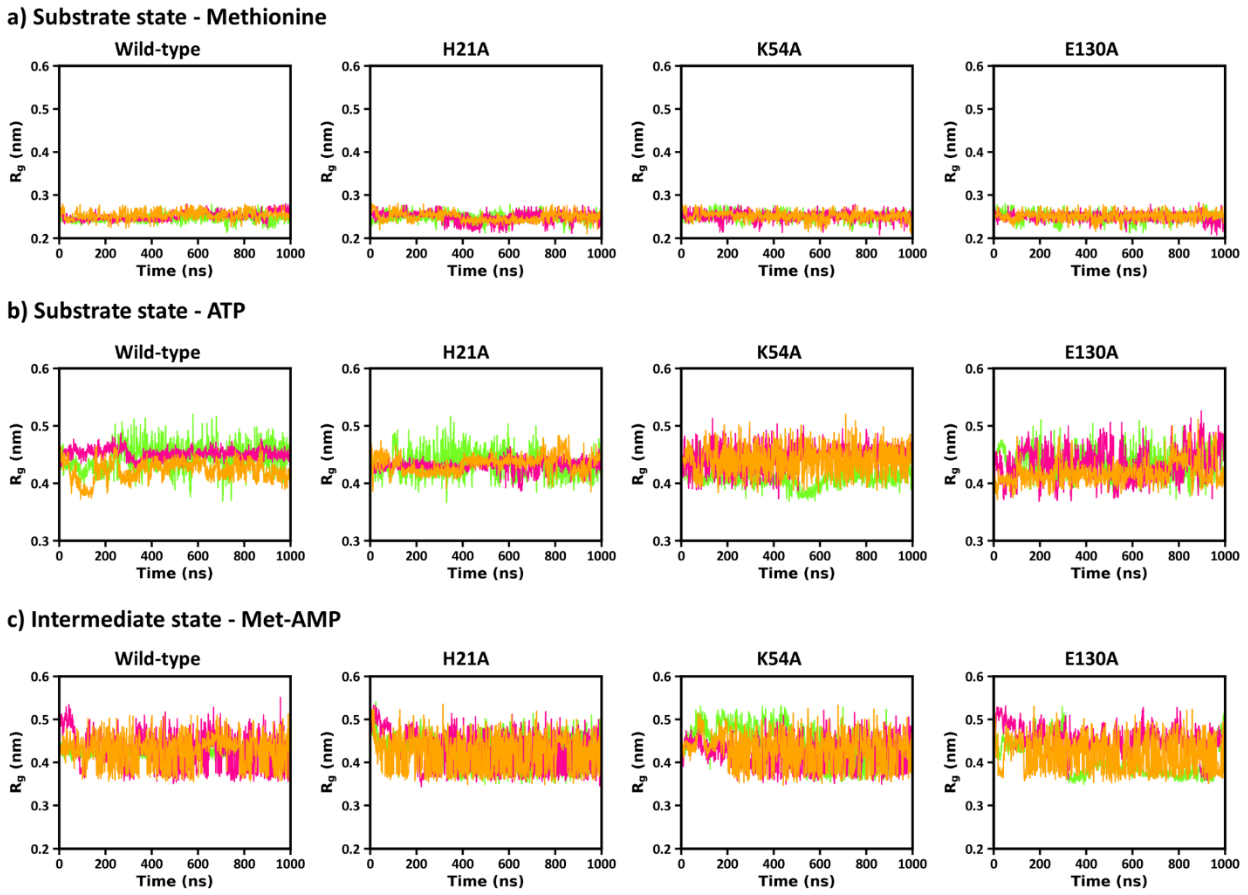


**Figure S7. $R_g$ of a ligand (Substrate state - Methionine and ATP, Intermediate state - Met-AMP).** $R_g$ value over time for different protein variants: Wild-type, H21A, K54A, and E130A in **(a-b)** substrate state and **(c)** intermediate state.

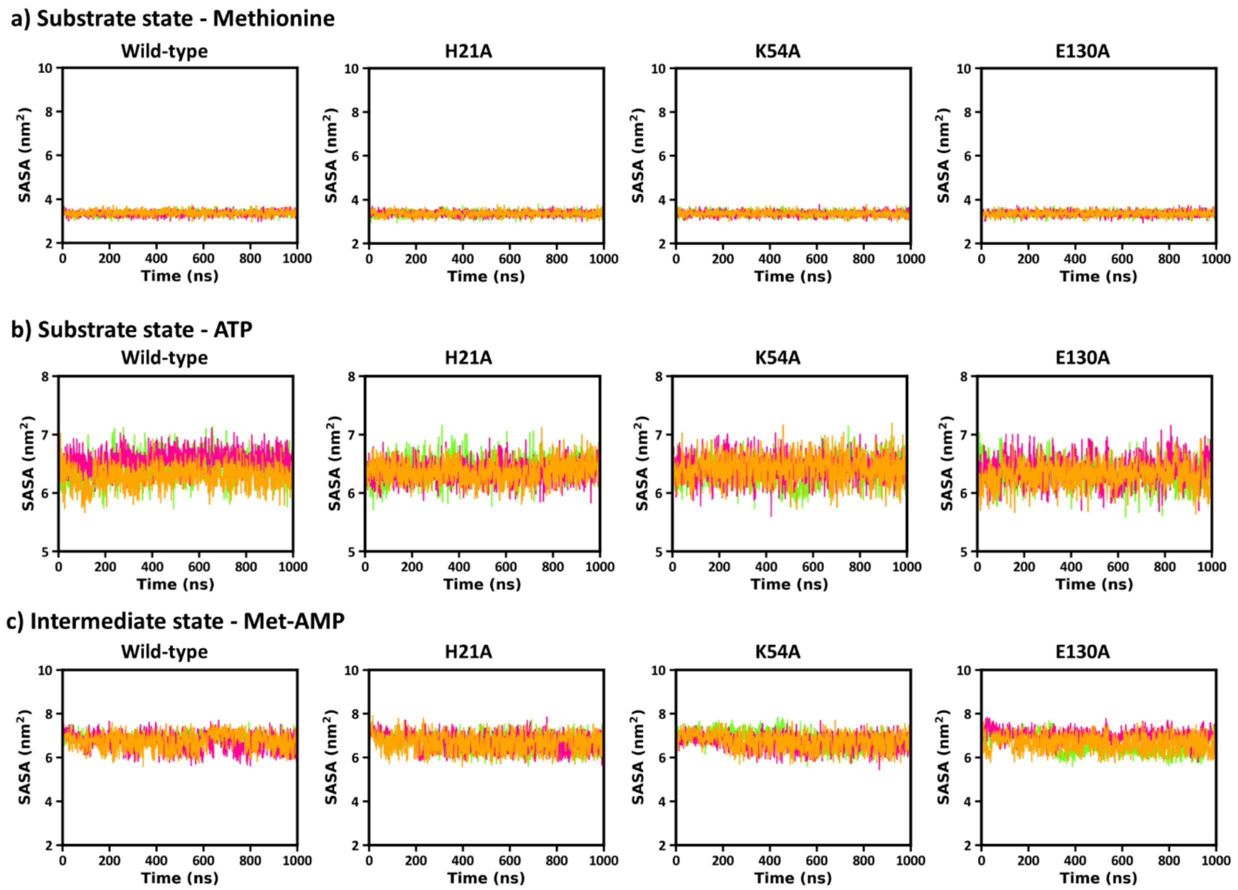


**Figure S8. SASA values of the ligand**. **(a-b)** Substrate state - methionine and ATP, **(c)** intermediate state - Met-AMP for the wild-type enzyme and three mutants (H21A, K54A, and E130A).

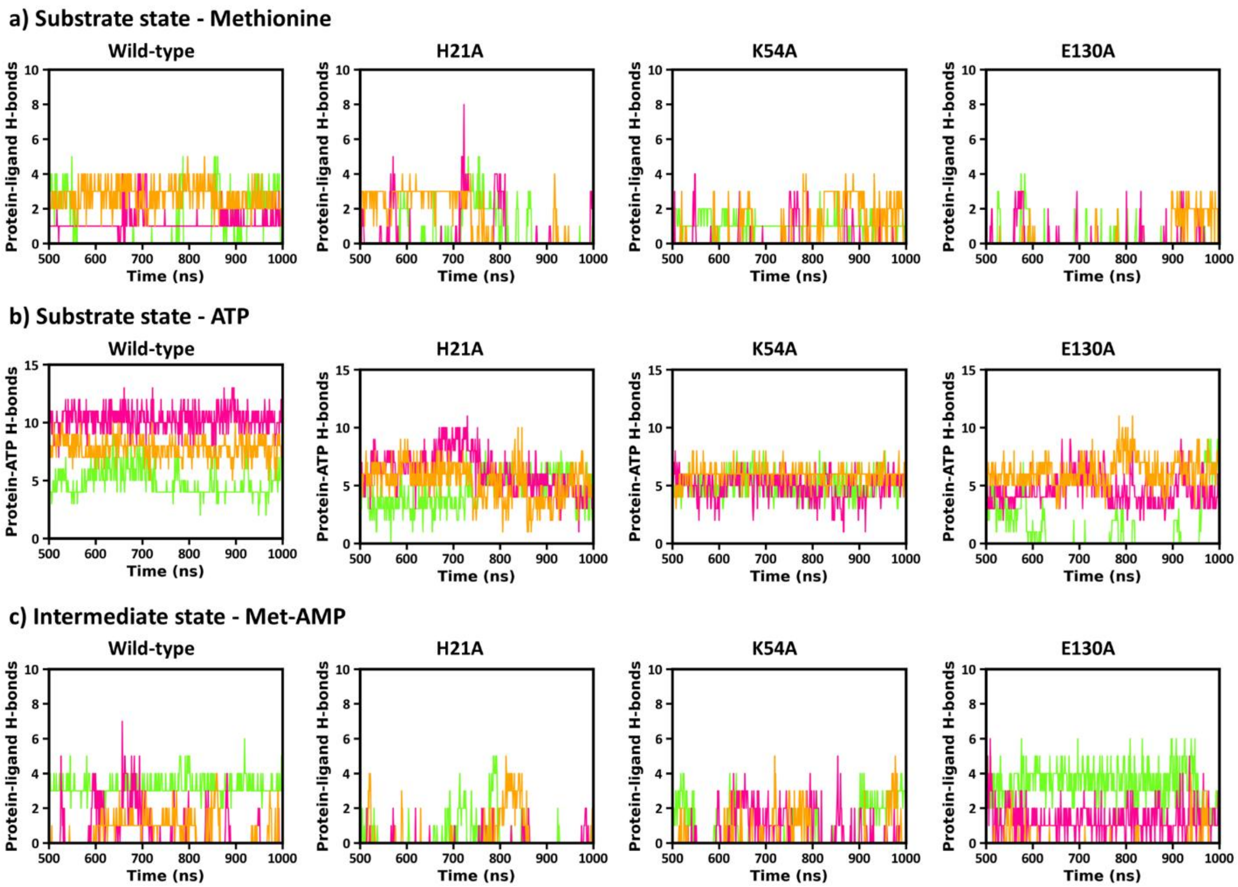


**Figure S9. Protein-ligand Hydrogen bond interactions. (a-b)** Substrate state - methionine and ATP, **(c)** intermediate state - Met-AMP for wild-type and three mutants.

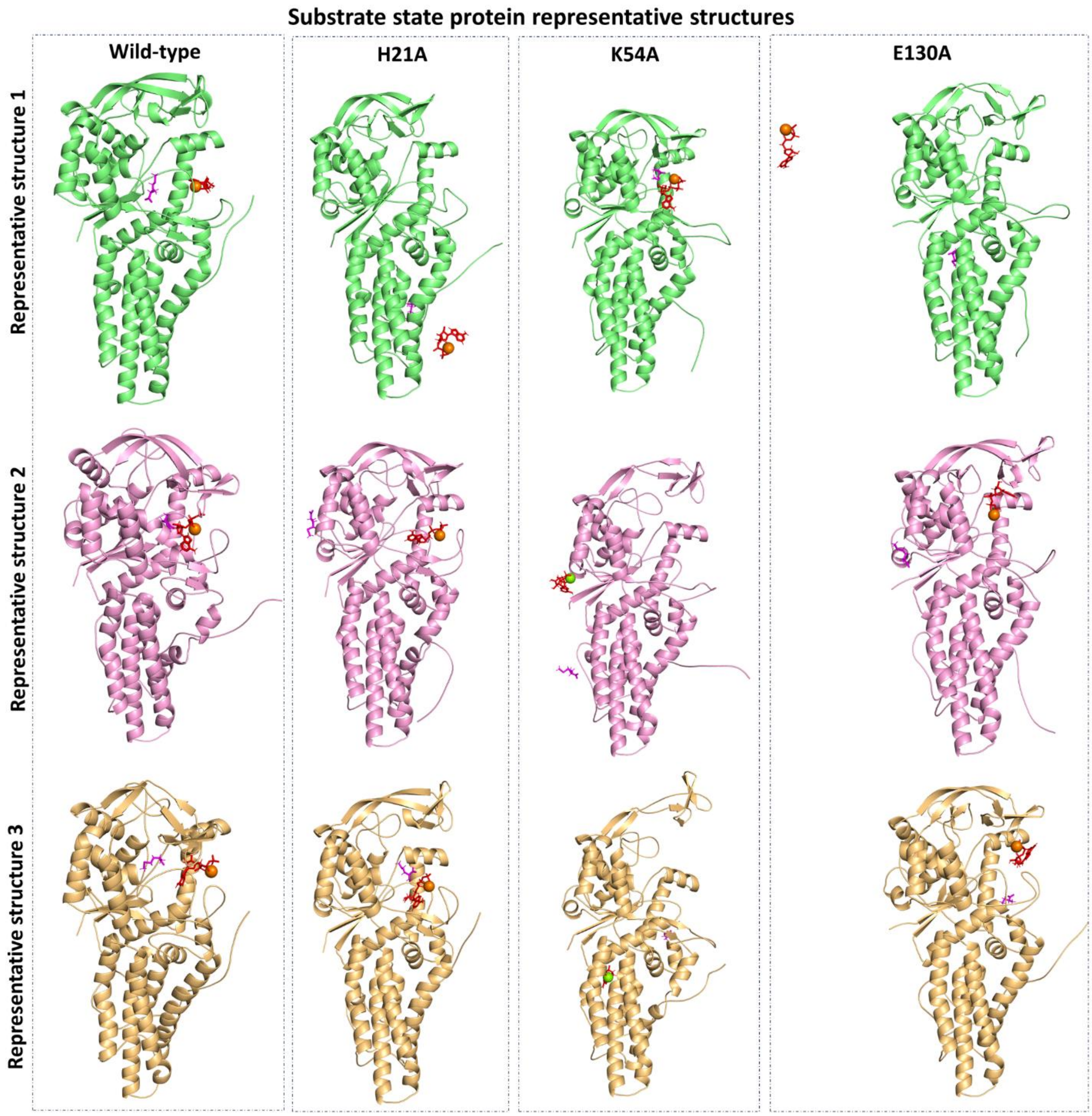


**Figure S10.** Representative structures for substrate state MetRS protein over three simulations.

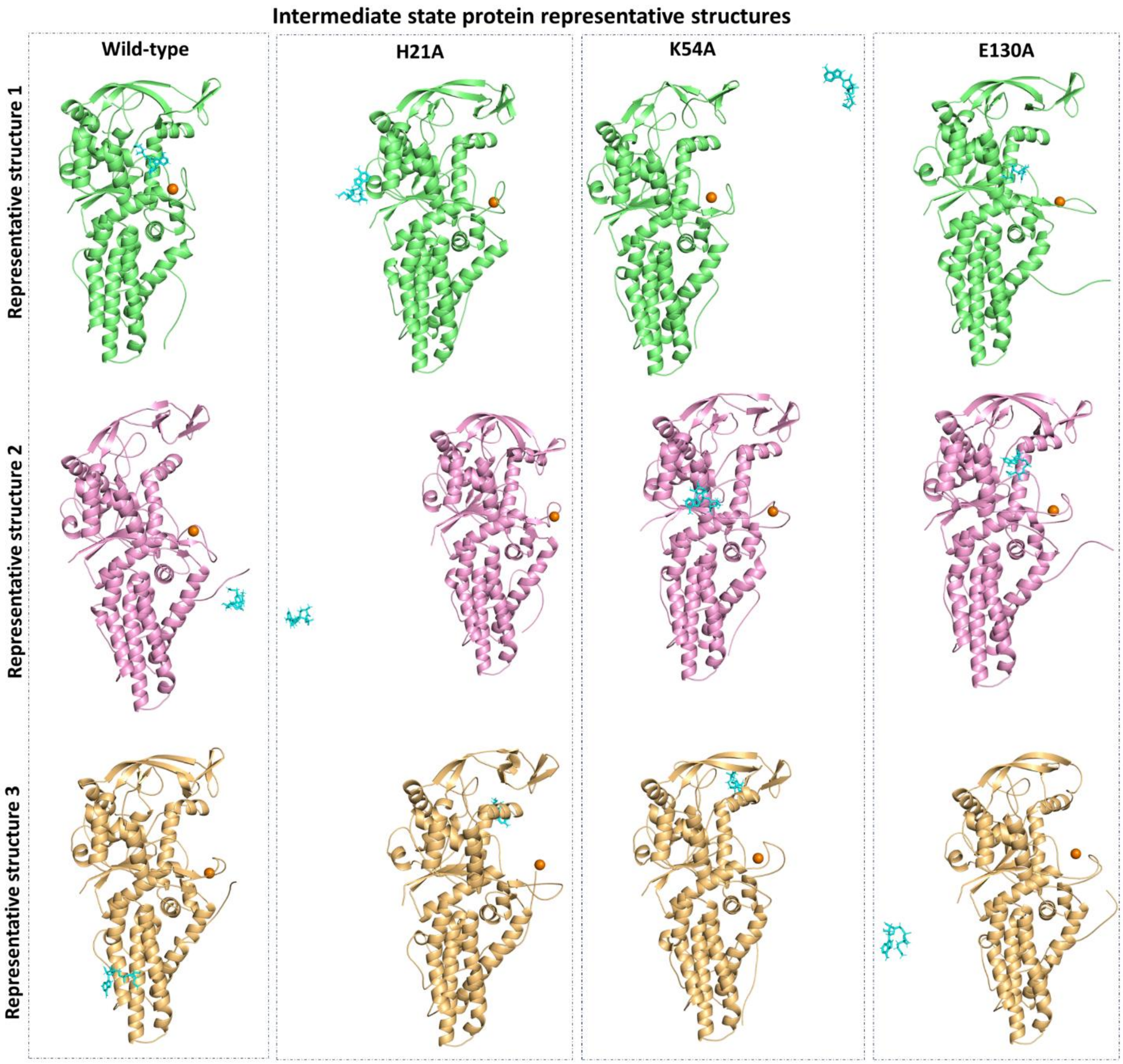


**Figure S11.** Representative structures for intermediate state MetRS protein over three simulations.

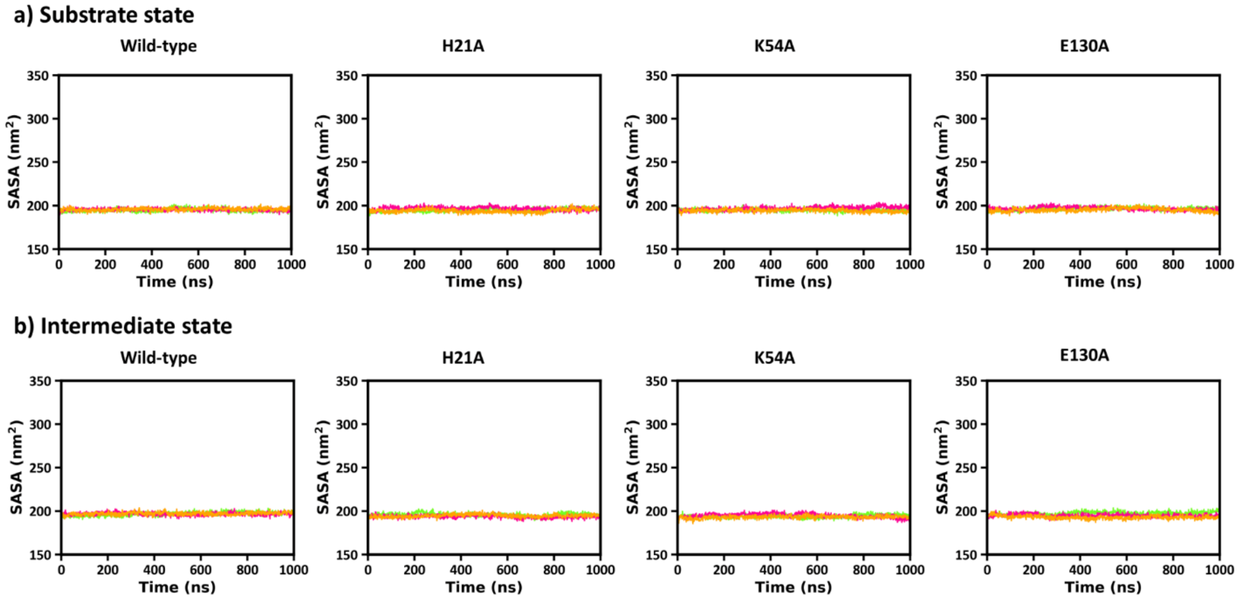


**Figure S12. Hydrophobic SASA of protein.** Hydrophobic SASA values for **(a)** substrate state and **(b)** intermediate state in wild-type and mutant models (H21A, K54A and E130A).

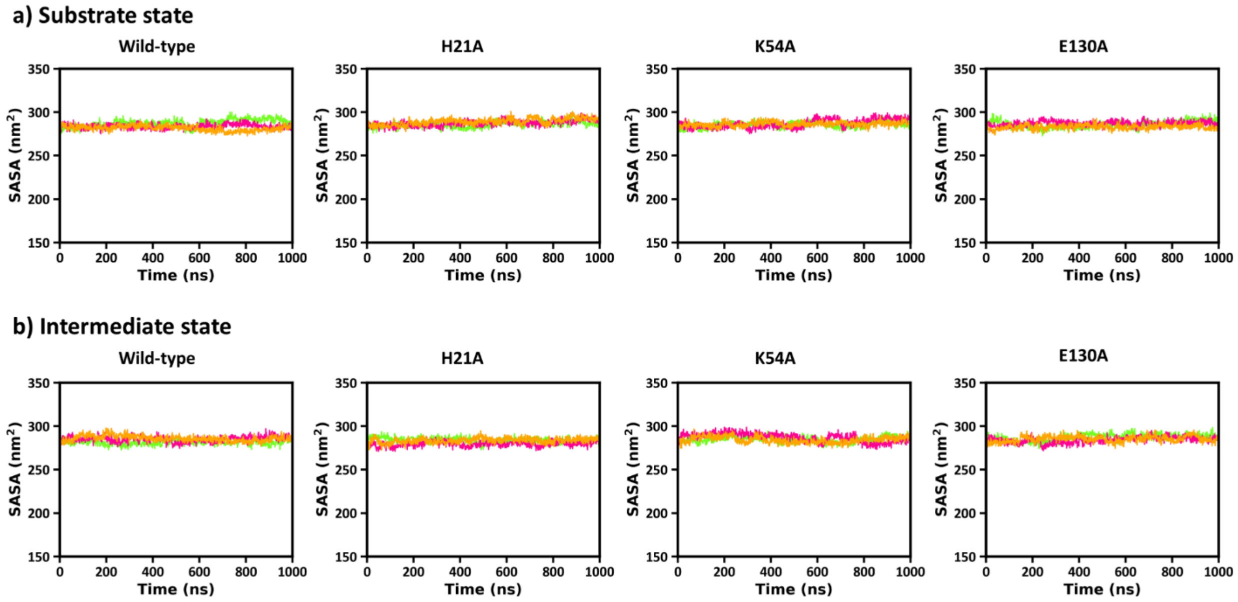


**Figure S13. Hydrophilic SASA of protein.** Hydrophilic SASA values for **(a)** substrate state and **(b)** intermediate state in wild-type and mutant models (H21A, K54A and E130A).

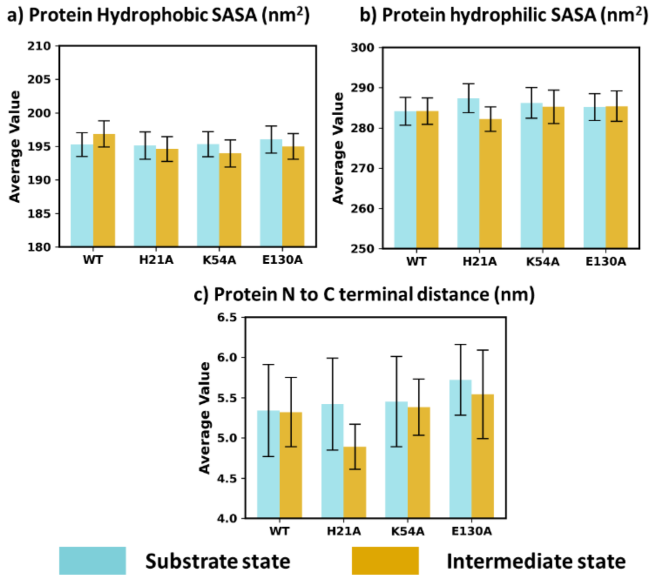


**Figure S14. Average and standard deviation (error bar) of protein properties. (a)** Hydrophobic SASA, **(b)** hydrophilic SASA, **(c)** protein RMSF and **(d)** N to C terminal distance of protein.

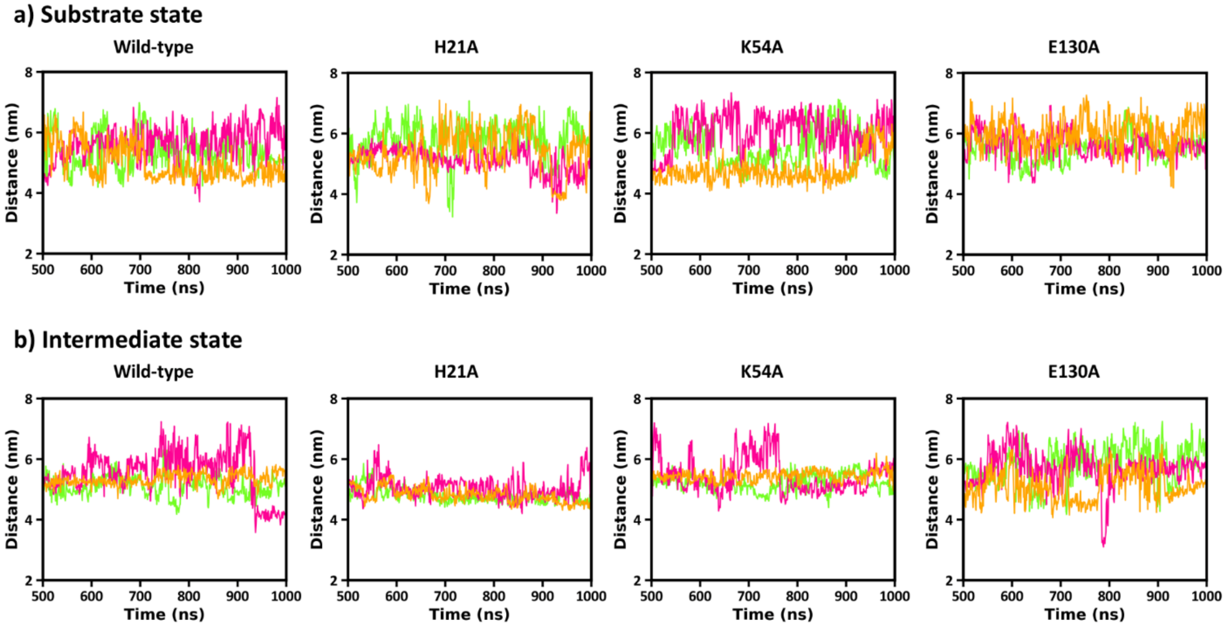


**Figure S15. Variation in N to C terminal distance of protein over time. (a)** Substrate state and **(b)** intermediate state for wild-type and different mutant models.

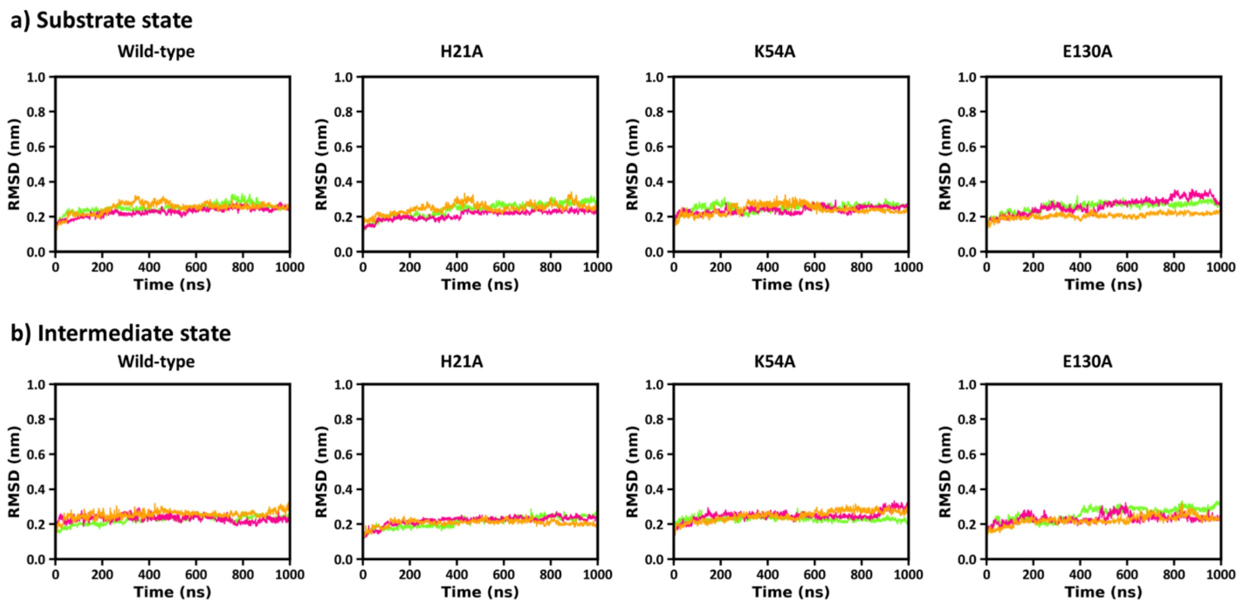


**Figure S16. RMSD values for the catalytic domain residues. (a)** Substrate and **(b)** intermediate state.

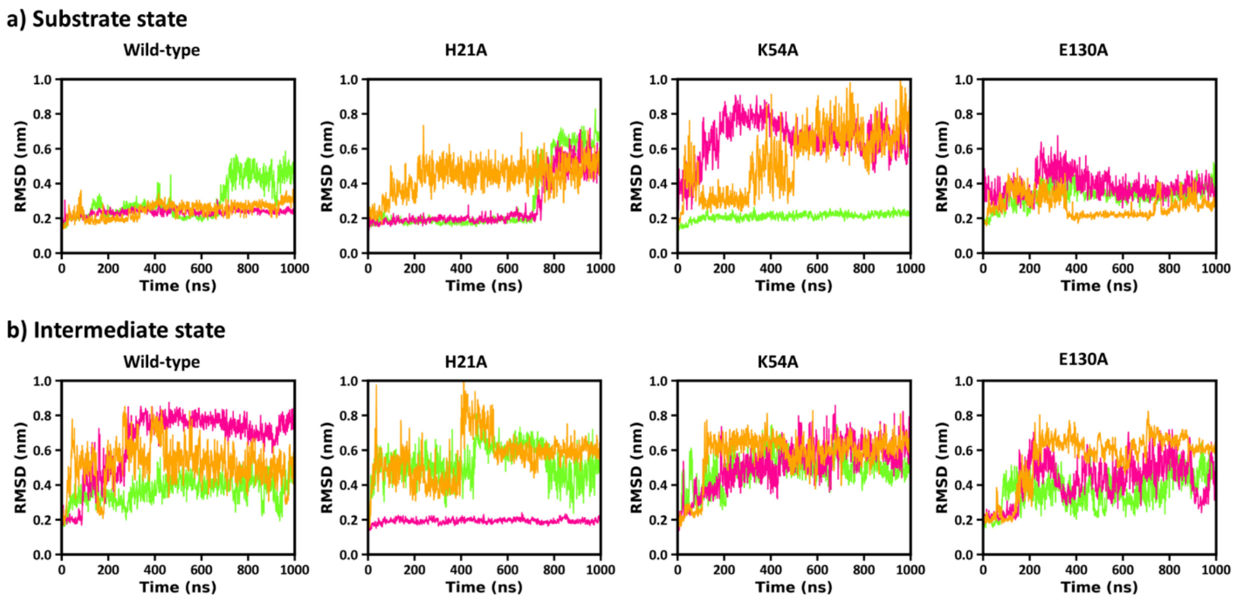


**Figure S17. RMSD values for the connective peptide (CP) domain residues. (a)** Substrate and **(b)** intermediate state.

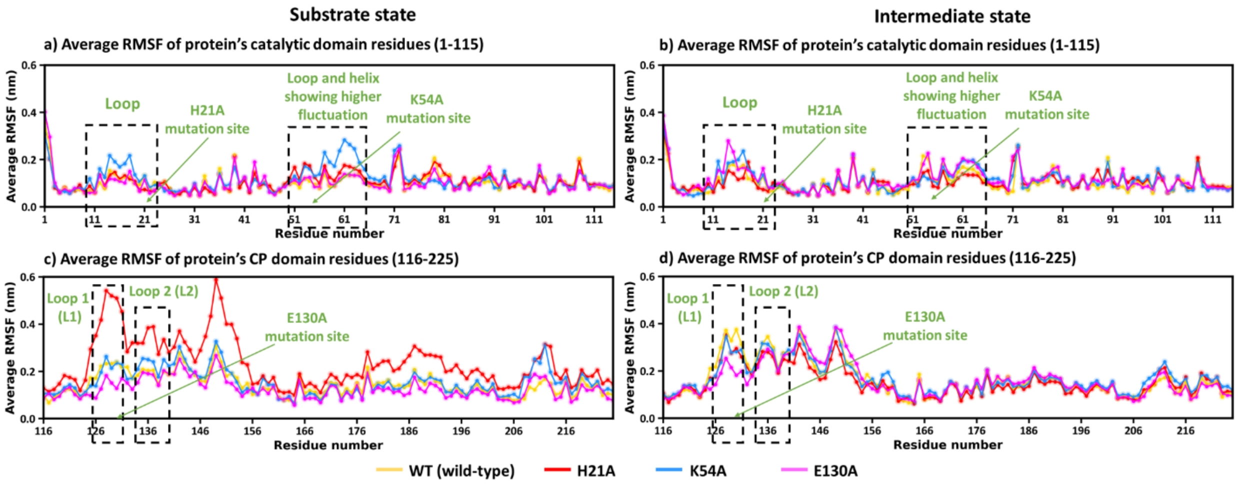


**Figure S18. Residue-wise average RMSF (over three simulations) of catalytic domain (1-115) and CP domain (116-225) residues in wild-type (yellow), H21A (red), K54A (blue), E130A (magenta) protein variants.** Average RMSF value of catalytic domain residues in **(a)** substrate state and **(b)** intermediate state. Average RMSF value of connective peptide domain residues in **(c)** substrate state and **(d)** intermediate state.

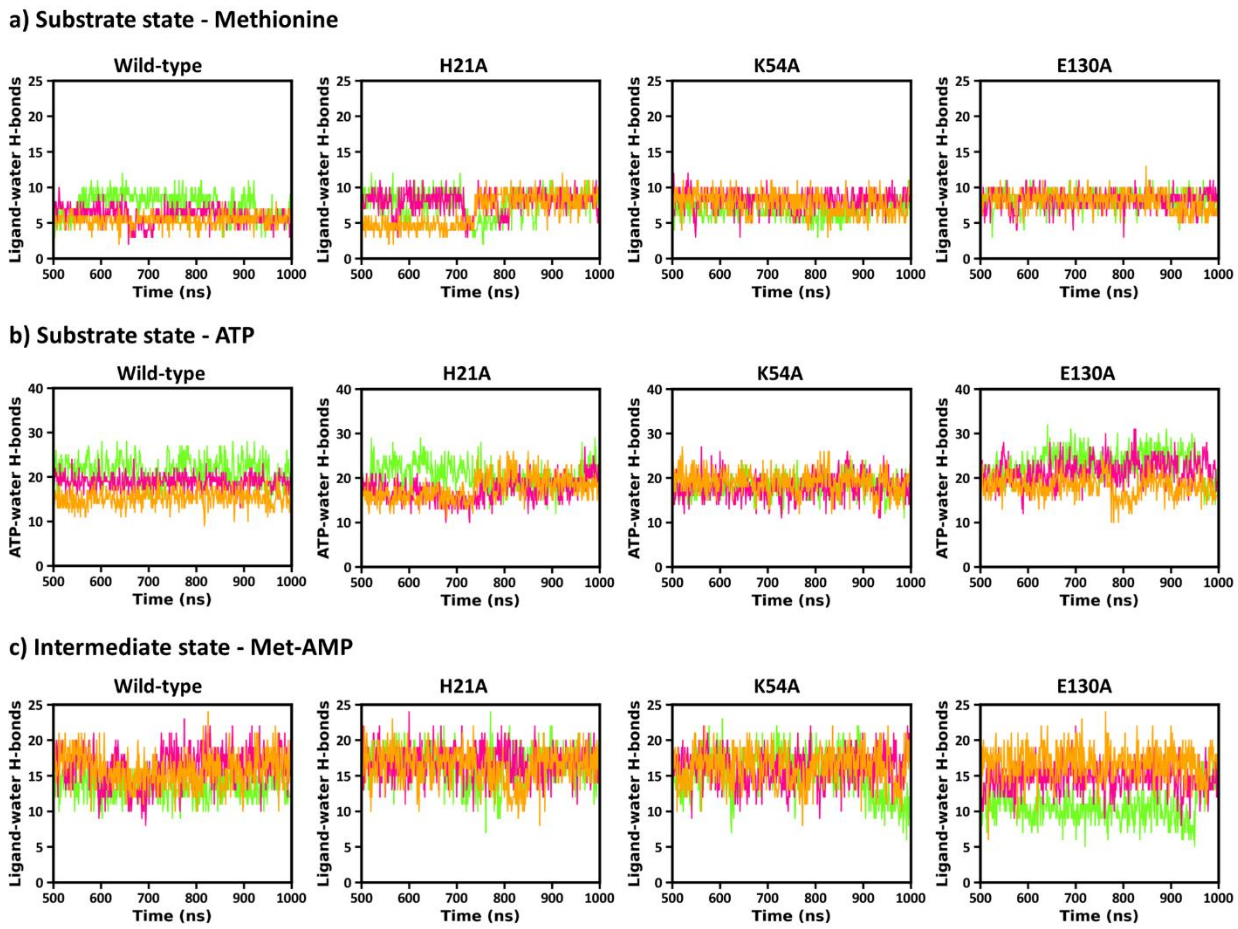


**Figure S19. Ligand-water hydrogen bond interactions (Substrate state - Methionine and ATP, Intermediate state - Met-AMP). (a-b)** Substrate state and **(c)** intermediate state.

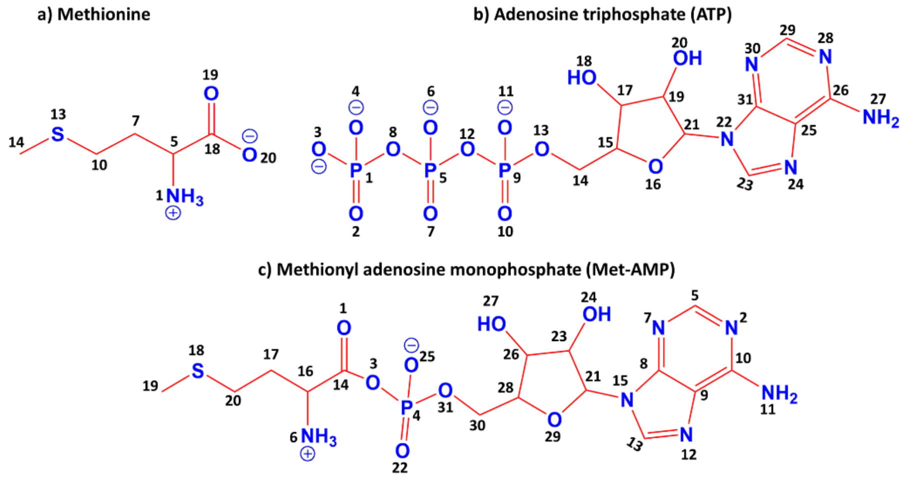


**Figure S20. The heavy atoms of (a) methionine, (b) ATP and (c) Met-AMP are numbered for RMSF calculations.**

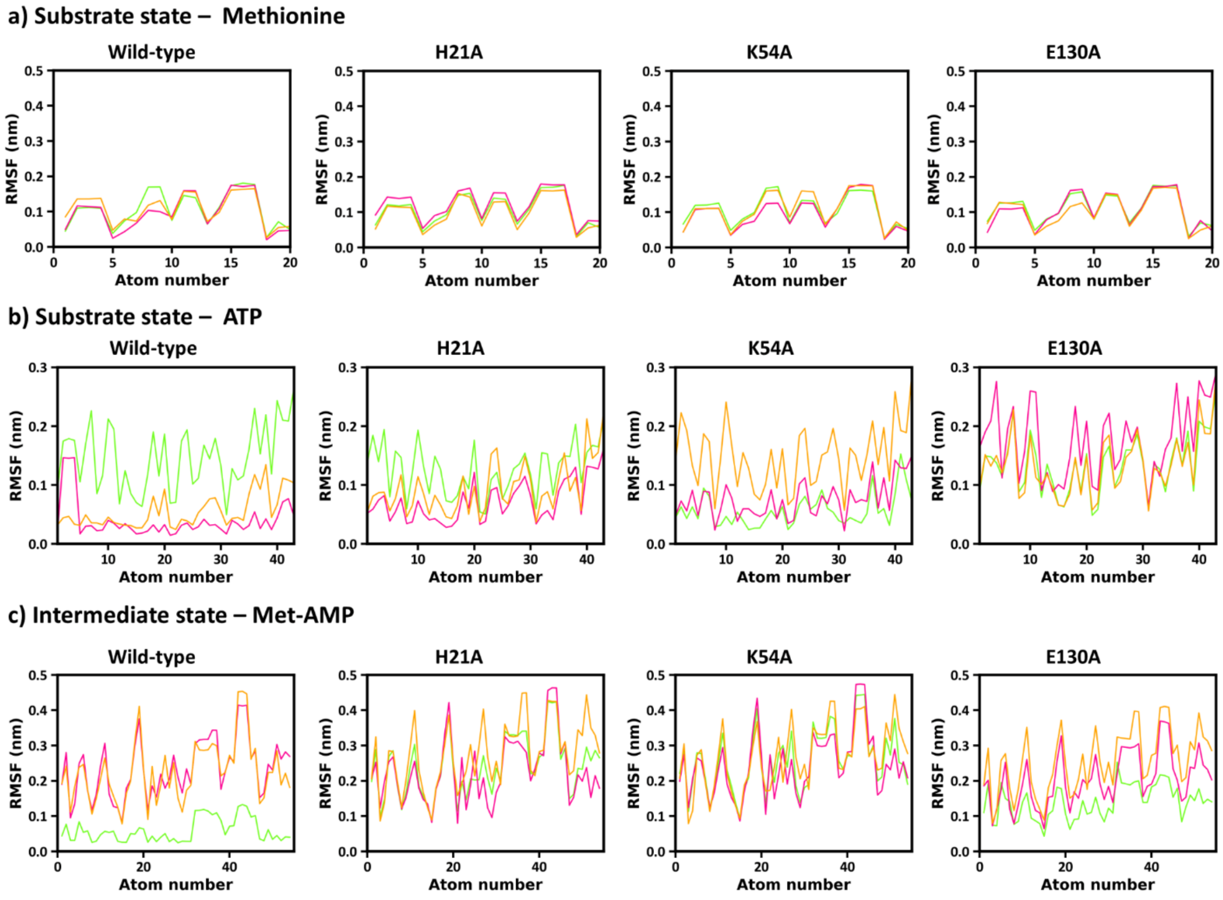


**Figure S21. RMSF values for ligand's heavy atoms (Substrate state - Methionine and ATP, Intermediate state - Met-AMP). (a-b)** Substrate state and **(c)** intermediate state.

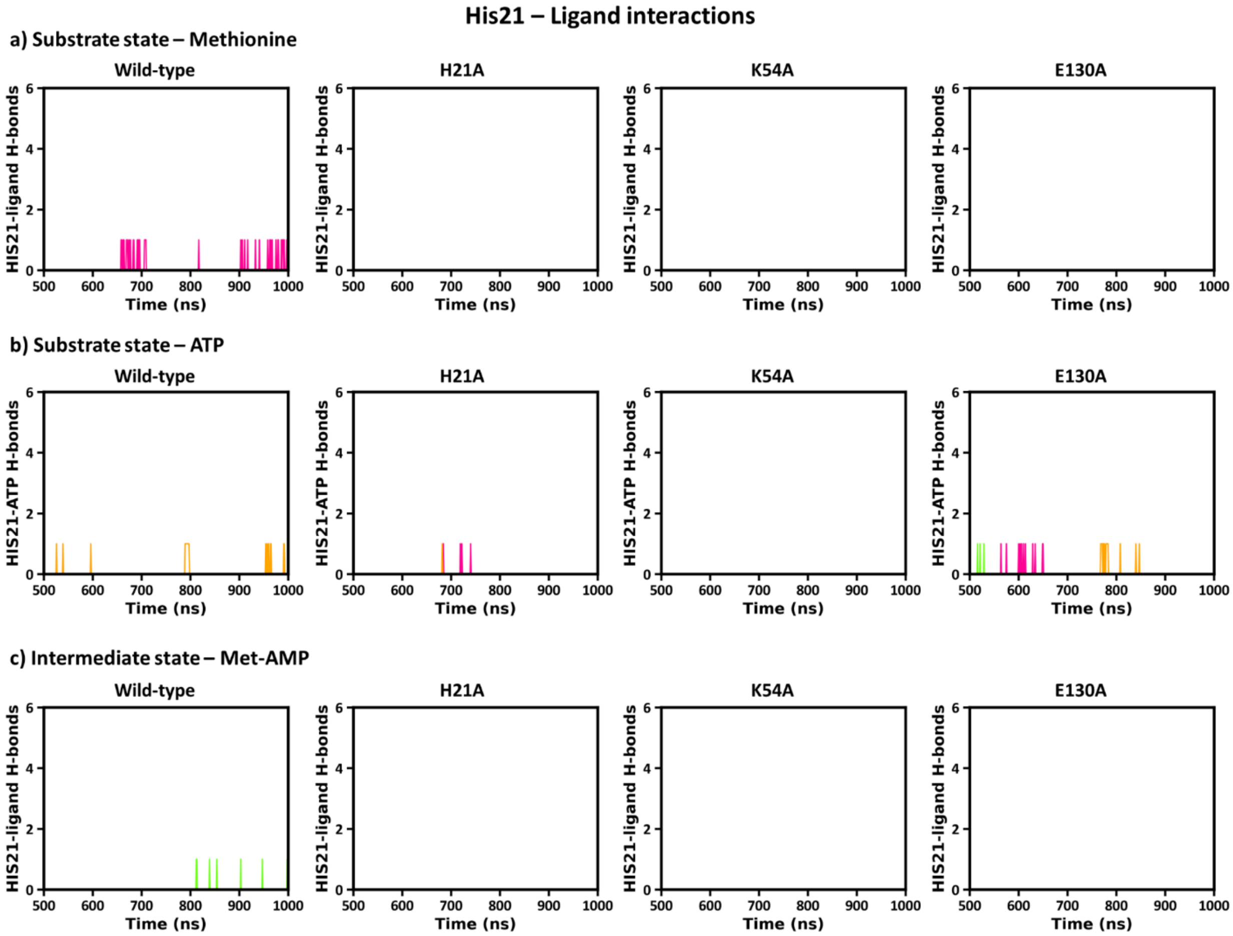


**Figure S22. Hydrogen bond interaction counts of His21 residue with ligand. (a-b)** Substrate state - Methionine and ATP, **(b)** Intermediate state - Met-AMP.

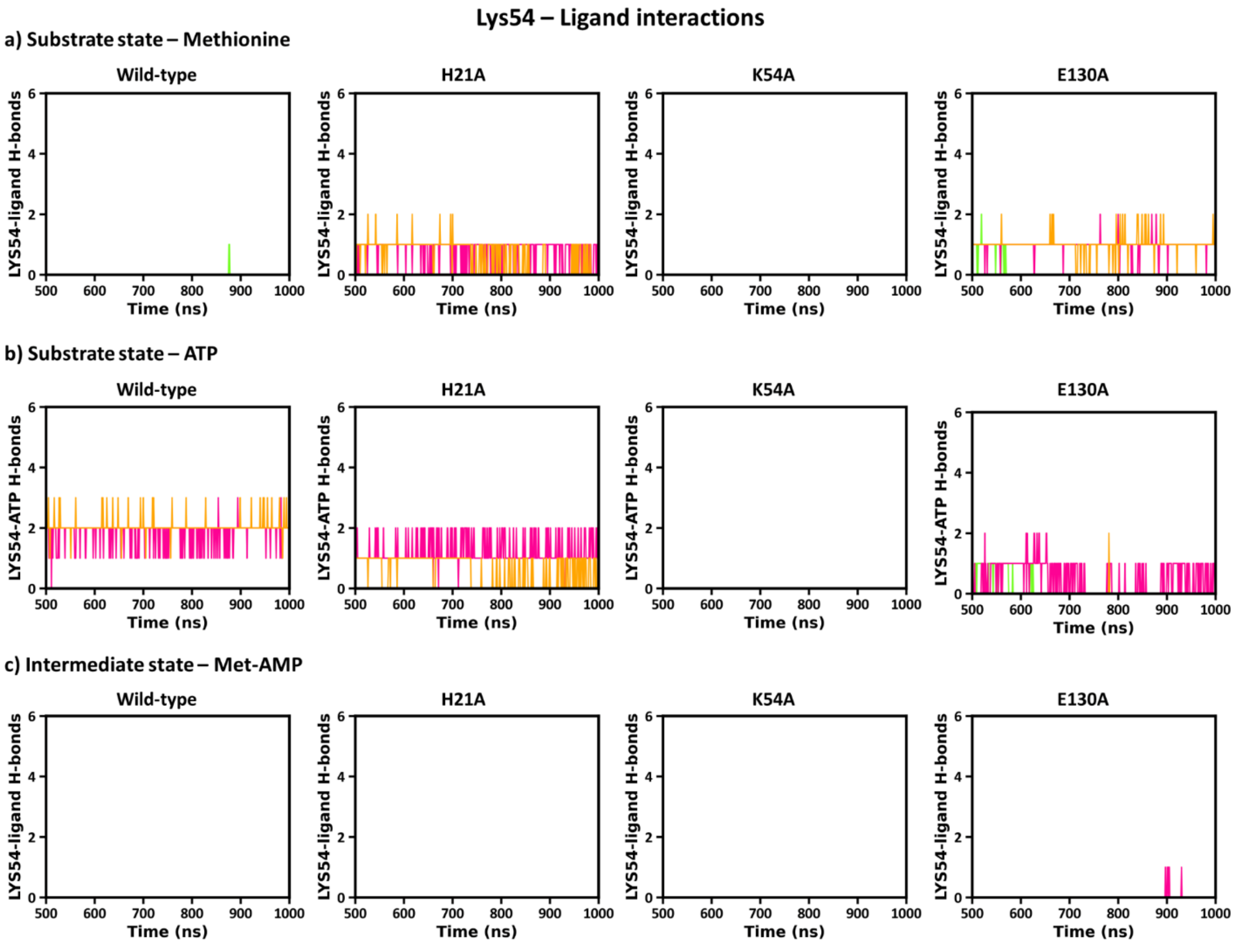


**Figure S23. Hydrogen bond interaction counts of Lys54 residue with ligand. (a-b)** Substrate state - Methionine and ATP, **(b)** Intermediate state - Met-AMP.

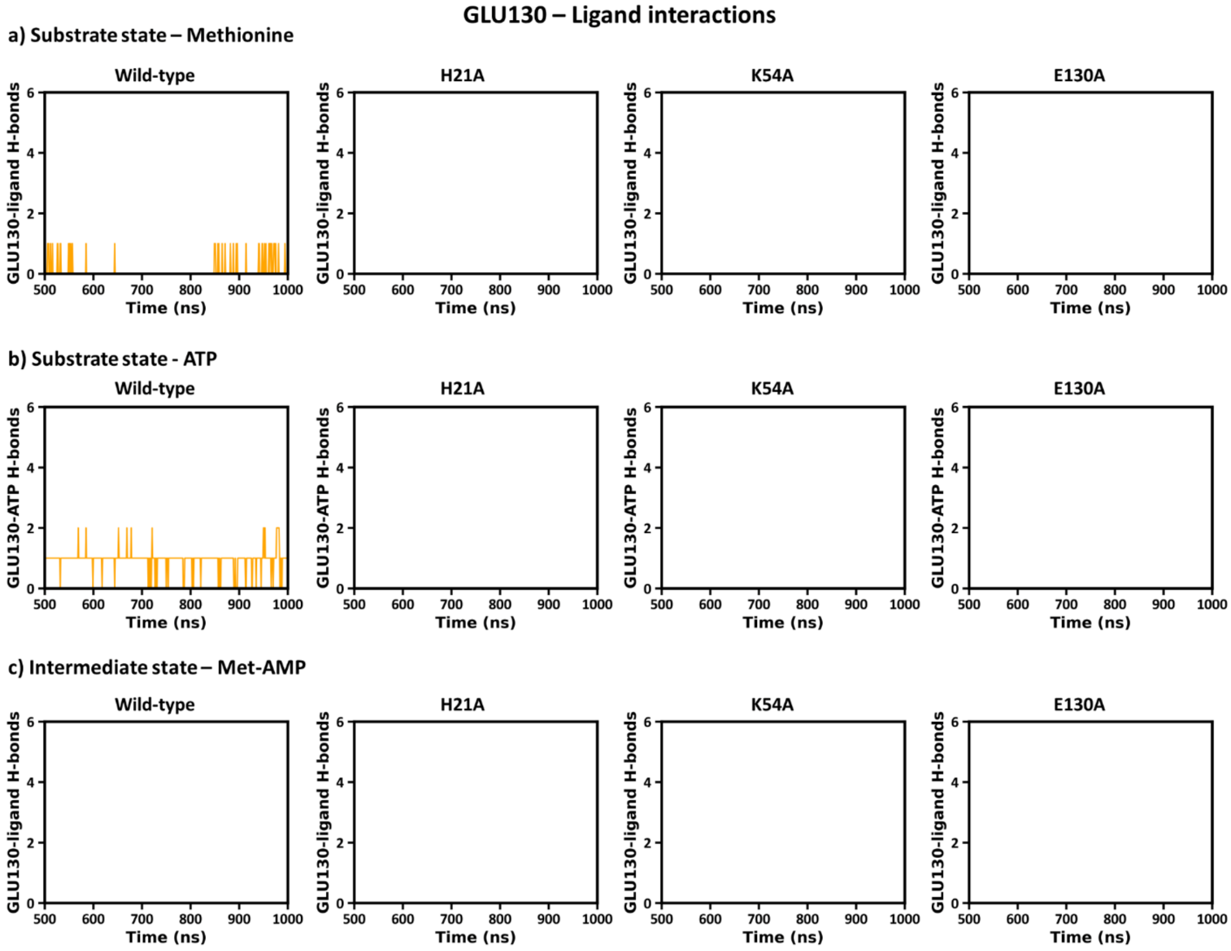


**Figure S24. Hydrogen bond interaction counts of Glu130 residue with ligand. (a-b)** Substrate state - Methionine and ATP, **(b)** Intermediate state - Met-AMP.

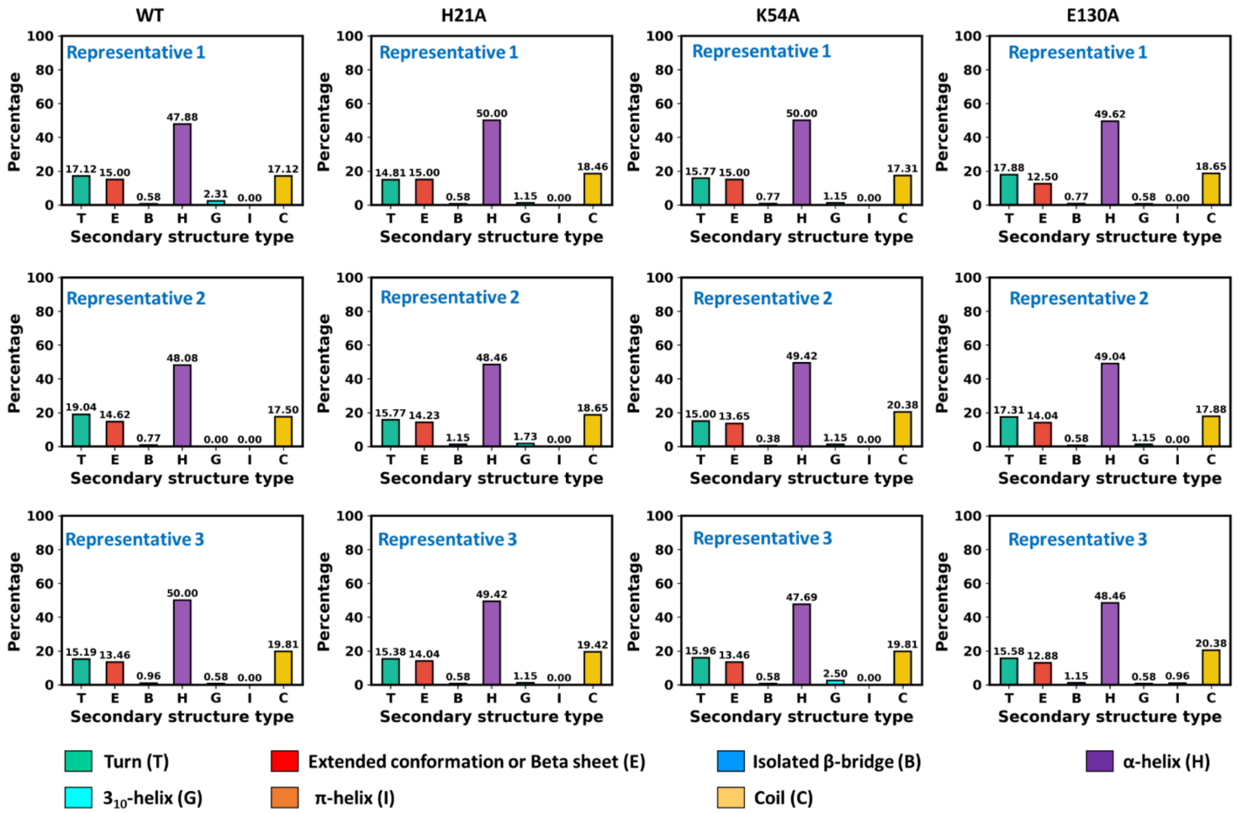


**Figure S25. Protein's secondary structure for substrate state representative structures**. Representative 1, Representative 2, Representative 3 obtained from each simulation.

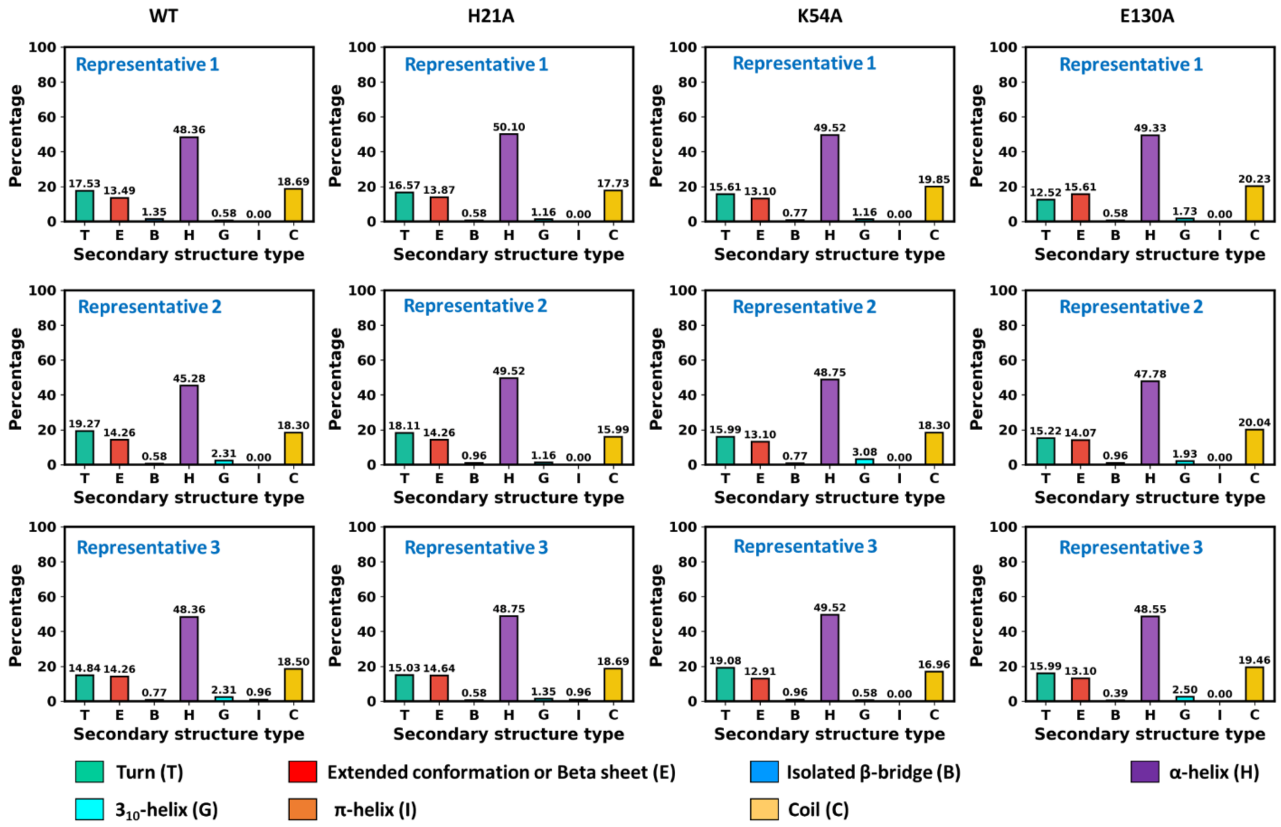


**Figure S26. Protein's secondary structure for intermediate state representative structures**. Representative 1, Representative 2, Representative 3 obtained from each simulation.

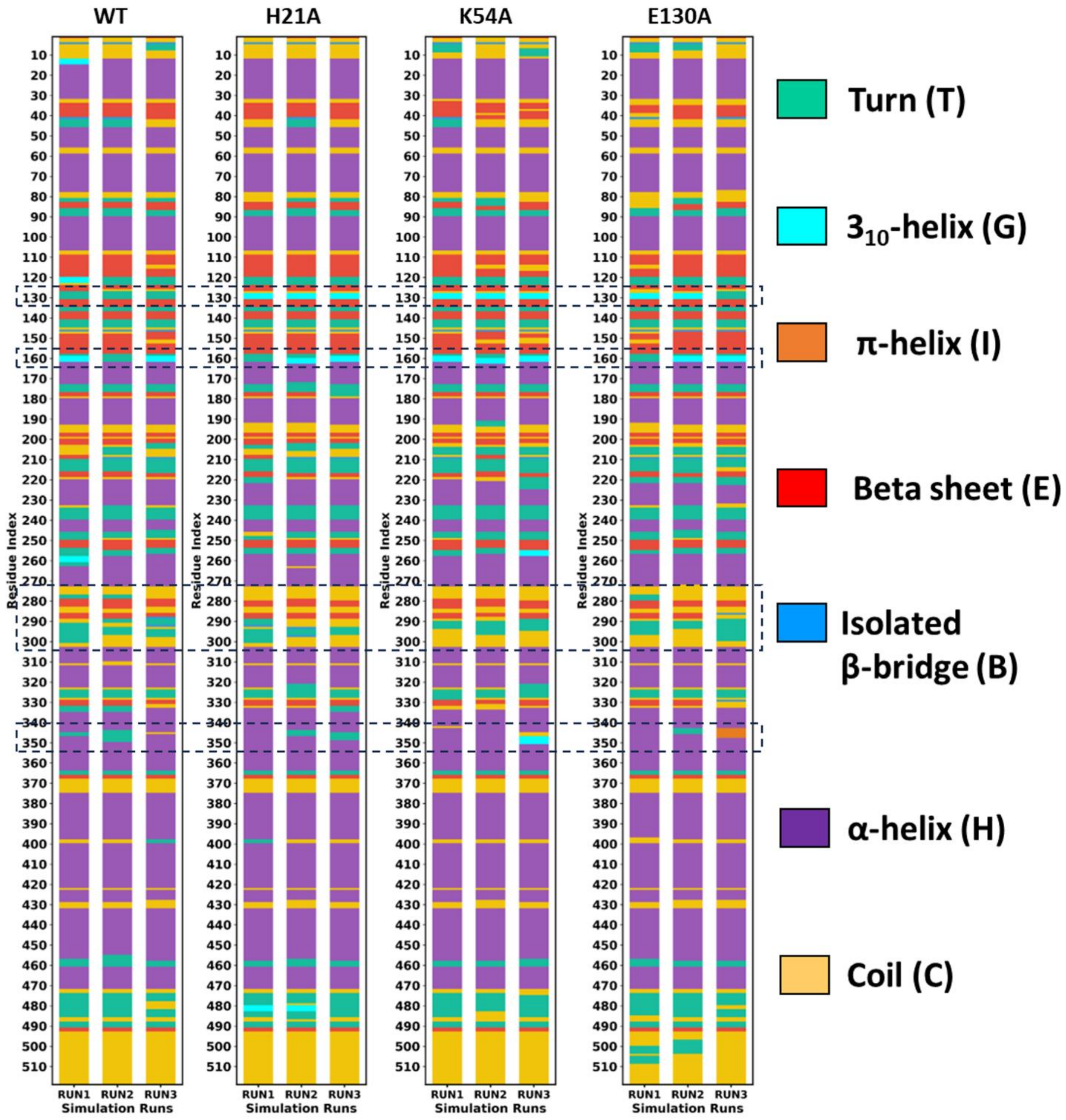


**Figure S27. Residue-wise secondary structure of substrate state representative structure protein in each simulation (RUN1, RUN2, RUN3) of WT, H21A, K54A and E130A model.**

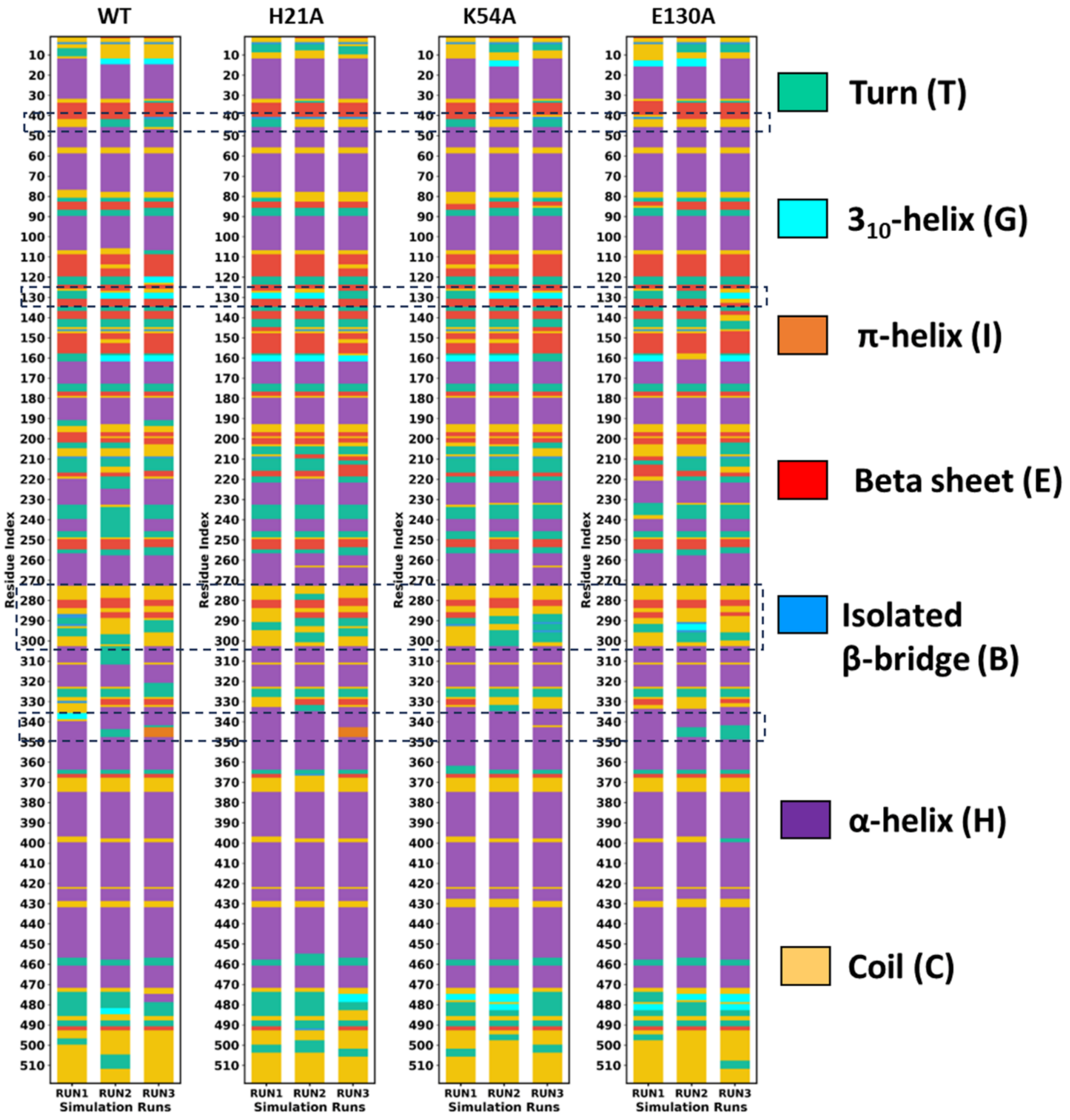


**Figure S28. Residue-wise secondary structure of intermediate state representative structure protein in each simulation (RUN1, RUN2, RUN3) of WT, H21A, K54A and E130A model.**

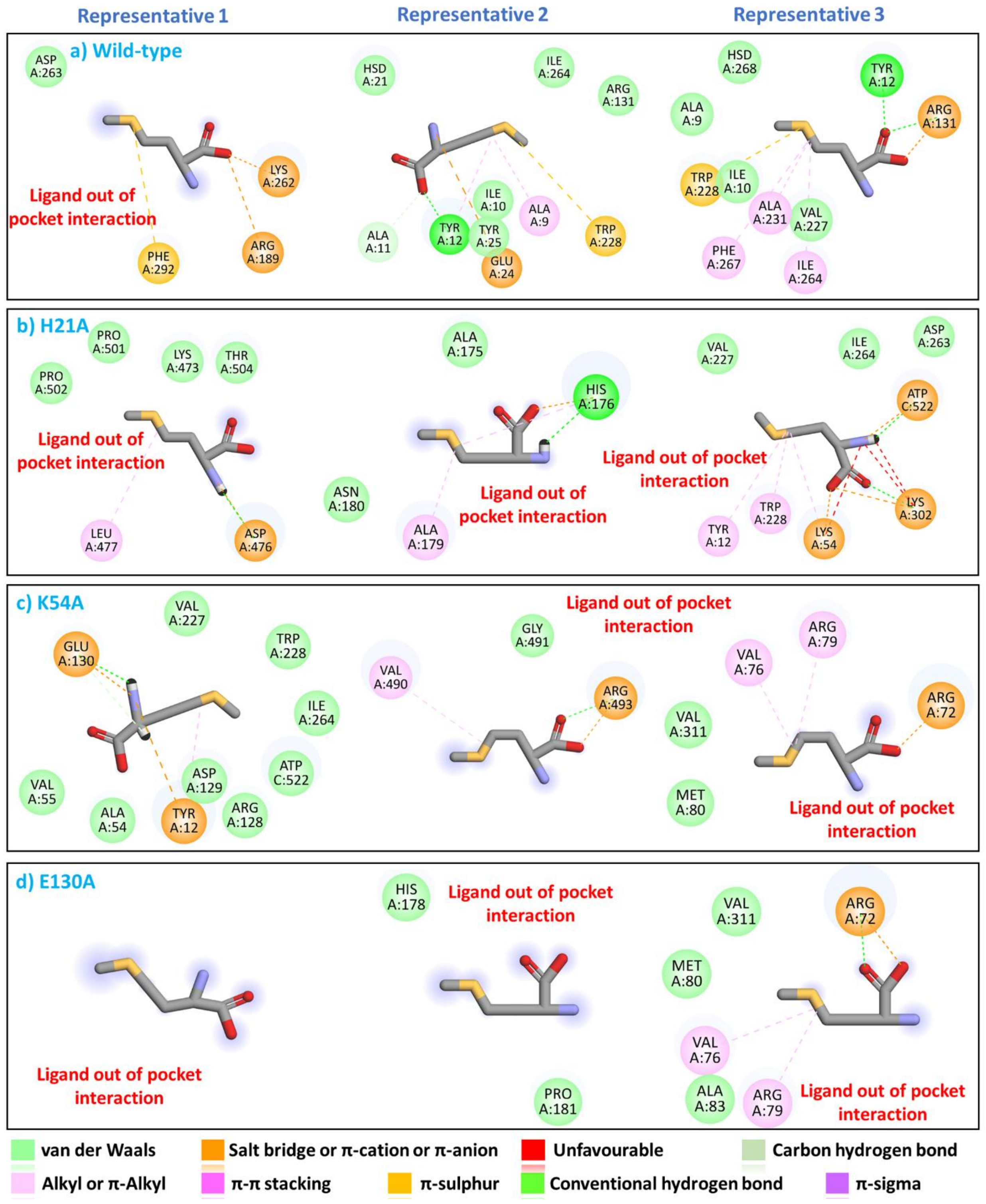


**Figure S29. Interactions formed by methionine with MetRS protein in representative structures.** Interactions are shown for each simulation of wild-type, H21A, K54A & E130A.

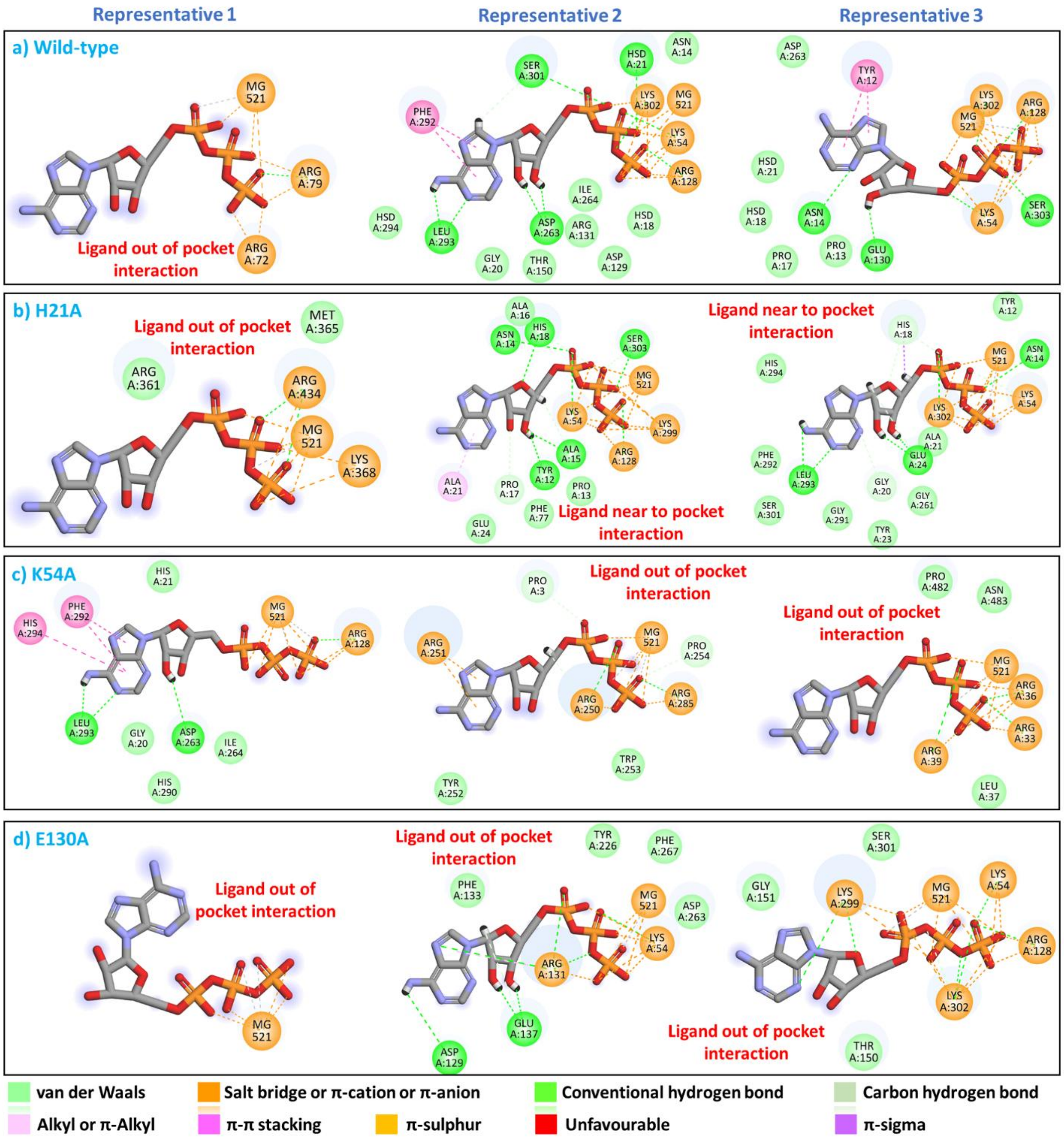


**Figure S30. Interactions formed by ATP with MetRS protein in representative structures.** Interactions are shown for each simulation of wild-type, H21A, K54A & E130A.

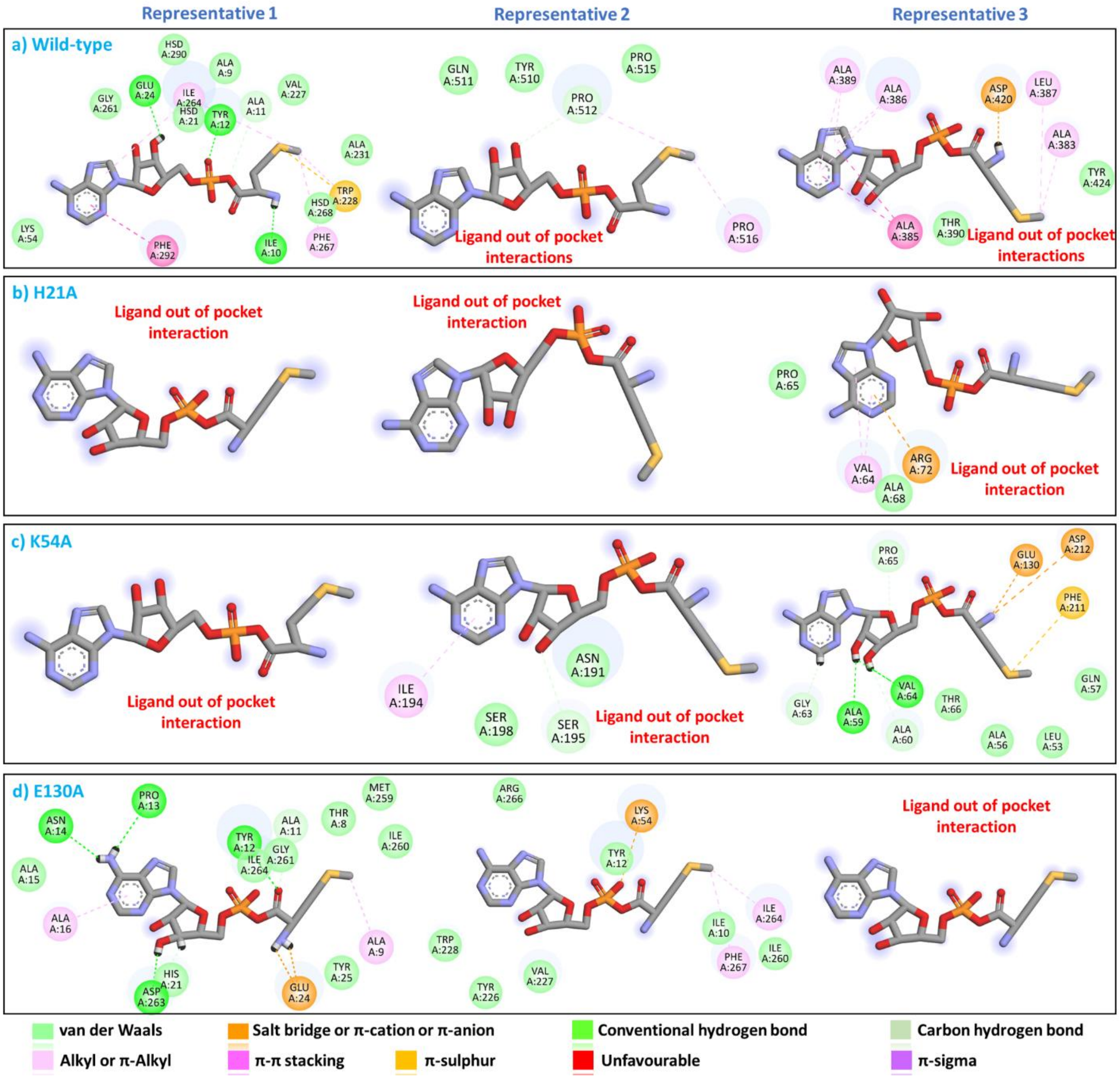


**Figure S31. Interactions formed by Met-AMP with MetRS protein in representative structures.** Interactions are shown for each simulation of wild-type, H21A, K54A & E130A.
